# Inter-individual gene expression variability implies stable regulation of brain-biased genes across organs in three ray-finned fishes

**DOI:** 10.1101/2024.11.11.623020

**Authors:** Christabel F. Bucao, Consolée Aletti, Alexandre Laverré, Sébastien Moretti, Alexandra Trouvé, Andrew W. Thompson, Brett L. Racicot, Catherine A. Wilson, Julien Bobe, Ingo Braasch, Yann Guiguen, John H. Postlethwait, Marc Robinson-Rechavi

## Abstract

Phenotypic variation among individuals provides the raw material for evolution, and gene expression is a key mediator between genetic and phenotypic variation. As for phenotypes, the range of gene expression is limited and biased by evolutionary and developmental constraints. Observed expression variability due to biomolecular stochasticity and cell-to-cell heterogeneity has been well-studied in isogenic populations of unicellular organisms. However, for multicellular organisms with a diversity of cells and tissues sharing the same genetic background, the interplay between expression variability, gene and organ function, and gene regulation remains an open question. Here, we use highly multiplexed 3’-end bulk RNA sequencing to generate transcriptome profiles spanning at least nine organs in outbred individuals of three ray-finned fish species: zebrafish, Northern pike, and spotted gar. Per organ, we quantify individual-to-individual gene expression variability independent of mean expression level. Lowly variable genes are enriched in cellular housekeeping functions whereas highly variable genes are enriched in stimulus-response functions. Furthermore, highly variable genes evolve under weaker purifying selection at the protein-coding sequence, indicating that intra-species expression variability predicts inter-species protein sequence divergence. Genes that are broadly expressed across organs are both highly expressed and lowly variable, whereas organ-biased genes are typically highly variable within their top organ. Among organ-biased genes, patterns of expression variance are dependent on the top organ. Specifically, brain- and gonads-biased genes have lowly variable expression across different organs, suggesting stabilizing selection. These patterns suggest that gene regulatory mechanisms evolve under organ-specific selective pressures.

## Introduction

Phenotypic variation among individuals is crucial for evolutionary processes, as it serves as the substrate for natural selection. Gene expression is a key mediator of this phenotypic diversity. Similar to how phenotypic variation is shaped and constrained by evolutionary and developmental factors, variation in gene expression among individuals is also subject to comparable limitations and biases, with some genes having a wider expression distribution than others. This gene expression variance is influenced by various ‘intrinsic’ (gene-specific) and ‘extrinsic’ factors spanning biological scales, including transcriptional or translational stochasticity (Kærn et al. 2005), cellular heterogeneity (Carter and Zhao 2021), genetic variation (Hill et al. 2020), epigenetic modifications (Choi and Kim 2008; Carter and Zhao 2021), as well as micro- and macro-environmental cues (López-Maury et al. 2008). The observed expression variance (or dispersion) is often used as a proxy measure for expression *variability*, broadly defined as the *propensity* of a gene to vary its expression level in response to stochastic, genetic, or environmental perturbations (though see (Wagner and Altenberg 1996)).

Focusing on transcriptional or translational stochasticity (‘expression noise’) in genetically identical cells, many studies on isogenic populations of unicellular organisms (e.g., *Escherichia coli* (Elowitz et al. 2002; Silander et al. 2012; Jones et al. 2014; Wolf et al. 2015; Urchueguía et al. 2021) and *Saccharomyces* (Fraser et al. 2004; Newman et al. 2006; Zhang et al. 2009; Lehner 2008, 2010; Bódi et al. 2017; Duveau et al. 2018)) have probed the relationship between expression noise and gene promoter features (Silander et al. 2012; Jones et al. 2014; Wolf et al. 2015; Urchueguía et al. 2021), as well as properties of the gene regulatory network (Kittisopikul and Süel 2010; Chalancon et al. 2012; Munsky et al. 2012). Evidence suggests that natural selection can shape expression noise according to gene function (Silander et al. 2012; Fraser et al. 2004; Newman et al. 2006; Lehner 2008; Richard and Yvert 2014) and that constraints on expression noise are mediated by environmental context (LaBoone and Assis 2024). Furthermore, a correlation between gene expression noise in isogenic yeast cells and broader expression variation in non-isogenic populations has been reported (Dong et al. 2011), suggesting that some shared regulatory mechanisms underlie both intra-genotype expression noise and inter-genotype expression variability (Sigalova et al. 2020). The extent to which these findings in unicellular organisms can be extrapolated to complex multicellular organisms, which have functionally specialized organs with distinct expression profiles despite a common genetic background, is not well understood. While many studies have advanced our understanding of genetic variants underlying variation in gene expression levels within a population (e.g., eQTL analysis (Dimas et al. 2009; Nica and Dermitzakis 2013)), much remains to be known about the relationship among expression variability, gene function, and gene regulation in animals. Although organ-specific enhancers exist to facilitate spatiotemporal control of gene expression (Schoenfelder and Fraser 2019), we expect some level of pleiotropy of regulation of gene expression in different organs. Thus selection on gene expression in different organs is not totally independent, and the expression variability of a gene within any given organ may potentially be constrained by antagonistic organ-specific expression optima.

Studies on gene expression variability in animals have primarily focused on select cell types or organs in model organisms (Faure et al. 2017; Barroso et al. 2018; Foreman and Wollman 2020), with particular interests in the role of stochasticity in cell differentiation (Simon et al. 2018) (including shifts in inter-cell (Ohnishi et al. 2014; Richard et al. 2016) or inter-embryo expression variability over development (Liu et al. 2020)), as well as expression variability in human organs and its association with disease (Ho et al. 2008; Li et al. 2010; Mar et al. 2011; Dinalankara and Bravo 2015; Simonovsky et al. 2019; Bashkeel et al. 2019; Alemu et al. 2014; Wolf et al. 2023). Of note, although most studies compare expression in different organs, the terminology of ‘tissues’ and ‘tissue-specificity’ is frequently used in the literature instead. While the functional relevance of expression variability is increasingly recognized (Geiler-Samerotte et al. 2013; De Jong et al. 2019; Mar 2019), how such variability is modulated by organ context remains largely unexplored, even in studies that sample multiple organs (Sigalova et al. 2020; Simonovsky et al. 2019; Alemu et al. 2014; Wolf et al. 2023). Few studies have been published on variability in outbred, non-model animal populations (but see (Horta-Lacueva et al. 2023; Hagai et al. 2018; Fair et al. 2020)). This is partly due to heavy resource requirements such as the need for many replicates for statistical significance, as well as the challenges associated with disentangling different sources of expression variation (McCall et al. 2016), whether in bulk or at the single-cell level.

Recently, the advent of low-cost high-throughput sequencing methods has made it increasingly feasible to generate sufficient transcriptome data to measure within-organ gene expression variability across several individuals. In this study, we present an analysis of inter-individual gene expression variability in multiple organs, using newly generated 3′-end Bulk RNA Barcoding and sequencing (BRB-seq) (Alpern et al. 2019) data in three ray-finned fish species. Characterizing gene expression variance within and between organs provides a a new framework to understand the interplay between gene function and organ-dependent constraints on gene regulatory evolution in complex multicellular organisms.

## Results

### Organ transcriptome profiles of ray-finned fishes

To investigate inter-individual gene expression variability within and between organs, we generated transcriptomic data across multiple organs using 3′-end Bulk RNA Barcoding and sequencing (BRB-seq) (Alpern et al. 2019) for three species of ray-finned fishes (Actinopterygii): two teleost species (*Danio rerio* (zebrafish) and *Esox lucius* (Northern pike)) and one non-teleost outgroup (*Lepisosteus oculatus* (spotted gar)). Zebrafish and Northern pike represent two of the major teleost clades (Otomorpha and Euteleosteomorpha, respectively) and neither species have undergone additional lineage-specific genome duplications after the teleost whole-genome duplication (Jaillon et al. 2004), while the spotted gar is a slowly evolving outgroup to the teleosts (Braasch et al. 2016). For each species, we sampled multiple outbred individuals spanning several organs in both sexes (when known) (**Figure 1A**) and applied a two-step filter to remove low quality samples based on sequencing quality (**Figures S1-S3**) and within-organ expression correlation (Materials and Methods). We observed that this stringent two-step filtering protocol is sufficient to remove any strong batch effects in our dataset (**Figures S4**-**S6**). Hierarchical clustering of zebrafish samples based on pairwise Pearson’s correlation shows that transcriptome profiles cluster into five major groups by related organ systems (**Figure 1B**; see **Figures S7-S9** for PCA plots). Reproductive organs (cluster 1) appear to have the most divergent expression profiles, while the following clusters can be observed for non-reproductive organs: gastrointestinal and urinary systems (kidney, intestine, and liver; cluster 2); nervous system (brain and eye; cluster 3); external organs (pectoral fin, skin, and gills; cluster 4); and lastly, muscular system (heart and muscle; cluster 5). Apart from the gonads, we do not observe strong clustering by sex except for the liver and pectoral fin. Although some differences can be observed in Northern pike and spotted gar, clustering of organs of the nervous system, muscular system, and external organs are generally consistent among species (**Figure S10**).

**Figure 1.**
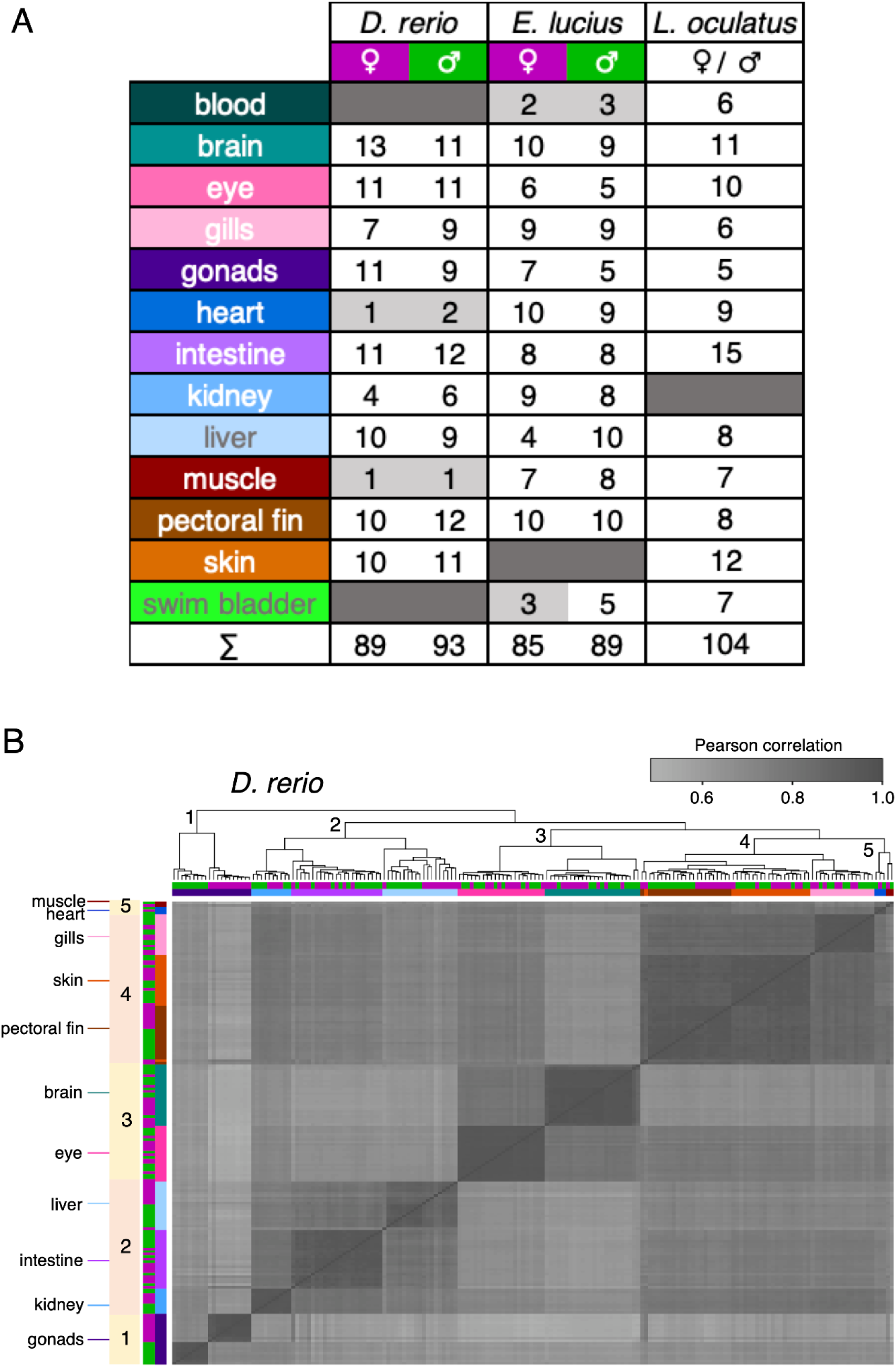
Study design for multi-organ analysis of inter-individual gene expression variability in fishes. **(A)** Summary of BRB-seq dataset. Table shows the number of individuals per organ (rows) and sex (columns) within each species, after removal of low-quality samples. There is no information on sex for spotted gar (*L. oculatus*). Organ-sex conditions with fewer than 4 replicates (light gray) were excluded from expression variability analyses. **(B)** Hierarchical clustering heatmap of organ transcriptomes (*n* = 182) in zebrafish (*D. rerio*). Samples cluster into 5 groups by related organ systems: reproductive system (1), gastrointestinal and urinary system (2), nervous system (3), external organs (4), and muscular system (5). Pairwise distances between samples are computed based on Pearson’s correlation of log-transformed, TMM-normalized counts per million (log_2_ TMM-CPM) of protein-coding genes (*n* = 19,466). The color keys for organ (inner color bars) and sex (outer color bars) are shown in (A).

### Gene expression variability is estimated independently of expression level

Here, we use *variance* (or, more generally, *statistical dispersion*) to refer to a measurable statistical feature of an ensemble, and *variability*’ to refer to a dispositional property (Wagner and Altenberg 1996) inferred from such measurements. Measuring expression variance as a proxy for quantifying variability has not been standardized in the literature, and different metrics have been applied to quantify variance at the bulk or single-cell level (Zheng et al. 2023). Different measures of statistical dispersion (e.g., standard deviation, variance, median absolute deviation (MAD), or coefficient of variation (CV)) are known to be strongly correlated with mRNA abundance (**Figure 2A, Figure S11**) (Bar-Even et al. 2006; Eling et al. 2019). To correct for this mean-dispersion relationship, many studies use regression to model variance as a function of expression level (Eling et al. 2019) and use either the ratio (Alemu et al. 2014; Barroso et al. 2018; Liu et al. 2020) or difference (Sigalova et al. 2020; Bashkeel et al. 2019) (i.e., model residual) of observed to expected variation as an adjusted estimate of expression variability. Although this removes the global correlation between expression mean and variation, this does not correct for the non-homogenous distribution of variation across the range of expression levels (Faure et al. 2017; Simonovsky et al. 2019). To address this, other studies apply percentile rank normalization (e.g., ‘local coefficient of variation’ (LCV) (Simonovsky et al. 2019; Horta-Lacueva et al. 2023), similar to the ‘relative noise rank’ (Faure et al. 2017)), in which the effect of mean expression is removed by ranking the expression variation of each gene relative to a window of genes with similar expression levels. Here, we used the local (residual) coefficient of variation as our estimate of expression variability because it is globally uncorrelated with mean expression level, as in regression-based approaches, and it also imposes a uniform expression variability distribution across expression levels. Although the latter is a very strong correction, by forcing a uniform distribution we maximally remove mean expression as an explanatory factor for downstream analyses.

**Figure 2.**
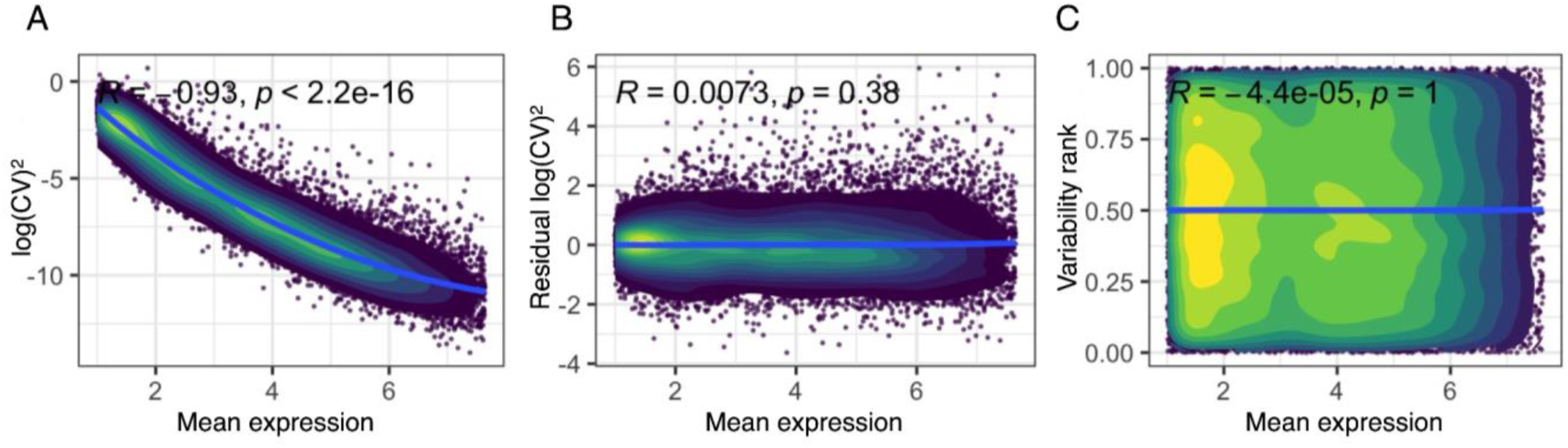
Measures of expression variance as a function of expression level. Data from female zebrafish brain (*n* = 13 samples). All values are means estimated from jackknife (*n -* 1) resampling. Mean expression refers to log-transformed, TMM-normalized counts per million (log_2_ TMM-CPM). **(A)** Log-transformed (squared) coefficient of variation (CV, defined as the ratio of standard deviation to the mean). **(B**) Residuals of log_2_ (squared) CV. **(C)** Local percentile rank of residual log(CV)^2^ (‘variability rank’) within sliding windows of 100 neighboring genes sorted by expression level. Blue lines are LOESS regression curves and *R* is Pearson correlation coefficient.

Since actual expression variance may be skewed by samples with more divergent expression profiles, we computed the LCV for each gene within an organ (for unsexed spotted gar) or organ-sex condition (for sexed Northern pike and zebrafish) using a jackknife resampling or ‘leave-one-out’ procedure to mitigate the effect of outliers. For each subset of *n* - 1 replicates, we first computed the log-transformed, squared coefficient of variation (log(CV)^2^, where CV is the ratio of standard deviation to the mean) and fitted a local polynomial regression (LOESS) to model log(CV)^2^ as a function of mean expression (**Figure 2B**). Second, we ranked the residual log(CV)^2^ (‘residual CV’) of each gene within sliding windows of 100 neighboring genes sorted by expression level, such that genes with rank 0.00 have the lowest dispersion whereas genes with rank 1.00 have the highest dispersion within a local gene neighborhood (**Figure 2C**). Lastly, we calculated the mean LCV over all *n* - 1 subsets within a condition and used this metric, which we call ‘variability rank’, as our estimate of gene expression variability (**Figures S12**-**S14**).

Because variability ranks are computed relative to the set of genes expressed within a specific condition, it is not an absolute measure of variability that is directly comparable between conditions. A gene with a high variability rank in the brain is considered highly variable only with respect to other genes expressed in the brain (e.g., a gene with a variability rank of 0.80 in the brain is not necessarily more variable, in the absolute sense, than a gene with a variability rank of 0.60 in the liver). By construction, variability ranks are thus generally organ-specific, with weak to moderate correlation across organs (**Figures S15**-**S17**).

One limitation of the variability rank metric is that it does not describe the modality of the underlying expression distribution. Overdispersion may be due to either a broader range of expression centered at a single mean (i.e., a unimodal distribution) or the presence of two or more expression peaks (a multimodal distribution), the latter of which may be indicative of hidden substructures within the population (Mar 2019). To investigate this possibility, we assessed whether more bimodal expression profiles can be observed for sets of increasingly more variable genes (**Figure S18**) (Supplementary Results; Supplementary Methods). Overall, we find weak evidence for bimodality in our dataset, only in some conditions, and only at very high variability ranks. Thus, we infer that higher variability is likely primarily driven by a wider range of expression centered around a single peak, rather than by bimodality of expression.

### Gene expression variability is associated with gene function

After applying a strong correction to remove the effect of mean expression, we verified that our estimate of gene expression variability retains a functionally relevant signal. For this, we performed a Gene Ontology (GO) enrichment analysis of GO biological process terms repeatedly represented (*n* ≥ 3 conditions) in genes with either low or high expression dispersion, using a conditional hypergeometric test (implemented in *GOstats* (Falcon and Gentleman 2007)) that corrects for dependencies between GO terms by accounting for the GO graph topology (Falcon and Gentleman 2007; Alexa et al. 2006). We observed that sets of lowly (respectively highly) variable genes show a substantial overlap in enriched GO biological process categories across several conditions. Underdispersed genes (variability rank ≤ 0.20) are consistently enriched in core cellular (housekeeping) functions such as protein localization and transport, protein modification, RNA processing, and organelle organization (**Figure 3A**). Conversely, overdispersed genes (variability rank ≥ 0.80) are enriched in functions such as signaling, chemotaxis, response to stimuli, and immune response (**Figure 3B**). Functional categories that are under-represented in lowly variable genes also tend to be over-represented in highly variable genes, and vice versa (**Figure S25**). We also performed a GO enrichment analysis for moderately dispersed genes (0.40 ≤ variability rank ≤ 0.60) but observed virtually no over-represented terms, and under-representation of terms related to defense response, immune response, and muscle movement (**Figure S26**). To complement these analyses, we tested for GO terms with significantly lowly or highly variable expression based on a competitive gene set test (Wu and Smyth 2012), which accounts for inter-gene correlation and reduces false positives (**Figure S27**). GO terms for lowly variable genes involve RNA processing, whereas those for highly variable genes include terms related to chemotaxis, response to stimuli, muscle contraction, and others. These findings are consistent with previous studies in humans (Alemu et al. 2014; Wolf et al. 2023) and *Drosophila* (Sigalova et al. 2020; Phipps-Tan et al. 2026) which have shown that lowly and highly variable genes are enriched in distinct functional categories.

**Figure 3.**
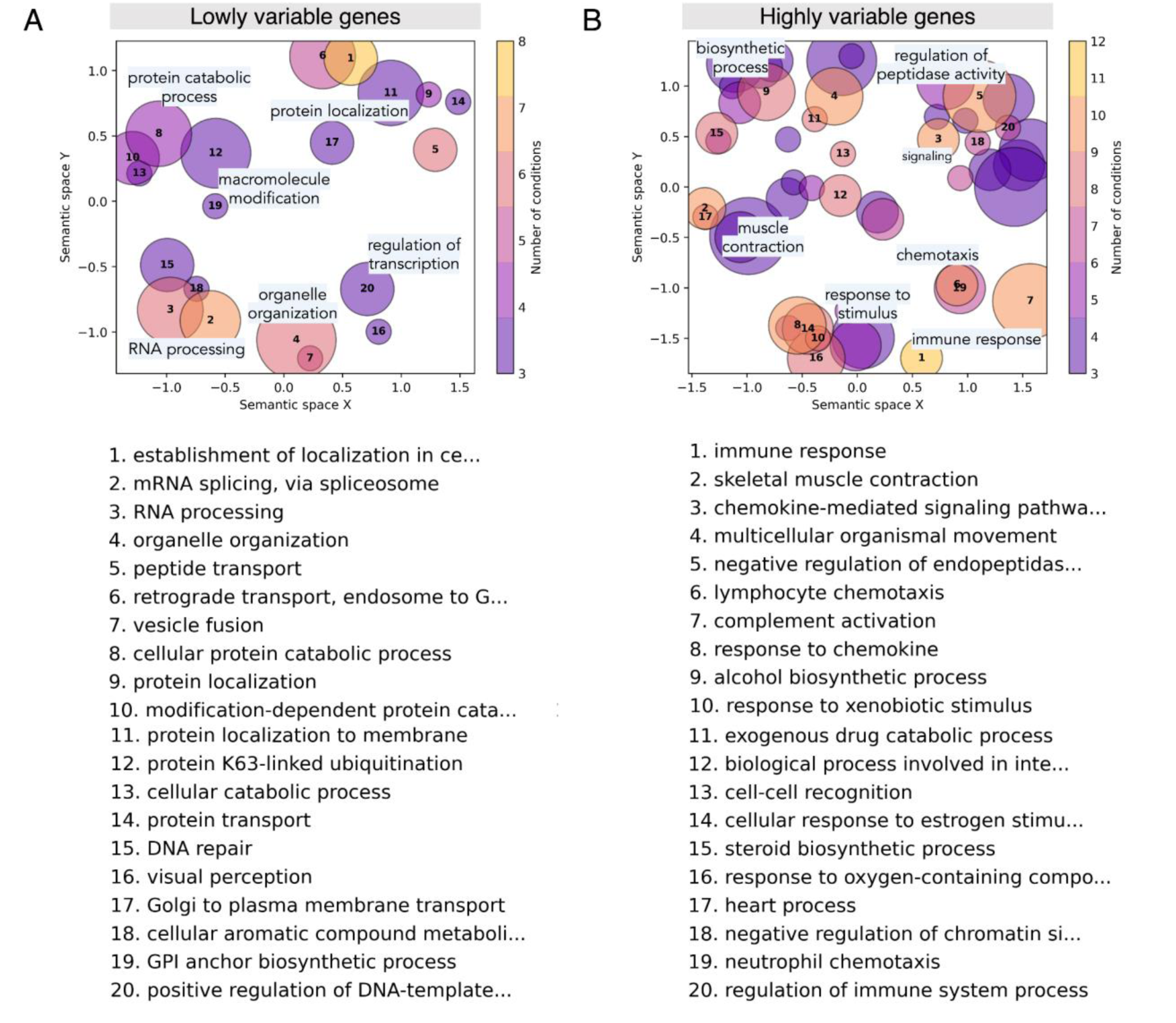
Semantic similarity bubble plots of enriched Gene Ontology (GO) biological process terms for (A) underdispersed (variability rank ≤ 0.20) and (B) overdispersed (variability rank ≥ 0.80) genes in zebrafish organs. Over-representation analysis of GO terms was performed separately for the set of under- and overdispersed genes in each organ-sex condition using the conditional hypergeometric test in *GOstats* (Falcon and Gentleman 2007). Terms that are significant (unadjusted *p*-value < 0.01) in at least three organ-sex conditions were retained. Redundancy reduction and visualization of terms was performed using *GO-Figure*! (Reijnders and Waterhouse 2021). Each circle represents a group of similar GO terms collapsed into a representative term. Representative terms are plotted in a semantic similarity space such that more similar terms are positioned closer together. The top 20 representative terms are listed and sorted by the number of organ-sex conditions in which the representative is significant. Circle color indicates the number of conditions in which a representative term is significant while circle size is scaled by the number of related terms summarized in a cluster.

### Variably expressed genes evolve under weaker purifying selection

By testing for an association between inter-individual gene expression variability and *d*_N_/*d*_s_, we investigated whether variability evolves under selection pressures that run in parallel to those acting on the protein-coding sequence. For this, precomputed selection statistics were retrieved from Selectome (Proux et al. 2009; Moretti et al. 2014), a database of conservative branch-site likelihood tests for selection conducted on vertebrate gene trees (Materials and Methods). For each species, the most underdispersed genes (median variability rank ≤ 0.2 averaged across conditions) have evolved under stronger negative selection (*d*_N_/*d*_s_) compared to the most overdispersed genes (median variability rank > 0.8 across conditions). As there is little variance in *d*_N_/*d*_s_ on the bulk of genes, the global inverse correlation is significant but relatively weak (**Figure 4**; **Figure S20**).

**Figure 4.**
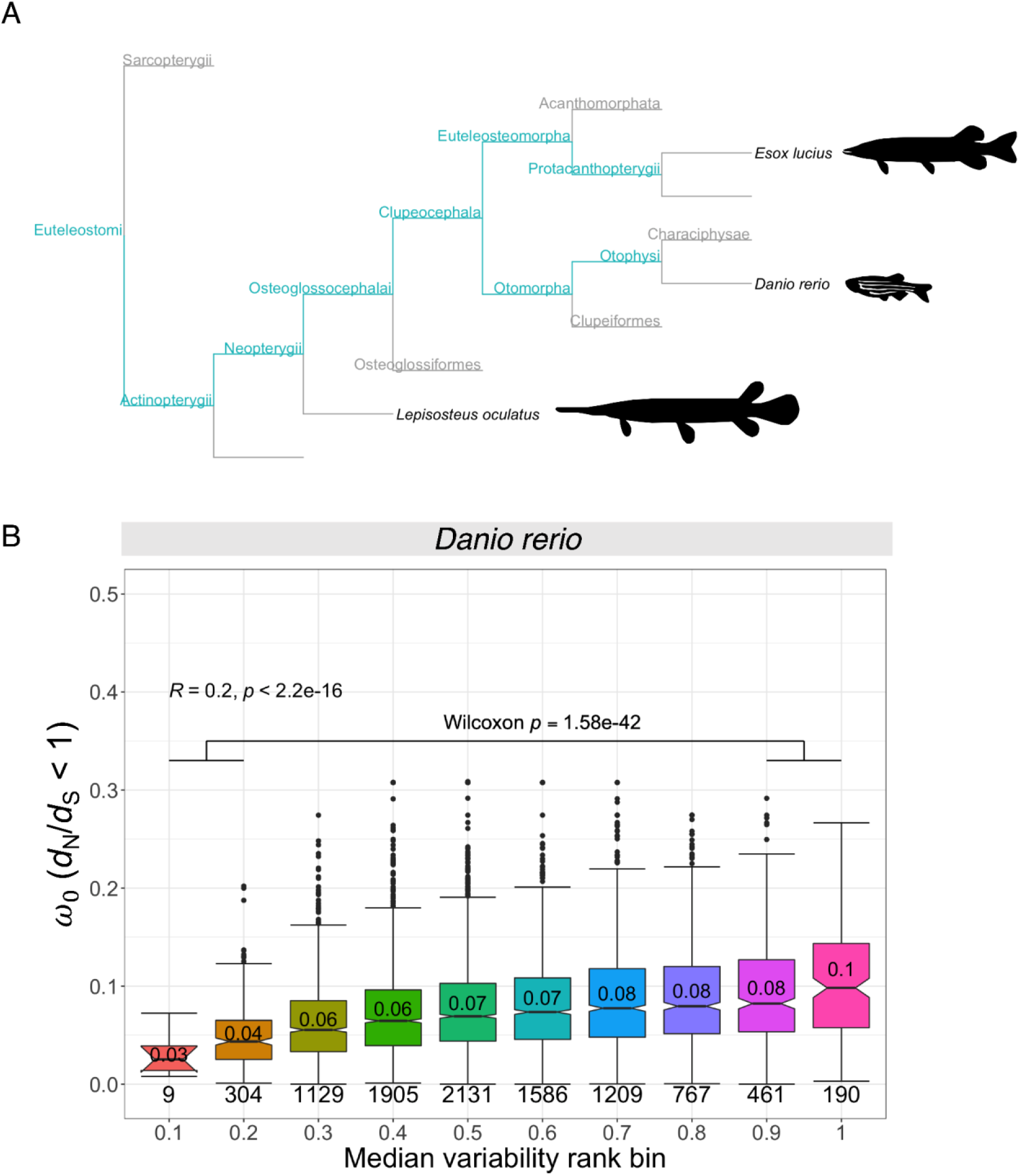
Purifying selection of zebrafish genes grouped by median expression variability across conditions. Selection on coding sequences computed from conservative branch-site likelihood tests were retrieved from the Selectome database (Moretti et al. 2014; Proux et al. 2009). **(A)** Species phylogeny with tested internal branches highlighted (turquoise). Tested branches included in all three species are: Euteleostomi, Actinopterygii, Neopterygii; for teleosts, i.e. zebrafish and pike, only: Osteoglossocephalai, Clupeocephala; for zebrafish only: Otomorpha, Otophysi; and for pike only: Euteleosteomorpha, Protacanthopterygii. Icons retrieved from PhyloPic: *D. rerio* by Jake Warner (CC0 1.0), *E. lucius* by Timothy Knepp and T. Michael Keesey (PDM 1.0), *L. oculatus* by Ingo Braasch (CC0 1.0). Edge lengths are not scaled by divergence time. **(B)** Zebrafish protein-coding genes were grouped into 10 bins of size 0.1 ([0,0.1), (0.1,0.2], …, (0.8,0.9], (0.9,1.0]) based on their median variability rank across organ-sex conditions. Boxplots show the distribution of ratios of nonsynonymous substitutions (*d*_N_) to synonymous substitutions (*d*_S_) of sites that have evolved under purifying selection (***ω***_0_ ratio, for *d*_N_/*d*_S_ < 1). The median ***ω***_0_ is indicated within each boxplot and the number of genes included are annotated below each. *R* is the Spearman’s correlation coefficient between ***ω***_0_ and the unbinned median variability across conditions, and its corresponding *p*-value. The Wilcoxon *p*-value corresponds to a Wilcoxon rank-sum test comparing the least variable genes (variability rank ≤ 0.2) against the most variable genes (variability rank > 0.8). See **Figure S28** for equivalent analyses in Northern pike and spotted gar.

The codon model used explicitly models the strength of negative selection and positive selection separately (Zhang et al. 2005; Davydov et al. 2019). We have previously shown that the strength of purifying selection does not affect the proportion of false positives for positive selection (Gharib and Robinson-Rechavi 2013). Thus, we compared evidence for positive selection for genes with consistently low, moderate, or high expression dispersion across multiple conditions within a species. We observed a statistically significant association between estimated variability and the proportion of branches with evidence for positive selection in zebrafish (Pearson’s chi-squared test, *χ*^2^= 21.866, *p* = 1.786 × 10^−5^) and Northern pike (*χ*^2^= 48.122, *p* = 3.552 × 10^−11^), but not in spotted gar (*χ*^2^= 1.4127, *p* = 0.4935). Specifically, we found depletion of branches under positive selection for frequently underdispersed genes (variability rank ≤ 0.20 in at least half of available conditions) in both zebrafish and pike, as well as enrichment of positive selection for frequently overdispersed genes (variability rank ≥ 0.80 at least half of conditions) in pike (**Table S4**). Thus, higher gene expression variability between individuals is associated with faster protein-coding sequence evolution between species, both due to relaxed negative selection and to increased positive selection.

### Brain-biased genes have low expression variability across non-nervous organs

We have shown that genes which are lowly variable across individuals in multiple conditions are enriched in housekeeping functions, while highly variable genes are enriched in stimulus-sensitive functions (**Figure 3**). Since housekeeping genes are, by definition, ubiquitously expressed across conditions (but see (Joshi et al. 2022) for a discussion), this suggests that the relationship works in both directions: genes expressed broadly across organs should also show low within-organ variability across individuals, consistent with stronger pleiotropic constraints. To test this hypothesis, we quantified expression breadth per gene using the expression specificity metric tau (*τ*), with range [0,1], in which genes with *τ* close to 0 are uniformly expressed across organs whereas genes with *τ* close to 1 are highly organ-specific (Yanai et al. 2005; Kryuchkova-Mostacci and Robinson-Rechavi 2017) (Materials and Methods). Importantly, *τ* only considers mean expression differences between conditions and does not take within-condition expression dispersion into account, thus it is independent of our estimate of variability. Across conditions in each species, results consistently show a positive association between variability rank and organ specificity (**Figures S29**-**S31**). Based on the lower quartile of the *τ* distribution averaged across species, we designated genes with *τ* ≤ 0.3 as ‘broadly expressed’ and genes with *τ* > 0.3 as ‘organ-biased’ (**Figure S32**). Note that we explicitly use the term ‘organ-biased’ rather than ‘organ-specific’ because our relaxed *τ* cutoff necessarily includes genes that are preferentially expressed in one organ but are still expressed in other organs at lower levels. Per organ, broadly expressed genes are generally underdispersed relative to organ-biased genes (**Figure 5A-C**, **Figure S33A-C, Figure S34A-C, Figures S35-S37**, **Figures S38-S40**). Similar results were obtained with a more stringent *τ* cutoff (*τ* > 0.5 as organ-biased) and with batch correction via *ComBat-seq* (Zhang et al. 2020) (**Figures S41-S43**, **Figures S44-S46**). These results are consistent with previous studies suggesting that expression variability is associated with both tissue and cell-type-specific expression (Fair et al. 2020; Einarsson et al. 2022).

**Figure 5.**
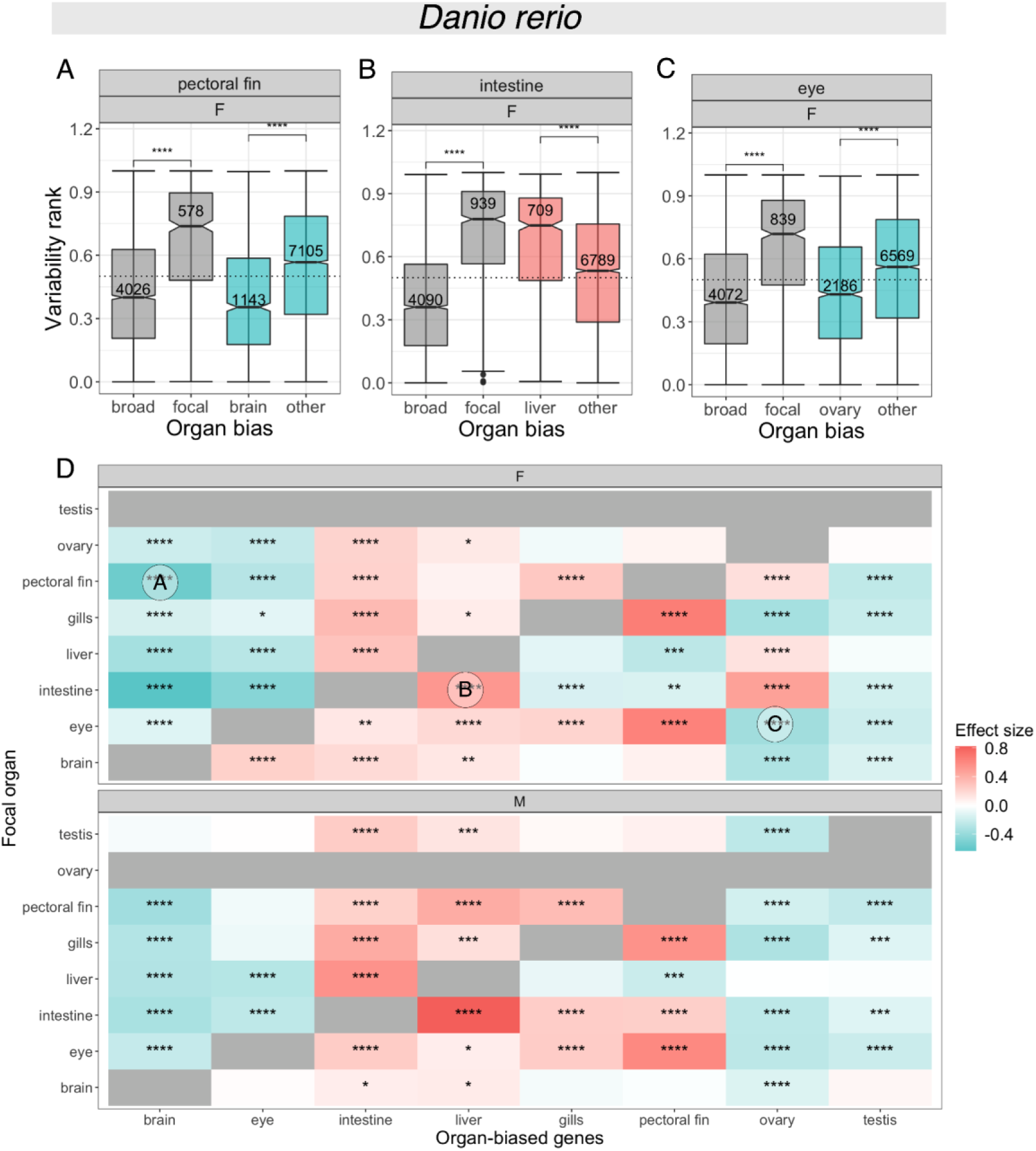
Relationship between expression variability and organ-biased expression for zebrafish. **(A)** Boxplot of expression variability ranks of genes expressed in female (‘F’) zebrafish pectoral fin, for sets of genes classified by organ bias and with an emphasis on brain-biased genes. Broadly expressed genes (‘broad’) have a measured organ expression specificity index (tau) of τ ≤ 0.3; genes with τ > 0.3 are considered organ-biased. Among the set of organ-biased genes, genes are further classified into three categories: focal-biased genes (‘focal’) are preferentially expressed in the focal organ (i.e., pectoral fin), brain-biased genes (‘brain’) are preferentially expressed in the brain, and the remaining (‘other’) are preferentially expressed in other organs. The number of genes included are annotated for each category. Comparisons between broadly expressed and focal-biased genes, and between brain-biased and other organ-biased genes were performed using Wilcoxon rank-sum tests (*p*-value significance levels: ‘****’: *p* ≤ 0.0001, ‘***’: *p* ≤ 0.001, ‘**’: *p* ≤ 0.01, ‘*’: *p* ≤ 0.05, ‘ns’: *p* > 0.05). The dashed line indicates a variability rank of 0.5. **(B)** Boxplot of variability ranks of genes expressed in female zebrafish intestine, with genes classified by organ bias and with an emphasis on liver-biased genes. **(C)** Boxplot of variability ranks of genes expressed in female zebrafish eye, with genes classified by organ bias and with an emphasis on ovary-biased genes. **(D)** Heatmap summarizing pairwise comparisons of expression variability ranks between subsets of organ-biased genes for female (‘F’, top) and male (‘M’, bottom) zebrafish. Rows correspond to focal organ whereas columns correspond to subsets of organ-biased genes. Pairwise comparisons were performed using Wilcoxon rank-sum tests with Benjamini-Hochberg (‘BH’) correction for multiple testing (adjusted *p*-value < 0.05). Effect size was computed using Glass’s delta (δ), with color scale shown in (L). Cells in gray are not included in the dataset. Three cells are annotated, corresponding to the following comparisons in boxplots (A-C): (A) brain-biased vs. other organ-biased genes expressed in female zebrafish pectoral fin, (B) liver-biased vs. other organ-biased biased genes expressed in female zebrafish intestine, and (C) ovary-biased vs. other organ-biased genes expressed in female zebrafish eye. A negative effect size entails lower expression variability relative to other organ-biased genes (A,C) (turquoise), whereas a positive effect size entails higher variability (B) (salmon). See **Figure S33** for equivalent analyses in Northern spike and **Figure S34** for spotted gar.

Since organ-biased genes have likely evolved under selection for an organ-specific function, one prediction is that these genes would have a narrower expression range within their top (focal) organ of expression, indicating more fine-tuned regulation for its putative primary function. On the contrary, we observed significant overdispersion of organ-biased genes within their focal organ. For example, genes that are most highly expressed in the pectoral fin are overdispersed within the pectoral fin, relative both to broadly expressed genes as well as to other organ-biased genes expressed in this organ (**Figure 5A, Figure S33A, Figure S34A**). This is consistent with a previous claim that highly variable genes within a cell type population are typically indicative of cell type function (Osorio et al. 2020). This pattern is lost when the top organ is assigned randomly, such that focal-biased genes have dispersion levels that are indistinguishable from the background of other organ-biased genes (**Figures S47-S48**). An alternative prediction of the selection on expression of organ-biased genes is that they would vary according to the fine physiological state of the organ in each individual. To test this, we investigated co-variation patterns, under the assumption that organ state changes would lead to coordinated change in genes which function together. Indeed, focal-biased genes show stronger evidence for co-variation in expression levels relative to broadly expressed and other organ-biased genes, based on the proportion of genes assigned to modules after a weighted gene co-expression network analysis (WGCNA) (**Figure S49**). Thus, overdispersion of focal-biased genes likely largely reflects differences in physiological state between individuals, whereas dispersion in other organ-biased genes reflects more stochastic variation. We conclude that expression of organ-biased genes within their focal organ tends to be highly plastic in response to genetic or environmental cues, and that context-dependent upregulation of expression of organ-biased genes may be mediated by *trans*-acting factors that amplify variation.

Patterns of expression dispersion across organs can reveal the extent to which the inter-individual expression variability of a gene is independently regulated per condition (Wolf et al. 2023). On the one hand, if control of variability is organ-specific, dispersion should differ across organ contexts. On the other hand, if variability is controlled by a shared underlying mechanism, dispersion should appear consistent across organs. The second scenario entails differences in the evolution of regulatory architecture depending on the top organ of a gene. We observe a consistent pattern across organ contexts that is dependent on the focal organ. For example, we find that brain-biased genes have consistently underdispersed expression across non-nervous system organs relative to other organ-biased genes, regardless of differences in cell type composition per organ (**Figure 5D, Figure S33D, Figure S34D, Figures S35-S37**). (See **Figure S41D, Figure S42D, Figure S43D** for the same analysis given a cutoff of *τ* > 0.5 and batch correction.) Likewise, we found that genes preferentially expressed in the gonads, especially the ovary, tend to be underdispersed across organs **(Figures S38-S40).** By contrast, liver-biased genes, which evolve under weaker purifying selection at the protein-coding sequence level (**Figure S50**) (Kryuchkova-Mostacci and Robinson-Rechavi 2015), tend to be overdispersed across multiple organs (**Figures S51**-**S53**), implying weaker global expression constraints. Taken together, these results are consistent with the expected pattern if gene-specific expression variability is regulated by a shared mechanism across organ contexts (Wolf et al. 2023) that is dependent on the top organ of a gene. We observed qualitatively similar organ-dependent patterns of expression dispersion after controlling for differences in transcriptome complexity (i.e., number of genes expressed) across organs (Ramsköld et al. 2009; Soumillon et al. 2013; Darbellay and Necsulea 2020) (**Figure S54**), with particularly robust results in zebrafish (**Figure S55D**), although the effect is weaker in Northern pike (**Figure S55H**) and spotted gar (**Figure S55L**). The pattern is consistent with a more stringent tau cutoff for organ-biased genes (*τ* > 0.5 as organ-biased), a different variability metric (residual MAD), and without jackknife resampling (**Figures S56**-**S58**). Conversely, no pattern is recovered in any of the species when the top organ is assigned randomly (**Figures S59**-**S61**), confirming that these results are biological and not due to biases in data generation or processing. These results suggest that organ-biased genes have evolved particular *cis*-regulatory mechanisms to facilitate either highly stable expression downregulation across other organs (for brain- and gonads-biased genes) or more variable expression (for liver-, pectoral fin-, and intestine-biased genes). Indeed, we observed that zebrafish promoter regions for each set of organ-biased genes can be distinguished from both matched random promoter sequences (**Figure S62**) and from each other (**Figure S63**) (Methods). Promoter regions from zebrafish brain-, ovary-, and testis-biased genes yield better model performance, indicating that these promoter regions share common sequence motifs that make them more distinguishable from random sequences, consistent with stronger promoter sequence constraints for lowly variable genes. By contrast, promoters regions from pectoral fin-, gill-, liver-, and intestine-biased genes have somewhat lower model performance than sets of random promoters, suggesting weaker sequence constraints (**Figure S64**).

Suboptimal expression can be more or less deleterious in different organs, due to differences in physiological function, cellular composition and turnover, and sensitivity to perturbations (Gu and Su 2007). In particular, it has been suggested that neural tissues are highly sensitive to protein misfolding, and that this sensitivity contributes to strong evolutionary constraints of genes expressed in the brain (Drummond and Wilke 2008). We confirm that protein-coding sequences of brain-biased genes have evolved under stronger purifying selection not only compared to other organ-biased genes, but also to broadly expressed genes (**Figure S50**), which is consistent with reports of strong negative selection on the mammalian brain relative to other organs (Khaitovich et al. 2005; Brawand et al. 2011; Necsulea and Kaessmann 2014; Kryuchkova-Mostacci and Robinson-Rechavi 2015; Cardoso-Moreira et al. 2019). We observed moderate rates of purifying selection on both testis- and ovary-biased genes (**Figure S50**), though highly testis-specific genes have been shown to be faster-evolving compared to other organ-specific genes in mammals (Kryuchkova-Mostacci and Robinson-Rechavi 2015). Overall, our findings show a generally positive association between inter-species protein sequence divergence and intra-species expression variability (**Figure 4**), suggesting that evolutionary constraints at the protein sequence level co-vary with organ-dependent patterns of expression variability. One interesting exception involves eye-biased genes, which evolve under strong purifying selection across the three fish species (**Figure S50**) but which tend to have somewhat overdispersed expression in pike (**Figure S33**) and spotted gar (**Figure S34**). We suggest that the differential variability of eye-biased genes may be an interesting subject for future research, particularly given the remarkable diversity of visual opsin gene repertoires in fishes and its relation to differences in light environment and ecology (Musilova et al. 2021).

From these results, we infer that (1) organ-biased genes are highly variable within their focal organ, likely due to context-sensitive expression, but that (2) their global variability reflects organ-dependent evolutionary constraints on both protein-coding and *cis*-regulatory sequences. We found that these organ-dependent patterns of expression variability are largely consistent in all three sampled species, thus constituting an evolutionarily conserved signal across at least 320 million years of evolution.

## Discussion

Until recently, most gene expression studies have focused on testing for differences in average expression levels between groups of interest, largely to the exclusion of expression variance and variability. Yet a more complex picture is now emerging: gene expression variability is increasingly implicated in disease (Ho et al. 2008; Li et al. 2010; Mar et al. 2011; Dinalankara and Bravo 2015) and drug efficacy (Simonovsky et al. 2019), aging (Bashkeel et al. 2019), circadian rhythm (Cortijo et al. 2019), cellular differentiation (Simon et al. 2018; Ohnishi et al. 2014; Richard et al. 2016), embryogenesis (Liu et al. 2020; Hasegawa et al. 2015), developmental canalization (Horta-Lacueva et al. 2023), and other biological processes. Here, we apply an evolutionary perspective based on a comparative analysis of inter-individual gene expression variability across multiple adult organs in three species of ray-finned fishes. We confirm that genes which are lowly variable among individuals are typically enriched in housekeeping functions, whereas highly variable genes are enriched in stimulus-sensitive functions, as reported previously in other species (Sigalova et al. 2020; Alemu et al. 2014; Wolf et al. 2023; Einarsson et al. 2022). This biologically intuitive result reaffirms that expression variability is a functionally relevant trait independent of average expression. There is a growing body of literature that recognizes the functional relevance of expression variability (Geiler-Samerotte et al. 2013; De Jong et al. 2019; Mar 2019) and how this variability is associated with promoter characteristics such as sequence, core promoter motifs, and transcriptional start site architecture (Sigalova et al. 2020; Einarsson et al. 2022; Schor et al. 2017). However, there have been few attempts to characterize how gene expression variability in animals is systematically modulated by the interplay between gene function and organ context, though some studies have sampled diverse organs (Simonovsky et al. 2019; Wolf et al. 2023). To our knowledge, this is one of the first studies to investigate how gene expression variability is influenced both by expression specificity and organ context.

Our cross-organ approach helps clarify the connection between two previously reported findings. First, although lowly and highly variable genes show distinct functional signatures, some studies report that gene expression variability tends to be tissue-specific (i.e., organ-specific) (Wolf et al. 2023), with little overlap in the set of most overdispersed genes between organs (Bashkeel et al. 2019; Alemu et al. 2014) or cell types (Osorio et al. 2020) (but see (Sigalova et al. 2020)). Second, organ- or cell-type-specific genes are found to have highly variable expression (Fair et al. 2020; Einarsson et al. 2022; Osorio et al. 2020). We verify that organ-biased genes are more variably expressed relative to broadly expressed genes, but in addition, we have also found that organ-biased genes are typically highly variable within their primary (focal) organ. Since the most variable genes within an organ are typically genes which are biased or specific to it, we think it is unsurprising to find few highly variable genes shared between multiple conditions. We infer that plastic expression within a focal organ is a functional feature of more specialized genes, and that expression is dynamically induced by context-specific regulatory cues, such as organ-specific transcription factor binding to enhancer elements or enhancer-promoter interactions. Promoters of specialized genes (e.g., developmental genes) have been shown to interact with different classes of enhancers compared to those of housekeeping genes (Zabidi et al. 2015). In addition, such specialized genes tend to have more enhancer-promoter interactions than housekeeping genes (Schoenfelder and Fraser 2019; Furlong and Levine 2018), and the number of interacting enhancers has been found to have an additive effect on gene expression levels (Schoenfelder and Fraser 2019; Javierre et al. 2016; Schoenfelder et al. 2015). Thus, to augment transcription of organ-biased genes in the relevant context, we expect an increase in regulatory inputs, which introduces more sources of expression variation (Urchueguía et al. 2021). Since gene-specific expression variability is influenced by a gene’s regulatory inputs (Urchueguía et al. 2021) and its position in the gene regulatory network (Puzović et al. 2023), we expect subsets of context-dependent genes to show co-variation in expression levels. Indeed, that is what we observe, with patterns of expression co-variation of organ-biased genes in the focal organ, and of broadly expressed but highly variable genes (**Figure S49**), putatively corresponding to variation in the physiological status of that organ between individuals.

Outside of their focal organ, organ-biased genes have basal expression levels that are relatively more stable among individuals. Moreover, their expression patterns show low co-variation; i.e., variability of organ-biased genes in a non-focal context appears to be largely a stochastic property of individual genes rather than of gene networks. We find evidence across species that the degree to which biased genes are stably downregulated across non-focal organs reflects organ-dependent evolutionary constraints. Notably, we observe consistently underdispersed expression of brain-biased genes across non-nervous organs, despite these organs having very different transcriptional repertoires, cell-type compositions, and local micro-environments. Low variability within a condition suggests stabilizing selection on expression, which is consistent with strong purifying selection on *cis*-regulatory sequences, and/or with buffering of expression variation by compensatory *cis*-*trans* effects (Signor and Nuzhdin 2018). Given that we observe consistent underdispersion across multiple organs having very different local micro-environments, we reason that stable basal expression of brain-biased genes is likely to be predominantly driven by strong purifying selection at the *cis*-regulatory level rather than by context-dependent *trans* effects. We also observed that gonads-biased genes have lowly variable expression in non-reproductive organs, which is consistent with recent findings that show that lowly variable genes in the *Drosophila* head are highly enriched in genes related to reproduction (Phipps-Tan et al. 2026). Furthermore, our finding that organ-biased genes have distinguishable promoter sequences (**Figure S62**) supports the view that inter-individual variability across organ contexts is at least partly under *cis*-regulatory control. Moreover, these results support stronger selection on promoters that regulate lower variability in non-focal organs.

We recognize that comparing gene expression variability between organs is challenging because any measured changes in dispersion may simply be due to technical variation (e.g., sampling design, sequencing depth or RNA quality (Simonovsky et al. 2019)) or ‘extrinsic’ sources of biological variation (e.g., cell type composition (Fair et al. 2020; McCall et al. 2016) or transcriptome complexity (Ramsköld et al. 2009; Soumillon et al. 2013)). Indeed, a limitation of bulk RNA-seq is that it does not allow us to distinguish between multiple sources of variation within an organ, whether it is expression variation within a cell type population, expression variation between cell type populations, or heterogeneity in cell type composition between individuals. Observing a consistent signal of underdispersion across conditions with heterogeneous backgrounds is thus particularly informative and suggests that our method captures a signal of stable *cis*-mediated downregulation that is independent of organ-context. We infer that brain-biased genes have evolved under strong purifying selection, not only at the protein-coding sequence level, but also at the *cis*-regulatory level. From this, we hypothesize that genes selected for function in the brain may have co-evolved with more stringent ‘enhancer grammar’ (e.g., number, type, order, affinity, spacing, or orientation of transcription factor binding sites) (Long et al. 2016; Jindal and Farley 2021) relative to other organ-biased genes. Our results highlight how analysis of gene expression variability across organs can yield additional insights into gene regulatory strategies in animals. Extending on these findings, we propose that intra-specific expression variability can potentially be informative of broader macro-evolutionary trends, such as patterns of duplicate gene retention and functional divergence after whole-genome duplication.

## Materials and Methods

### Sample preparation

#### Sample collection

##### Zebrafish

Thirteen female and twelve male zebrafish (*Danio rerio*), strain AB, were raised at the University of Oregon Aquatic Animal Care Services. Animals were raised in E5 medium (Westerfield 2007) and fed rotifers from 4-10dpf (Best et al. 2010). After 10dpf, fish were moved to a circulating fish water system and fed Zeigler Adult Zebrafish Diet. Three-month old adults were euthanized by cold shock (Westerfield 2007), and organs were dissected and immediately stored in RNAlater. Procedures were approved by the University of Oregon IACUC protocol #18-13.

##### Northern pike

Ten female and ten male one-year-old Northern pike (*Esox lucius*) were sourced from a commercial hatchery (Aqua 2B, Pouancé, France). The fish were euthanized using a lethal dose of tricaine methanesulfonate (MS 222, 400 mg/L), supplemented with 150 mg/L of sodium bicarbonate. Following euthanasia, organs were immediately dissected and preserved in RNAlater. All animal procedures were conducted in full compliance with French and European legislation (French decree 2013-118 and European Directive 2010-63-UE), governing the ethical use and care of laboratory animals in scientific research.

##### Spotted gar

Spotted gars (*Lepisosteus oculatus*) were obtained as embryos from hormone-induced spawns of wild-caught broodstock from bayous near Thibodaux, Louisiana and raised at Michigan State University for 23 or 31 months (April 2018 - March/November 2020) in 150 to 300 gallon tanks on a 12h light : 12h dark cycle and a diet starting from *Artemia* brine shrimp to feeder fish (mostly fathead minnows). Twenty-two (22) unsexed, pre-mature individuals of 20.1 - 23.3 cm standard length (tip of snout to caudal peduncle) and 24.8 - 29.0cm total length (tip of snout to tip of caudal fin) were euthanized in 300 mg/L MS-222 (Sigma). Tissues were dissected and stored in RNAlater (Invitrogen) at −80℃ until shipped frozen to the University of Lausanne. All spotted gar experiments were approved by the Institutional Animal Care & Use Committee (IACUC) of Michigan State University (animal use protocol PROTO201900309).

#### RNA extraction

Tissue samples were thawed on ice, removed from RNA*later* RNA Stabilization Solution (Invitrogen, catalog no. AM7024) and transferred to new Eppendorf tubes containing 1 mL TRIzol Reagent (Invitrogen, ref. 15596018) and about 10 - 15 SiLi beads (Type ZS, 1.4 - 1.6 mm) (Sigmund Lindner, ref. 9315-33). Excluding blood, samples were homogenized at 4°C using PreCellys Evolution (Bertin Technologies, ref. P000062 _PEV0) for at least 2 cycles at 6600 rpm for 30 s. Phase separation of homogenized samples was performed following the TRIzol Reagent user guide (MAN0001271). For Northern pike and spotted gar, RNA extraction with TRIzol Reagent was carried out following the same manual. After washing and resuspending RNA, samples were treated with RNase-free DNase I (Thermo Scientific, catalog no. EN0525) to remove DNA in accordance with the user guide (MAN0012000). For zebrafish, column-based RNA extraction with TRIzol was performed using the PureLink RNA Mini Kit (Invitrogen, ref. 12183018A), following manufacturer’s instructions (MAN0000406). DNA digestion during on-column RNA purification was done using the PureLink DNase Set (Invitrogen, ref. 12185010). All samples were eluted in 50 - 60 μl RNase-free water. Following extraction, RNA concentration was quantified with the QuantiFluor RNA System (Promega, ref. E3310). A subset of samples was also assessed for RNA quality using the 5200 Fragment Analyzer System (Agilent Technologies, M5310AA) with PROSize 3.0.1.6 data analysis software (Agilent Technologies). Samples with low RNA concentration or signs of degradation were removed from subsequent steps. RNA preparations were diluted to a minimum concentration of 50 ng/μl, transferred to low-binding 96-well plates, and stored in −80°C prior to sequencing.

### Data generation

#### Bulk RNA barcoding and sequencing (BRB-seq)

Samples were sent to Alithea Genomics (Lausanne, CH) for library preparation and sequencing using highly multiplexed 3′-end bulk RNA barcoding and sequencing (BRB-seq) (Alpern et al. 2019) over three phases (corresponding to projects AG0012, AMP0027, and AMP0020) (**Table S1**) (Supplementary Data). BRB-seq libraries were prepared using the MERCURIUS BRB-seq library preparation kit for Illumina and following the manufacturer’s manual (Alithea Genomics, #18013). For project AG0012 (Northern pike, spotted gar), libraries were sequenced using Illumina NovaSeq 6000, generating multiplexed, asymmetric paired-end reads, where read R1 (25 bp) contains the sample barcode and UMI and read R2 (80 bp) contains the cDNA fragment sequence to be aligned to the reference genome. For projects AMP0027 (spotted gar) and AMP0020 (zebrafish), libraries were paired-end sequenced using Illumina NextSeq 550 (R1 = 21 bp, R2 = 55 bp).

#### Read alignment and quantification

Read trimming, alignment, and quantification steps were performed by Alithea Genomics using STARsolo (preprint) (Kaminow et al. 2021) from STAR 2.7.9a (Dobin et al. 2013). R2 reads were trimmed (“--clipAdapterType CellRanger4”) and aligned to the following reference genomes: for spotted gar, LepOcu1 (Ensembl 103 (Yates et al. 2020)); zebrafish, GRCz11 (Ensembl 101 (Yates et al. 2020)); Northern pike, Eluc_v4 (Ensembl 101). To generate both raw and UMI-deduplicated counts, the following parameter was used: “--soloUMIdedup NoDedup 1MM_Directional”. UMI-deduplicated count matrices were used for downstream analyses.

### Data analysis

Excluding the promoter sequence analysis, data analyses were performed using R 4.1.0 (R Core Team 2021), Bioconductor 3.13 (Huber et al. 2015), and RStudio 2023.03.0+386 (RStudio Team 2021) unless specified otherwise.

#### Sample filtering

Sample libraries were filtered in two main steps independently for each species. First, samples with low sequencing quality were removed based on a minimum number of uniquely mapped reads or number of genes with nonzero counts (“detected genes”), following Liu et al. (Liu et al. 2020). For example, in zebrafish, samples with less than 500,000 uniquely mapped reads or less than 10,000 detected genes were excluded from downstream analysis. To account for differences in reference genome annotation quality, we selected species-specific minimum uniquely mapped reads and detected genes. (See **Table S2** for filtering criteria.) For each species, we visually inspected the plot of detected genes against uniquely mapped reads to estimate the point in which the relationship between the two variables becomes nonlinear and set this point as the minimum threshold (**Figure S1A**, **Figure S2A**, **Figure S3A**). We verified that samples which do not meet specified minimum criteria are visible outliers on a principal component analysis (PCA) (**Figure S1B-C**, **Figure S2B-C**, **Figure S3B-C**).

Second, samples that were poorly correlated with other samples within its annotated organ were removed. We performed iterative removal of lowly correlated samples based on a protocol described by Fukushima & Pollock (Fukushima and Pollock 2020). For each iteration, we first generated the all-against-all pairwise Pearson’s correlation matrix of retained samples based on gene-filtered and normalized expression data in the form of log-transformed, TMM-normalized counts per million (log_2_ TMM-CPM) (Gene filtering and normalization). Then, for each sample, we computed its mean Pearson’s correlation with each organ. We removed any sample which is more highly correlated with another organ than the one in which it is annotated to. This process was repeated until no more samples fit removal criteria. Considering the set of protein-coding genes in each species, we verified that all retained samples cluster by organ based on hierarchical clustering of the pairwise Pearson’s correlation matrix (**Figure 1B, Figure S10**) and PCA of gene-filtered and normalized expression (**Figures S7-S9**). Gene biotype information was retrieved for each species from Ensembl 105 (Howe et al. 2021).

For computing expression specificity (Organ expression specificity), we required a minimum of 1 replicate per organ-sex for Northern pike and zebrafish, and 2 replicates per organ for spotted gar (**Table S2**). For estimating expression variability (Gene expression variability) and testing its association with gene function (Gene Ontology (GO) statistical over- and under-representation analysis) and coding-sequence selection (Selection analysis), we required a minimum of 4 replicates per organ (for spotted gar) or organ-sex condition. Any conditions which did not meet the minimum number of replicates after quality filtering were excluded from downstream analysis (**Table S3**). See **Tables S2**-**S3** for samples retained after each step of the sample filtering pipeline and **Figure 1A** for a complete breakdown of the number of replicates retained per condition.

#### Gene filtering and normalization

At the start of each step of the sample filtering pipeline (Sample Filtering), we removed non-expressed genes from the full raw count matrix by retaining only genes with both mean and median counts per million (CPM) greater than 1.0 in at least one organ. From the gene-filtered raw count matrix, we performed library size normalization over all conditions using the trimmed mean of M-values (TMM) method in *edgeR* (3.34.1) (Robinson et al. 2009), then re-converted counts to CPM and transformed CPM to log_2_-scale (log_2_ TMM-CPM). To remove negative values, log_2_ TMM-CPM was vertically offset by the absolute value of the global expression minimum such that the minimum value in the normalized expression matrix is always 0.

#### Gene expression variability estimation

To estimate gene expression variability within an organ (for spotted gar) or organ-sex condition (for Northern pike and zebrafish), we considered only the set of conditions with at least 4 replicates after sample filtering. Gene filtering and normalization steps were performed as described in the previous section, but with each condition treated separately. As an additional filter to remove lowly expressed genes specific to a condition, we removed genes with mean or median log_2_ TMM-CPM < 1.0. Since actual expression variation may be sensitive to outliers, we used a jackknife resampling (‘leave-one-out’) procedure to compute expression dispersion statistics and generate a less biased estimate of gene expression variability. That is, for a given condition with *n* replicates, we formed all *n* possible subsets of *n* - 1 replicates and computed expression dispersion metrics for each subset, then computed the mean for each statistic over all subsets. We initially considered different measures of expression dispersion (Supplementary Methods), with the aim to correct for the strong covariation between expression variation and average expression (**Figure S11**). From the metrics we considered, we selected the local (residual) coefficient of variation (LCV) (Faure et al. 2017; Simonovsky et al. 2019) as our estimate of expression variability, as it maximally removes the association between mean expression level and expression dispersion (**Figure 2C; Figures S12**-**S14**). The LCV was computed in two steps then averaged over all *n* - 1 subsets, as described below.

##### Residual coefficient of variation

The coefficient of variation (CV) is the ratio of the standard deviation to the mean. First, we computed the squared CV and applied a log_2_ transformation (log_2_(CV)^2^) for each gene, following Faure et al. (Faure et al. 2017). Next, using the ‘loess’ function from the *stats* package in R, we fit a LOESS regression curve (with span = 0.6 and default parameters) to model log_2_(CV)^2^ as a function of mean expression level. The difference between observed and predicted log_2_(CV)^2^ is the residual log_2_(CV)^2^ (‘residual CV’, for brevity) after adjusting for expression level. Genes with residual CV > 0 overdispersed, while genes genes with residual CV < 0 are underdispersed.

##### Local (residual) coefficient of variation (LCV)

Given residual CVs for each gene, we performed a percentile rank normalization step following the ‘local coefficient of variation’ (Simonovsky et al. 2019; Horta-Lacueva et al. 2023) or ‘relative noise rank’ (Faure et al. 2017) metric recommended in previous publications. First, genes were sorted by mean expression level. Next, we defined a sliding window of 100 neighboring genes and a step size of 1, with a focal gene at the center of each window. At each iteration of the sliding window, the residual CV of the focal gene was ranked against the residual CVs of genes within the window, on a percentile rank scale of 0.00 (lowest dispersion) to 1.00 (highest dispersion). We computed the LCV statistic only for complete windows; as such, for each *n* - 1 subset, we do not have variability estimates for the 50 most lowly and 50 most highly expressed genes.

##### Summarizing jackknife results

Since transcriptome-level expression distributions are positively skewed, with the most highly expressed genes spanning a wide expression range, expression variability tends to be poorly estimated for outlier genes with very high expression. As such, for each condition, we removed the top 5% most highly expressed genes based on jackknifed mean expression level. Then, we computed the mean LCV over all *n* - 1 subsets and used this metric, which we call ‘variability rank’, as our estimate of expression variability.

#### Gene Ontology (GO) statistical over- and under-representation analysis

For each organ-sex condition in zebrafish, we retrieved three sets of genes categorized by expression variability: genes with variability rank ≥ 0.80 (highly variable), genes with rank ≤ 0.20 (lowly variable), and genes with 0.40 ≤ rank ≤ 0.60 (moderately variable). For each variability category per condition, we used *GOstats* (2.58.0) (Falcon and Gentleman 2007) to conduct conditional hypergeometric tests for statistical over-and under-representation of GO biological process terms, based on the org.Dr.eg.db (3.13.0) zebrafish genome annotation (Carlson 2021). Per test, the background gene set was defined as the set of genes which have expression variability estimates within the organ-sex condition. For each variability category, we retained GO terms that were significantly over-represented (or under-represented) (unadjusted *p*-value < 0.01) in at least three conditions. Redundancy reduction and visualization of GO terms was done using the Python package *GO-Figure!* (1.0.0) (Reijnders and Waterhouse 2021) (**Figure 3, Figures S25**-**S26**).

As an alternative to performing GO enrichment tests on pre-defined sets of lowly or highly variable genes, we tested whether GO terms are significantly lowly or highly variable based on a competitive gene set analysis implemented using *cameraPR* from *limma* (Wu and Smyth 2012). Sets of GO terms and their mapping to genes were retrieved from org.Dr.eg.db (3.13.0). Of these, terms containing at least 20 and at most 1000 genes were used to define gene sets (*n* = 3628 terms across BP, MF, and CC ontologies) used as input to *cameraPR*. Per organ-sex condition, we used variability rank as the precomputed statistic and applied a rank-based test. Separately for GO terms associated with highly or lowly variable expression (FDR < 0.05), we retained terms that were significant in at least three conditions and summarized the results using *GO-Figure!* (Reijnders and Waterhouse 2021) (**Figure S27**).

#### Coding-sequence selection analysis

Given a list of protein-coding genes (Ensembl 105 (Howe et al. 2021)) and taxonomic branches for each species, we retrieved precomputed selection statistics from Selectome (https://selectome.org/), a database of evidence for positive selection based on a conservative branch-site likelihood test applied to *Euteleostomi* gene trees (Ensembl 98) (Proux et al. 2009; Moretti et al. 2014). For each gene - internal branch (i.e., taxonomic clade), the Selectome dataset provides a measure of selective pressure for sites under purifying selection (*ω_0_*ratio for sites with *d*_N_/*d*_S_ < 1), as well as a likelihood-ratio test statistic for positive selection. The tested internal branches for each species are as follows. For spotted gar: Euteleostomi, Actinopterygii, Neopterygii. For zebrafish: Euteleostomi, Actinopterygii, Neopterygii, Osteoglossocephalai, Clupeocephala, Otomorpha, Otophysi. For Northern pike: Euteleostomi, Actinopterygii, Neopterygii, Osteoglossocephalai, Clupeocephala, Euteleosteomorpha, Protacanthopterygii (**Figure 4A**). We included only distinct gene - internal branch entries in our downstream analysis, as duplicate gene - internal branch entries may be a result of uncertainties in the gene tree. For each species, the background gene set consisted of protein-coding genes that are included in the Selectome dataset and that have expression variability data in at least half of the available conditions.

First, to test for an association between expression variability and purifying selection, we grouped genes into 10 bins of size 0.1 based on their median variability rank computed across available conditions (per species) and compared the *ω_0_*distribution across bins (**Figure 4**; **Figure S28**). Second, to check for an association between variability and positive selection, we computed the proportion of gene - internal branches with evidence for positive selection based on a *q*-value cutoff of *q* < 0.05 on the branch-site likelihood test (**Table S4**). For this, genes that were frequently lowly, moderately, or highly variable (i.e., the same classification in at least half of available conditions) were grouped into three large categories based on the following thresholds: rank ≤ 0.20 for lowly variable genes, rank ≥ 0.80 for highly variable genes, and 0.20 < rank < 0.80 for moderately variable genes. To test for significance, we conducted a randomization test by permuting the *q*-values on the background gene list (*n* = 2000 permutations), in addition to a Pearson’s chi-squared test.

#### Organ expression specificity

##### Data preprocessing

To compute organ expression specificity within a species, we required at least 2 individuals per organ for spotted gar, and at least 1 individual per organ-sex condition for Northern pike and zebrafish. Furthermore, to make expression specificity measurements comparable between species, we considered only the set of organs with sufficient samples in every species, i.e., brain, eye, intestine, liver, heart, muscle, gills, pectoral fin, and gonads. After excluding organs with missing data (blood, kidney, skin, swim bladder), we repeated our gene filtering and normalization step starting from raw read counts (Gene filtering and normalization).

##### Expression specificity index

For each retained gene per species, organ expression specificity was measured using the tau (*τ*) index, which is defined as:

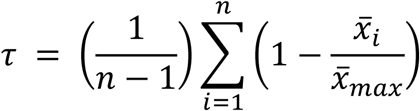

where **n** is the number of organs, *x̅_i_* is the mean expression of gene **x** in organ **i**, and *x̅_max_* is the maximum mean expression of gene **x** within an organ 1 ≤ *i* ≤ *n* (Yanai et al. 2005; Kryuchkova-Mostacci and Robinson-Rechavi 2017). The tau index gives a single measure of organ specificity per gene, ranging from broadly expressed across all organs (*τ* → 0) to specifically expressed in a single organ (*τ* → 1). For species with differentiated sexes (Northern pike and zebrafish), the ovary and testis were considered separate anatomical entities. However, for non-reproductive organs in the same species, we did not consider each sex separately.

#### Organ-biased expression and gene expression variability

Per species, genes with tau (*τ*) ≤ 0.3 were classified as broadly expressed, while genes with tau above this threshold were classified as organ-biased. (See **Figures S41-S43** for an alternative cutoff of *τ* ≤ 0.5.) For genes which are organ-biased, the ‘focal organ’ is defined as the organ in which the gene is maximally expressed.

##### Multiple pairwise comparisons

For each species (spotted gar) or species-sex (Northern pike and zebrafish), let *S*_1_ be the set of *m* organs which (a) have been used in expression specificity computations, and (b) have gene expression variability estimates (**Figure 1**):

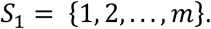

Let *S*_2_ be the set of *n* ≥ *m* organs which satisfy (a) but not (b):

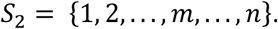

For each organ *k* ∈ *S*_1_, we compared the variability rank distribution within *k* for broadly expressed genes (*τ* ≤ 0.3) versus genes which are organ-biased (*τ* > 0.3) and with top expression in focal organ *k*. For example, given the pectoral fin as the focal organ, we compared the variability ranks of broadly expressed genes against pectoral-fin-biased genes (**Figure 5A, 5E, 5I**).

Next, still given *k*, we iterated through the set of other (non-focal) organs in *S*_1_ which are not *k*:

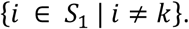

For each iteration of *i* ≠ *k*, we defined a set of *i*-biased genes, which are organ-biased and with top expression in non-focal organ *i*, but still expressed in focal organ *k*. Then, we compared the variability rank distribution within *k* for genes which are *i*-biased versus other organ-biased genes with top expression in the set *J* of (non-focal) organs in *S*_2_ which are neither *k* nor *i*:

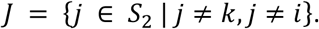

As an example, for expression variability in the pectoral fin, we compared the variability ranks of brain-biased (or liver-biased, ovary-biased, etc.) genes expressed in the pectoral fin against the variability ranks of the set of other organ-biased genes that are neither pectoral-fin-biased nor brain-biased (or liver-biased, ovary-biased, etc.), and are also expressed in the pectoral fin (**Figure 5A, 5E, 5I**).

For each pairwise comparison of variability ranks of *i*-biased versus (other-biased) *J*-biased genes, iterated for each organ *k* ∈ *S*_1_, we tested for a difference in group means using Wilcoxon-rank sum tests, and applied a Benjamini-Hochberg (‘BH’) correction for multiple testing. In addition, we calculated the effect size using Glass’s δ, using the set of *J*-biased genes as the reference group:

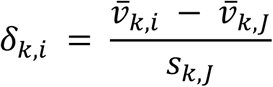

where *v̅*_*k*,*i*_ is the mean variability rank of *i*-biased genes in focal organ *k*, *v̅*_*k*,*J*_ is the mean variability rank of *J*-biased genes, and *s*_*k*,*J*_ is the sample standard deviation of the variability ranks of *J*-biased genes. We visualized the distribution of effect sizes using a heatmap, where rows correspond to focal organ *k* and the columns correspond to sets of *i*-biased genes (**Figure 5D, 5H, 5L**). We showed pairwise comparisons that are considered statistically significant after BH multiple testing correction (adjusted *p*-value < 0.05).

##### Analysis on randomized data

Given the gene x organ mean expression matrix of a species, we permuted the values of the mean expression vector per gene so that the focal organ of each gene is assigned randomly without changing tau. Since tau is unchanged, the same set of genes are classified as broadly expressed (*τ* ≤ 0.3) and organ-biased, but the subclassifications of organ-biased genes (e.g., brain-biased, liver-biased, etc.) differ. We pre-set two seeds (12345, 67890) for our sample randomizations. Multiple pairwise comparisons of expression variability ranks for subsets of organ-biased genes were performed as previously described (**Figures S59**-**S61**).

##### Analysis on nonzero expression matrix

To control for differences in number of genes expressed among organs (**Figure S54**), we repeated the same analysis (i.e., gene expression normalization, expression variability estimation, comparisons of expression variability between subsets of genes) as previously described, considering only genes that have nonzero counts across all samples within a species (**Figure S55**).

#### Organ-biased expression and coding-sequence selection

Per species, we retrieved expression specificity measurements for each gene (Organ expression specificity) and filtered selection statistics for each gene - internal branch (Coding-sequence selection analysis), as described in previous sections. We annotated each gene from the Selectome dataset with its top (focal) organ of expression (i.e., organ bias). If a gene has *τ* ≤ 0.3, its organ bias is labeled ‘broad’. We compared the *ω_0_*distribution of genes across organ bias categories to check for any association between focal organ and strength of purifying selection at the coding-sequence level (**Figure S50**).

#### Promoter sequence analysis

To test whether organ-biased promoters can be distinguished from random sequences and from each other based on variability patterns, we trained gapped *k*-mer SVM models (gkmSVM) on promoter sequences. Analyses were first performed on each organ-biased promoter set, using subsamples of up to 1,000 genes to avoid sample-size bias **(Figure S62)**. Organ-biased promoter sets were also tested against promoters from the other organ-biased sets to evaluate whether they could be distinguished from one another **(Figure S49)**. We then grouped zebrafish organ-biased genes into two categories: those typically lowly variable in other organs (brain, ovary, testis, eye) and those highly variable in the others (liver, pectoral fin, gills, intestine), with up to 5,000 genes per set. In addition, three random gene sets were generated **(Figure S64)**. Promoter regions were defined as the 2-kb sequence upstream of the canonical TSS (Ensembl Biomart release 114) (Dyer et al. 2025), and exonic regions were removed with BEDTools subtract (Quinlan and Hall 2010).

For each promoter set, a matched negative set of genomic sequences was drawn from the DanRer11 assembly, matched for length, repeat content, and GC content. This was done by building a BSgenome object with UCSC annotations (Pagès 2025) and sampling with the genNullSeqs function from the gkmSVM R package (Ghandi et al. 2016). Models were trained with gkmtrain (LS-GKM) (Lee 2016) using default parameters, and performance was assessed by 5-fold cross-validation using the -x option and receiver operating characteristic (ROC) curves.

## Acknowledgements

We would like to thank Allyse Ferrara and Quenton Fontenot (Nicholls State University, USA) for providing gar embryos and members of the Braasch Lab for help with gar husbandry. We thank the four reviewers for their comments which improved this study. We acknowledge having used ChatGPT (GPT-4) (OpenAI, 2023, https://chat.openai.com) to improve the clarity of the manuscript. Any errors or omissions are our own. M.R.-R., I.B., Y.G., and J.H.P. conceived the experimental design. C.F.B. and M.R.-R. conceived the design of data analysis. A.W.T., B.L.R., and I.B. sourced and dissected the spotted gar samples; C.A.W. and J.H.P., the zebrafish samples; J.B. and Y.G., the Northern pike samples. C.A. performed RNA extraction. S.M. provided Selectome data for positive selection analysis. A.L. and A.T. performed the promoter sequence analysis. C.F.B. performed the data analysis and generated the figures with input from M.R.-R. C.F.B. wrote the original draft of the manuscript with input from M.R.-R; C.F.B., A.L., A.W.T., J.B., I.B., Y.G., J.H.P., and M.R.-R wrote the final version of the manuscript. C.F.B., A.L., A.W.T., J.B., I.B., Y.G., J.H.P., and M.R.-R. contributed to the interpretation and discussion of results.

## Data Access

Demultiplexed FASTQ files generated via BRB-seq for zebrafish (192 samples), Northern pike (252 samples), and spotted gar (177 samples) are available on the NCBI Sequence Read Archive (SRA) under accession number PRJNA1195359.

Data analysis scripts are available on GitHub: https://github.com/mrrlab/fish-variability-across-organs

Additionally, scripts and supplementary data files are available on Zenodo: https://doi.org/10.5281/zenodo.14063787

