## Supplementary Information for "Inter-individual gene expression variability implies stable regulation of brain-biased genes across organs in three ray-finned fishes"

### Supplementary Results

**Bimodality test**

One limitation of the variability rank metric is that it does not describe the modality of the underlying expression distribution. Overdispersion may be due to either a broader range of expression centered at a single mean (i.e., a unimodal distribution) or the presence of two or more expression peaks (a multimodal distribution), the latter of which may be indicative of hidden substructures within the population (Mar 2019). To investigate this possibility, we assessed whether more bimodal expression profiles can be observed for sets of increasingly more variable genes. For each condition with at least 10 replicates, we grouped genes into variability rank bins and plotted the kernel density distribution of expression level z-scores to check visually for bimodality in each bin (**Figure S18A; Figure S19A**). Second, for each gene, we computed the bimodality index (BI), a metric ranging from $[0,\infty)$ that quantifies bimodality based on a two-component Gaussian mixture model, with larger values corresponding to a stronger bimodality signal (Wang et al. 2009). For some conditions (e.g., female Northern pike brain) we observed an increase in bimodality for the most variable genes (**Figure S18B**); however, this was not always the case (e.g., Northern pike pectoral fin) (**Figure S20**). (See **Figure S21** and **Figure S22** for the corresponding analyses on zebrafish and spotted gar, respectively.) The BI cutoff in which strong bimodality can be observed is sample-size dependent (Wang et al. 2009). Since it is likely that there is low power to detect bimodality at the given sample sizes, we tested for sensitivity by simulating a highly bimodal expression distribution that combines the same number of $n/2$ replicates from two different conditions (e.g., female Northern pike brain and ovary). Compared to observed data of the same total sample size $n$, we recovered more strongly bimodal expression z-score profiles (**Figure S19B**, **Figure S23**) and a higher BI distribution (**Figure S24**). From this, we found that the effect size is small even for conditions in which we detect some evidence for bimodality for highly variable genes. Overall, we find weak evidence for bimodality in our dataset, only in some conditions, and only at very high variability ranks; thus, we infer that higher variability is likely primarily driven by a wider range of expression centered around a single peak.

### Supplementary Methods

#### Other metrics to estimate gene expression variability

To estimate gene expression variability within a condition, several dispersion statistics were computed in a jackknife resampling ('leave-one-out') procedure. Apart from the local (residual) coefficient of variation, the following expression dispersion metrics were also calculated for each *n* - 1 subset:

*Adjusted standard deviation*

The adjusted standard deviation ('adjusted SD') is the ratio of observed SD to predicted SD (Liu et al. 2020). Using the 'loess' function from the *stats* package in R, we fit a LOESS regression curve (with span = 0.6 and default parameters) to model expression SD as a function of mean expression. Given this metric, genes with adjusted SD > 1 are overdispersed, whereas genes with 0 < adjusted SD < 1 are underdispersed relative to the expected value.

*Residual standard deviation*

We computed the $log_2$-transformed standard deviation for each gene and fit a LOESS regression with the same parameters to model $log_2$(SD) as a function of mean expression. The difference between observed and predicted $log_2$(SD) is the residual $log_2$(SD) ('residual SD', for brevity) after adjusting for mean expression. Given this metric, genes with residual SD > 0 are overdispersed, whereas genes with residual SD < 0 are underdispersed relative to the model prediction.

For each gene per condition, we computed the mean for each expression dispersion statistic over all *n* - 1 subsets.

*Residual median absolute deviation (MAD) without jackknife resampling*

We verified our analysis by applying a different variability metric based on the residual MAD, and without applying jackknife resampling to the dataset (**Figures S56**-**S58**). A LOESS regression curve with span = 0.6 and default parameters was used to model MAD as a function of mean expression.

#### Bimodality test

For each organ (for spotted gar) or organ-sex condition (for Northern pike and zebrafish) with at least 10 replicates, we grouped protein-coding genes into 10 variability rank bins of size 0.1 and plotted the kernel density distribution of expression level z-scores to qualitatively check for strong (i.e., visually observable) bimodality in each bin. The expression z-score ($z$) was computed from the normalized, log-transformed expression matrix ($log_2$ TMM-CPM) and defined as:

$$z = \frac{x_i - \mu}{\sigma}$$

where $x_i$ is the observed expression of a gene in replicate $i$, $\mu$ is the mean expression of a gene across all replicates $n$ within a condition, and $\sigma$ is the standard deviation.

Per gene within a condition, we computed the bimodality index (BI) proposed by Wang et al. (2009) and implemented in the *BimodalIndex* (1.1.9) package (Wang et al. 2009). BI is a continuous metric that quantifies the strength of bimodality by fitting the expression data to a two-component Gaussian mixture model with a common standard deviation. We used expression level z-scores as input to compute BI for each gene. Higher BI values indicate stronger bimodality based on the distribution of samples between the two groups or the standardized distance between groups. The BI cutoff in which strong bimodality can be observed visually is dependent on sample size, with higher BI values necessary at smaller sample sizes (Wang et al. 2009). To estimate the upper limit of BI that can be detected for a given sample size $n$ = 10, we simulated a highly bimodal expression distribution by combining 5 randomly selected replicates from female Northern pike brain and 5 replicates from female Northern pike ovary. We performed gene filtering and normalization on the raw count matrix of this pseudo-condition (Gene filtering and normalization) and computed jackknifed estimates of expression variability rank (Gene expression variability estimation), both as described previously, and retained only protein-coding genes. We computed the bimodality index for each gene given this pseudo-condition and compared the BI distribution with that based on observed data from 10 female Northern pike brain samples (**Figure S18**). For both observed and mixed data, we plotted the kernel density distribution of expression level z-scores of genes sorted by variability rank (**Figure S19**) as well as bimodality index **(Figure S23)** to check for onset of strongly bimodal expression profiles. In addition, we compared the BI distribution of genes binned by variability rank (**Figure S24**).

**Weighted gene co-expression network analysis (WGCNA)**

Modules of co-expressed genes were identified via weighted gene co-expression network analysis using the *WGCNA* (1.74) package (Langfelder and Horvath 2008). To ensure reliable network inference, only organ or organ-sex conditions with at least 10 biological replicates were included. For each condition, subsets of the normalized gene expression matrix ($\log_2$ TMM-CPM) were formed by grouping genes based on organ bias category (“broad”, “focal”, “other”) and variability rank bins of width 0.2. For each expression matrix subset, gene co-expression networks were constructed using blockwiseModules(), using a signed correlation network with a soft-thresholding power of 10 and minimum module size of 30 genes. To quantify modular structure, we calculated the proportion of genes assigned to the dummy (gray) module, which represents genes not assigned to a co-expression module (**Figure S49**).

To assess whether observed co-expression structure exceeds that expected from random gene-gene associations, we applied WGCNA to identify pseudo-modules from randomized data. For each expression matrix subset, a randomized dataset was generated by rowwise permutation of the gene x sample matrix. This preserves the mean and variance of each gene’s expression profile while removing the gene-by-gene covariance structure driving co-expression patterns. Each permuted dataset was then analyzed using the same WGCNA pipeline as the observed data, and modularity metrics were compared between observed and randomized datasets.

### Supplementary Tables

**Table S1. Sequencing information for each project**

| Project | Species | Reference Genome | Multiplexed Libraries | Demultiplexed Samples | Sequencer | R1 (bp) | R2 (bp) |
|---|---|---|---|---|---|---|---|
| AG0012 | Northern pike (*E. lucius*) | Eluc_v4 (Ensembl 101) | 5 | 258 | Illumina NovaSeq 6000 | 25 | 80 |
| AG0012 | Spotted gar (*L. oculatus*) | LepOcu1 (Ensembl 103) | 1 | 24 | Illumina NovaSeq 6000 | 25 | 80 |
| AMP0027 | Spotted gar (*L. oculatus*) | LepOcu1 (Ensembl 103) | 2 | 155 | Illumina NextSeq 550 | 21 | 55 |
| AMP0020 | Zebrafish (*D. rerio*) | GRCz11 (Ensembl 101) | 3 | 192 | Illumina NextSeq 550 | 21 | 55 |

**Note:** 'Demultiplexed samples' refer to number of samples prior to quality filtering. See **Supplementary Data** for sample metadata.

**Table S2. Sample filtering criteria for organ expression specificity analysis**

| Species | Filtering criteria | | | Sample drop-off | | |
|---|---|---|---|---|---|---|
| | Minimum uniquely mapped reads | Minimum detected genes | Minimum replicates per condition | Samples | QC 1: Sequencing quality | QC 2: Within-organ correlation |
| Spotted gar (*L. oculatus*) | 200,000 | 6,000 | 2 | 177* | 106 | 104 |
| Northern pike (*E. lucius*) | 300,000 | 10,000 | 1 | 252* | 191 | 174 |
| Zebrafish (*D. rerio*) | 500,000 | 10,000 | 1 | 192 | 186 | 182 |

**Note:** We required a minimum of 1 individual per organ-sex or 2 individuals per organ (as in spotted gar) to compute expression specificity of each gene.
*Sequenced samples with inconsistent metadata (spotted gar: *n* = 2; Northern pike: *n* = 6) were removed prior to downstream processing.

**Table S3. Sample filtering criteria for expression variability analysis**

| Species | Filtering criteria | | | Sample drop-off | | |
|---|---|---|---|---|---|---|
| | Minimum uniquely mapped reads | Minimum detected genes | Minimum replicates per condition | Demultiplexed Samples | QC 1: Sequencing quality | QC 2: Within-organ correlation |
| Spotted gar (*L. oculatus*) | 200,000 | 6,000 | 4 | 177* | 106 | 104 |
| Northern pike (*E. lucius*) | 300,000 | 10,000 | 4 | 252* | 183 | 166 |
| Zebrafish (*D. rerio*) | 500,000 | 10,000 | 4 | 192 | 178 | 177 |

**Note:** We required a minimum of 4 individuals per organ- or organ-sex condition to estimate expression variability per gene.

**Table S4. Positive selection analysis**

| Species | Expression variability category | No evidence for positive selection ($q$-value > 0.05) | Evidence for positive selection ($q$-value < 0.05)** | Percentage of branches with $q$-value < 0.05 (%) | Enrichment | $p$-value*** |
|---|---|---|---|---|---|---|
| Spotted gar (*L. oculatus*) | low | 75 | 9 | 10.7 | 0.89 | 0.746 |
| | moderate | 3878 | 529 | 12.0 | 1.00 | 0.825 |
| | high | 267 | 44 | 14.1 | 1.17 | 0.257 |
| | background* | 4741 | 650 | 12.1 | | |
| Northern pike (*E. lucius*) | low | 313 | 12 | 3.7 | 0.37 | $< 5 \times 10^{-4}$ |
| | moderate | 22910 | 2513 | 9.9 | 0.98 | 0.004 |
| | high | 1060 | 186 | 14.9 | 1.48 | $< 5 \times 10^{-4}$ |
| | background | 25802 | 2890 | 10.1 | | |
| Zebrafish (*D. rerio*) | low | 1090 | 53 | 4.6 | 0.55 | $< 5 \times 10^{-4}$ |
| | moderate | 31477 | 2923 | 8.5 | 1.01 | 0.110 |
| | high | 2443 | 214 | 8.1 | 0.96 | 0.520 |
| | background | 39018 | 3575 | 8.4 | | |

**Note:**

*The background set also includes genes which have not been classified as frequently lowly, moderately, or highly variable, based on filtering criteria (Materials and Methods).

**Number of gene - internal branch entries with statistically significant ($q$-value < 0.05) evidence for sites under positive selection based on the branch-site likelihood-ratio test. Data retrieved from Selectome (Proux et al. 2009; Moretti et al. 2014).

Internal branches:

(A) For spotted gar: Euteleostomi, Actinopterygii, Neopterygii

(B) For Northern pike: Euteleostomi, Actinopterygii, Neopterygii, Osteoglossocephalai, Clupeocephala, Euteleosteomorpha, Protacanthopterygii

(C) For zebrafish: Euteleostomi, Actinopterygii, Neopterygii, Osteoglossocephalai, Clupeocephala, Otomorpha, Otophysi

***Two-tailed $p$-values based on permutation of $q$-values on the background gene set ($n$ = 2000 permutations)

### Supplementary Figures

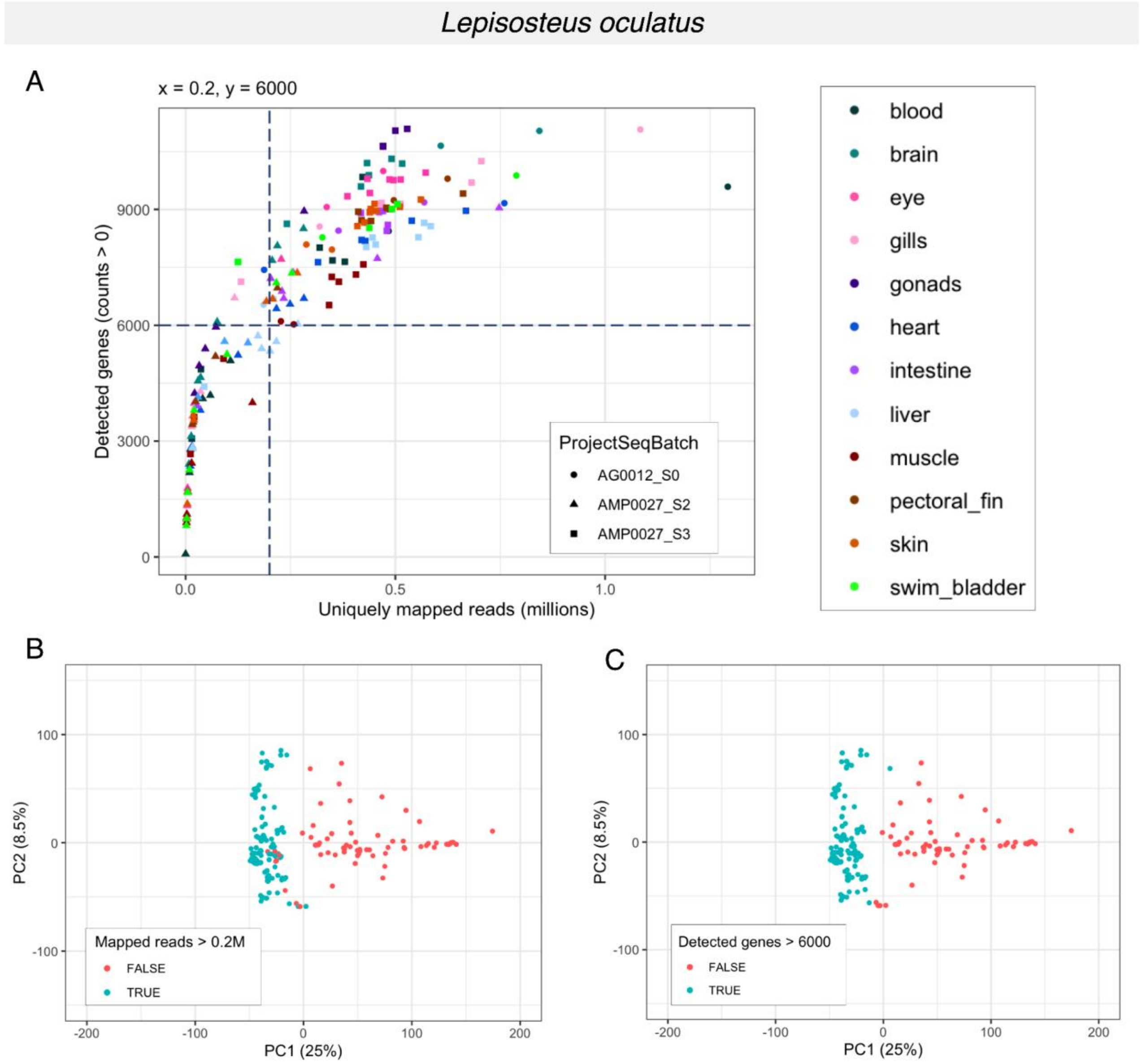


**Figure S1. Removal of spotted gar samples with low sequencing quality. (A)** Relationship between number of detected genes and uniquely mapped reads. Color and shape indicate organ and sequencing batch, respectively. Because of differences in gene number and annotation quality between species, species-specific cutoffs were applied to remove low quality samples. Dashed vertical and horizontal lines indicate minimum thresholds for number of uniquely mapped reads (*x*) and detected genes (*y*), respectively. For spotted gar: spotted gar: $x > 0.2$ million uniquely mapped reads, $y > 6000$ detected genes. **(B-C)** Principal component analysis (PCA) of organ gene expression profiles. Expression levels are log-transformed, TMM-normalized counts per million ($\log_2$ TMM-CPM), normalized across all conditions. Percentage of variance for the first and second principal components are indicated on the x- and y-axes, respectively. Biplots are colored by species-specific cutoffs for **(B)** uniquely mapped reads and **(C)** detected genes. Only samples above the thresholds (blue, 'TRUE') were considered for downstream analyses; samples that did not meet either criteria (red, 'FALSE') were discarded. For spotted gar, the number of samples before ($n_1$) and after ($n_2$) the first quality filtering step are $n_1 = 177$ and $n_2 = 106$, respectively.

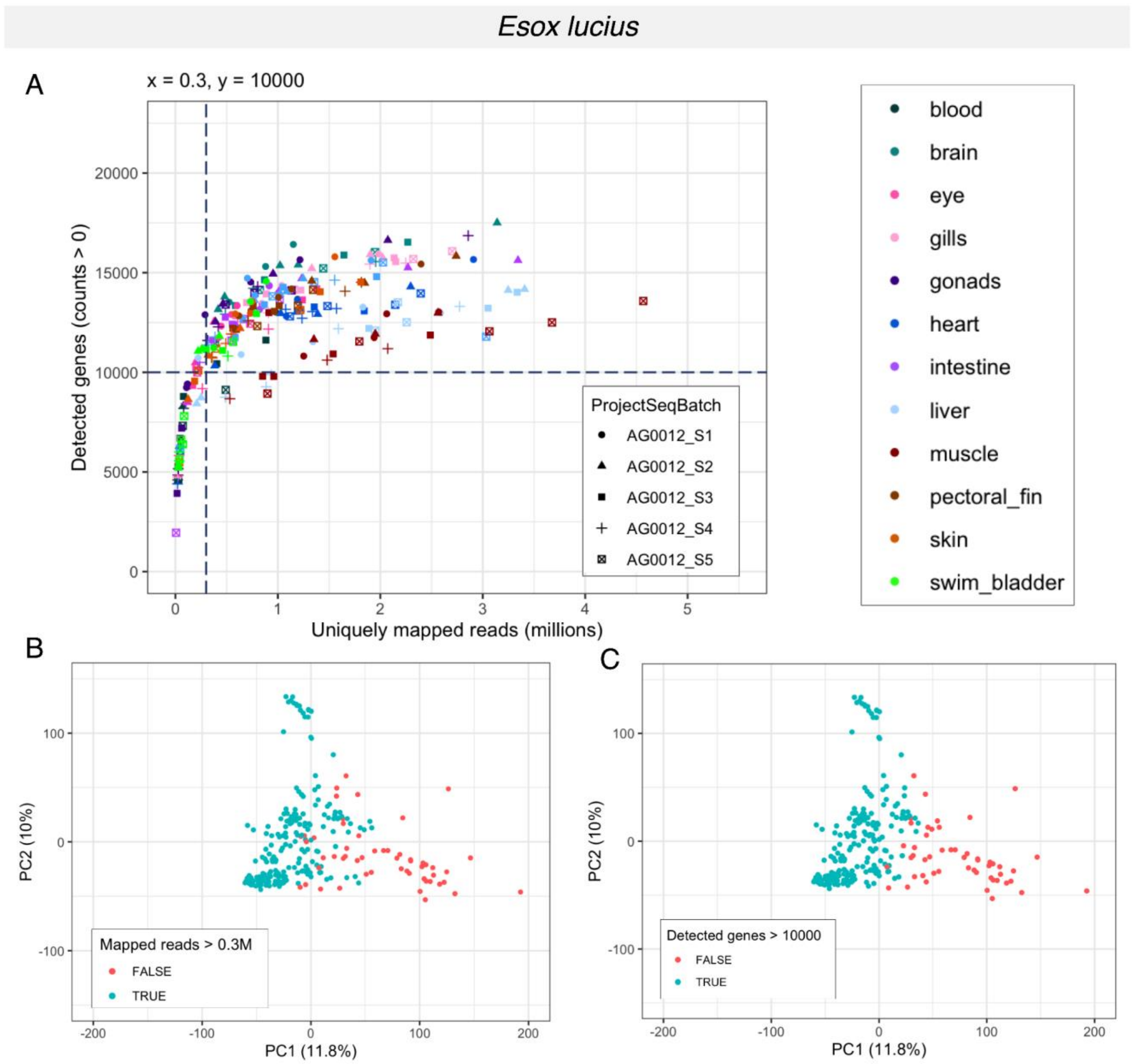


**Figure S2. Removal of Northern pike samples with low sequencing quality. (A)** Relationship between number of detected genes and uniquely mapped reads. Color and shape indicate organ and sequencing batch, respectively. Because of differences in gene number and annotation quality between species, species-specific cutoffs were applied to remove low quality samples. Dashed vertical and horizontal lines indicate minimum thresholds for number of uniquely mapped reads (*x*) and detected genes (*y*), respectively. For Northern pike: *x* > 0.3 million uniquely mapped reads, *y* > 10000 detected genes. **(B-C)** Principal component analysis (PCA) of organ gene expression profiles. Expression levels are log-transformed, TMM-normalized counts per million ($\log_2$ TMM-CPM), normalized across all conditions. Percentage of variance for the first and second principal components are indicated on the x- and y-axes, respectively. Biplots are colored by species-specific cutoffs for **(B)** uniquely mapped reads and **(C)** detected genes. Only samples above the thresholds (blue, 'TRUE') were considered for downstream analyses; samples that did not meet either criteria (red, 'FALSE') were discarded. For Northern pike, the number of samples before ($n_1$) and after ($n_2$) the first quality filtering step are $n_1$ = 252 and $n_2$ = 191, respectively.

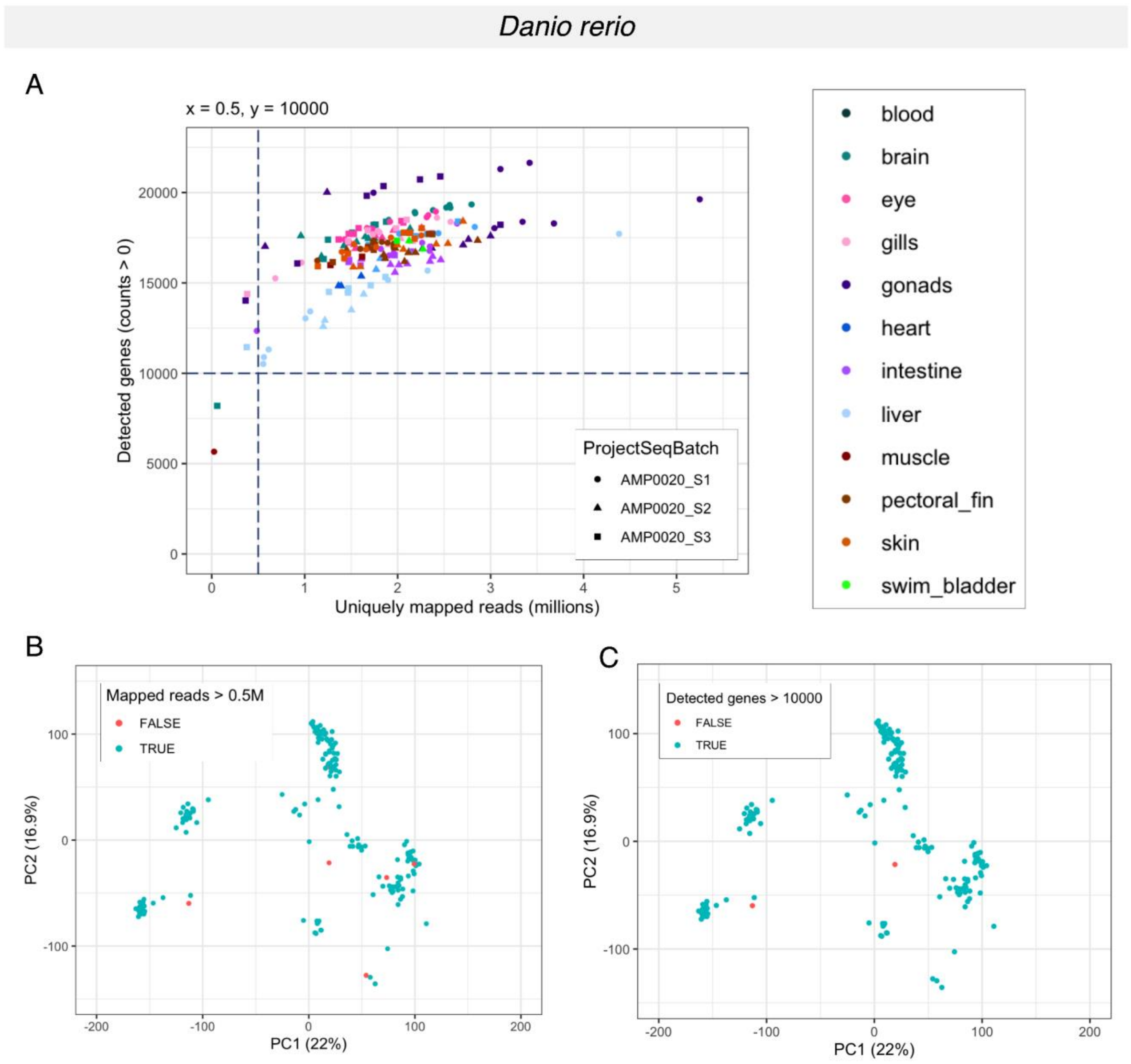


**Figure S3. Removal of zebrafish samples with low sequencing quality. (A)** Relationship between number of detected genes and uniquely mapped reads. Color and shape indicate organ and sequencing batch, respectively. Because of differences in gene number and annotation quality between species, species-specific cutoffs were applied to remove low quality samples. Dashed vertical and horizontal lines indicate minimum thresholds for number of uniquely mapped reads (*x*) and detected genes (*y*), respectively. For zebrafish: $x$ > 0.5 million uniquely mapped reads, $y$ > 10000 detected genes. **(B-C)** Principal component analysis (PCA) of organ gene expression profiles. Expression levels are log-transformed, TMM-normalized counts per million ($\log_2$ TMM-CPM), normalized across all conditions. Percentage of variance for the first and second principal components are indicated on the x- and y-axes, respectively. Biplots are colored by species-specific cutoffs for **(B)** uniquely mapped reads and **(C)** detected genes. Only samples above the thresholds (blue, 'TRUE') were considered for downstream analyses; samples that did not meet either criteria (red, 'FALSE') were discarded. For zebrafish, the number of samples before ($n_1$) and after ($n_2$) the first quality filtering step are $n_1$ = 192 and $n_2$ = 186, respectively.

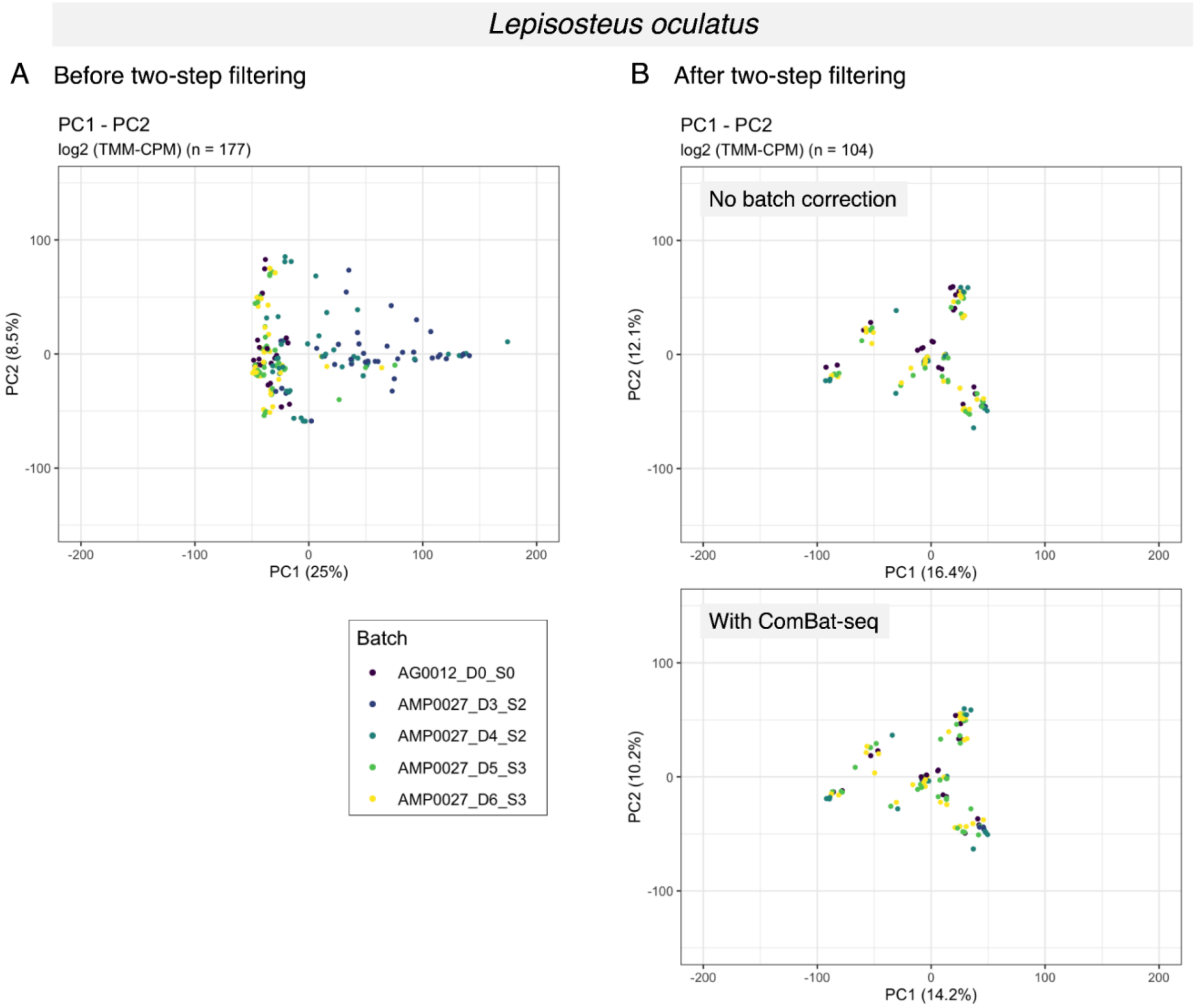


**Figure S4. Principal component analysis (PCA) of spotted gar organ transcriptomes colored by extraction and sequencing batch.** Expression data is in the form of $log_2$ TMM-CPM. The percentage of variance for principal components 1 and 2 is indicated on the respective axes. **(A)** All samples included before two-step sample filtering ($n$ = 177). **(B)** Samples remaining after two-step sample filtering and with at least 4 replicates per organ ($n$ = 104). Top right: No batch correction was applied to the raw count matrix prior to gene filtering and normalization. Bottom right: A batch-corrected count matrix was generated prior to gene filtering and normalization using *ComBat-seq* (Zhang et al. 2020), with sequence and extraction batch as the batch variable and organ as the grouping variable.

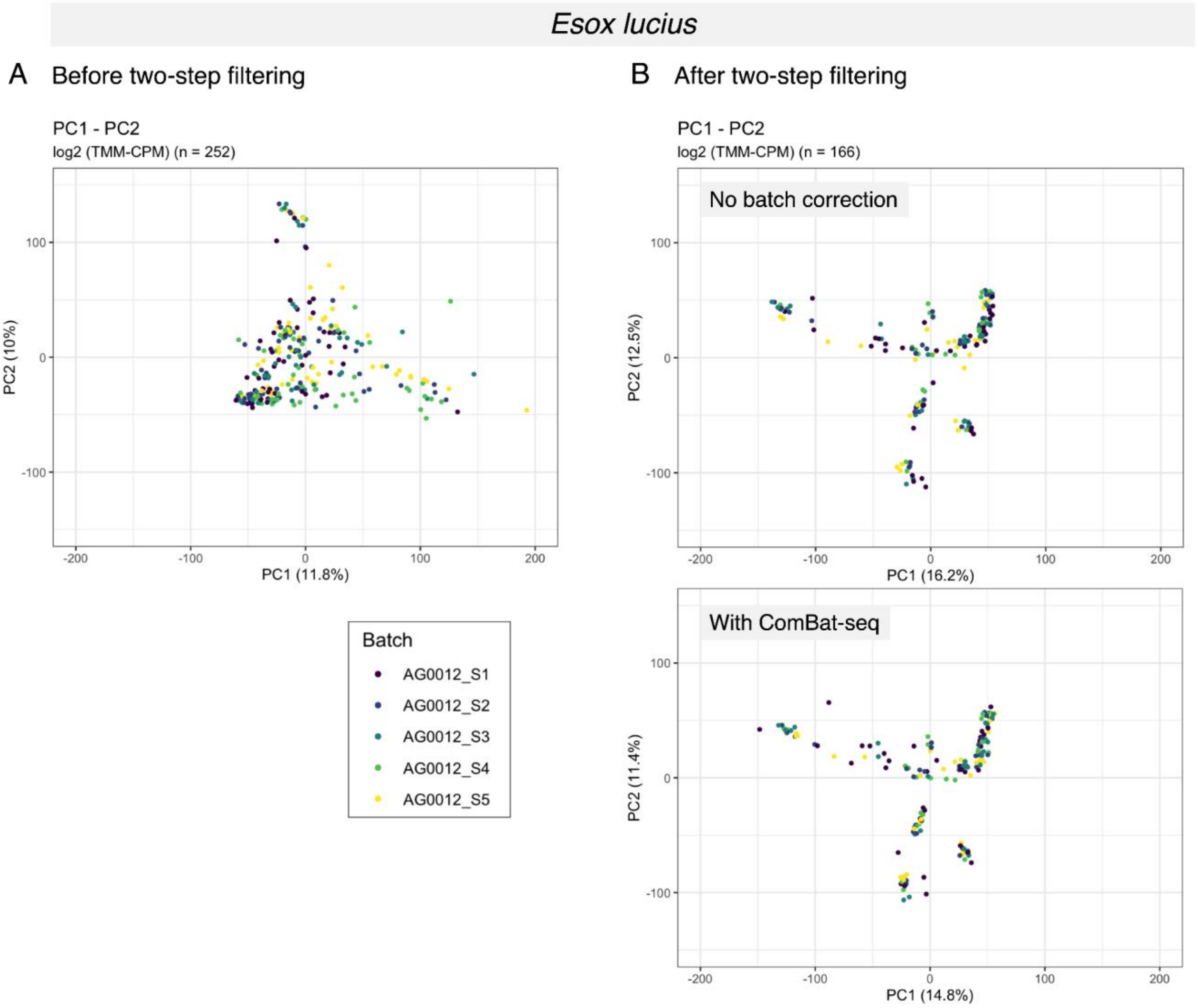


**Figure S5. Principal component analysis (PCA) of Northern pike organ transcriptomes colored by extraction and sequencing batch.** Expression data is in the form of $log_2$ TMM-CPM. The percentage of variance for principal components 1 and 2 is indicated on the respective axes. **(A)** All samples included before two-step sample filtering (*n* = 252). **(B)** Samples remaining after two-step sample filtering and with at least 4 replicates per organ (*n* = 166). Top right: No batch correction was applied to the raw count matrix prior to gene filtering and normalization. Bottom right: A batch-corrected count matrix was generated prior to gene filtering and normalization using *ComBat-seq* (Zhang et al. 2020), with sequence and extraction batch as the batch variable and organ-sex as the grouping variable.

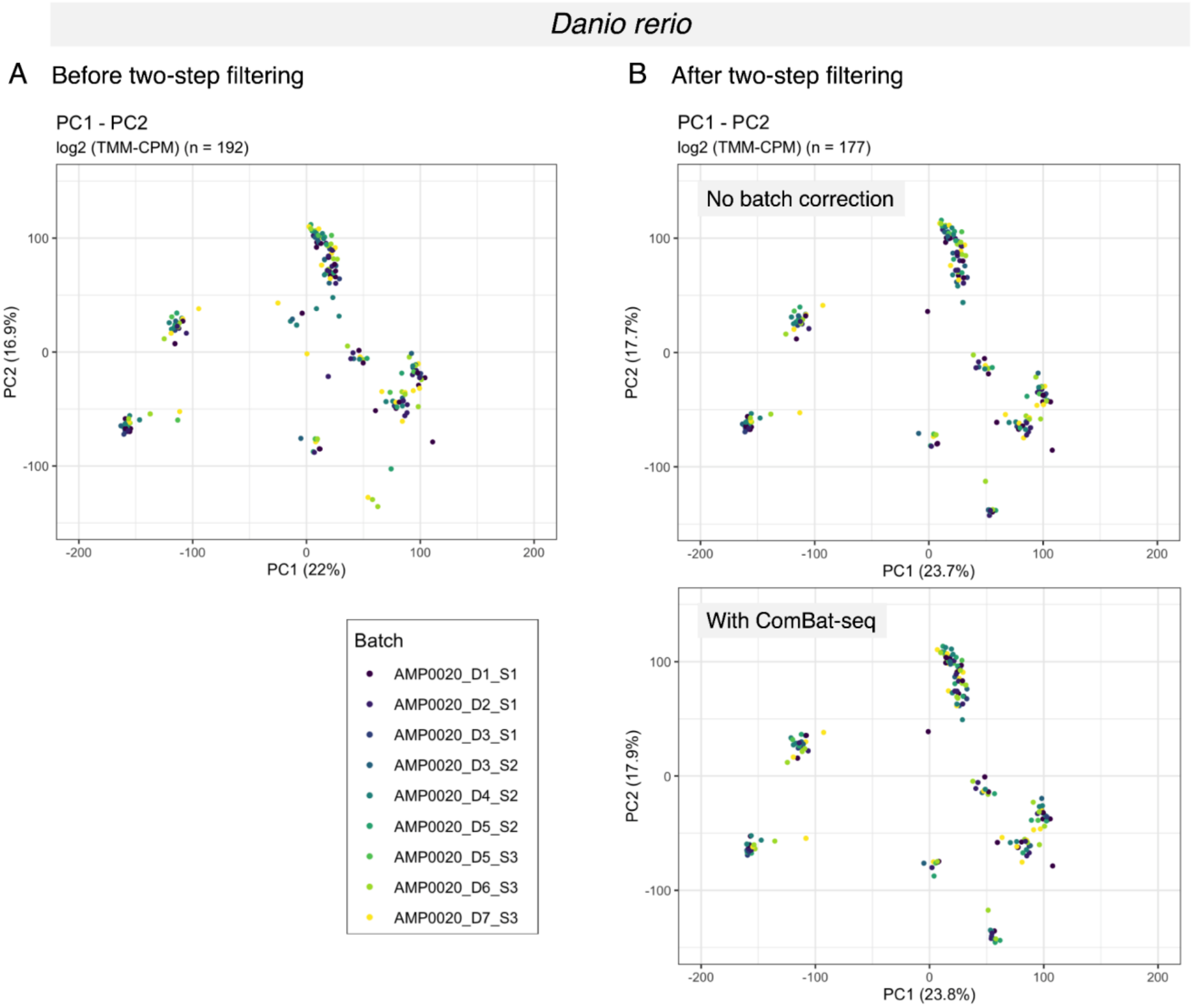


**Figure S6. Principal component analysis (PCA) of zebrafish organ transcriptomes colored by extraction and sequencing batch.** Expression data is in the form of $log_2$ TMM-CPM. The percentage of variance for principal components 1 and 2 is indicated on the respective axes. **(A)** All samples included before two-step sample filtering ($n$ = 192). **(B)** Samples remaining after two-step sample filtering and with at least 4 replicates per organ ($n$ = 177). Top right: No batch correction was applied to the raw count matrix prior to gene filtering and normalization. Bottom right: A batch-corrected count matrix was generated prior to gene filtering and normalization using *ComBat-seq* (Zhang et al. 2020), with sequence and extraction batch as the batch variable and organ-sex as the grouping variable.

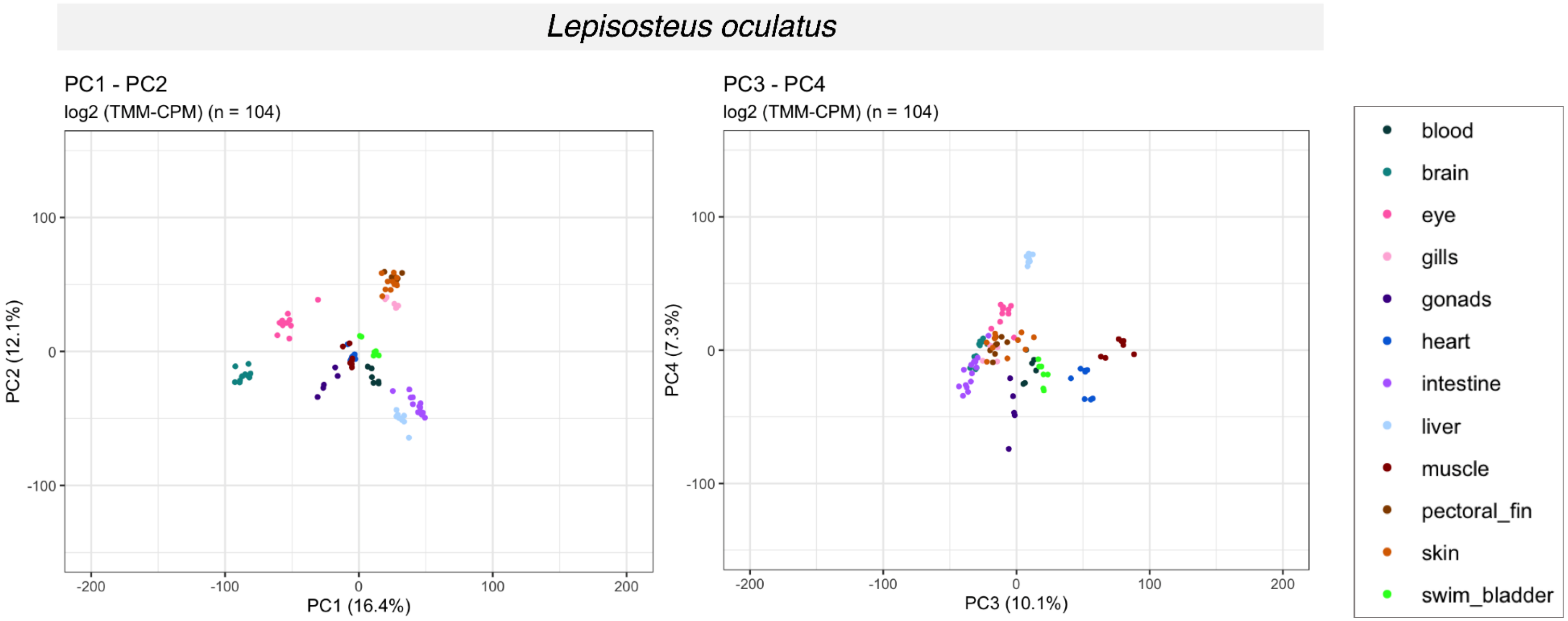


**Figure S7. Principal component analysis (PCA) of spotted gar organ transcriptomes after two-step sample filtering.** Only conditions with at least 4 replicates are included (*n* = 104 samples). Left: PCA biplot of the first and second principal components. Right column: PCA biplot of the third and fourth principal components. The percentage of variance for each component is indicated on the respective axis. Expression data is in the form of $\log_2$ TMM-CPM, normalized across all conditions.

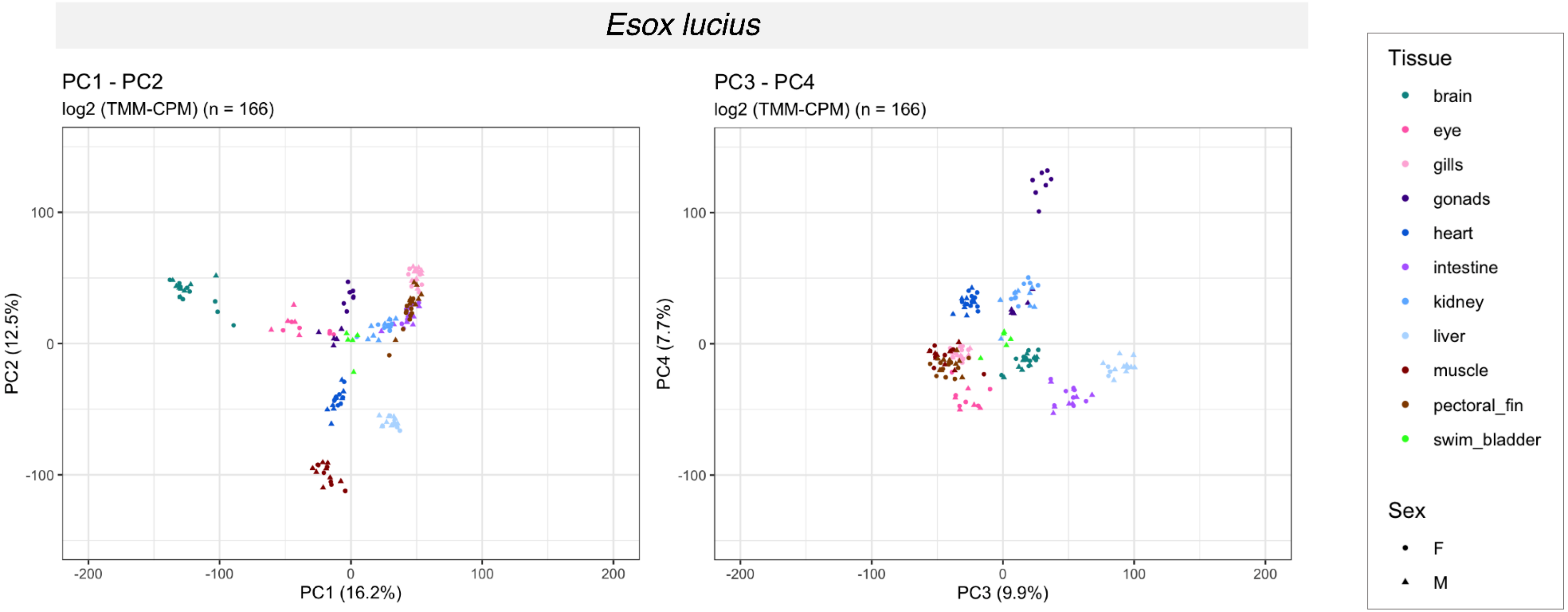


**Figure S8. Principal component analysis (PCA) of Northern pike organ-sex transcriptomes after two-step sample filtering.** Only conditions with at least 4 replicates are included (*n* = 166 samples). Left: PCA biplot of the first and second principal components. Right column: PCA biplot of the third and fourth principal components. The percentage of variance for each component is indicated on the respective axis. Expression data is in the form of $log_2$ TMM-CPM, normalized across all conditions.

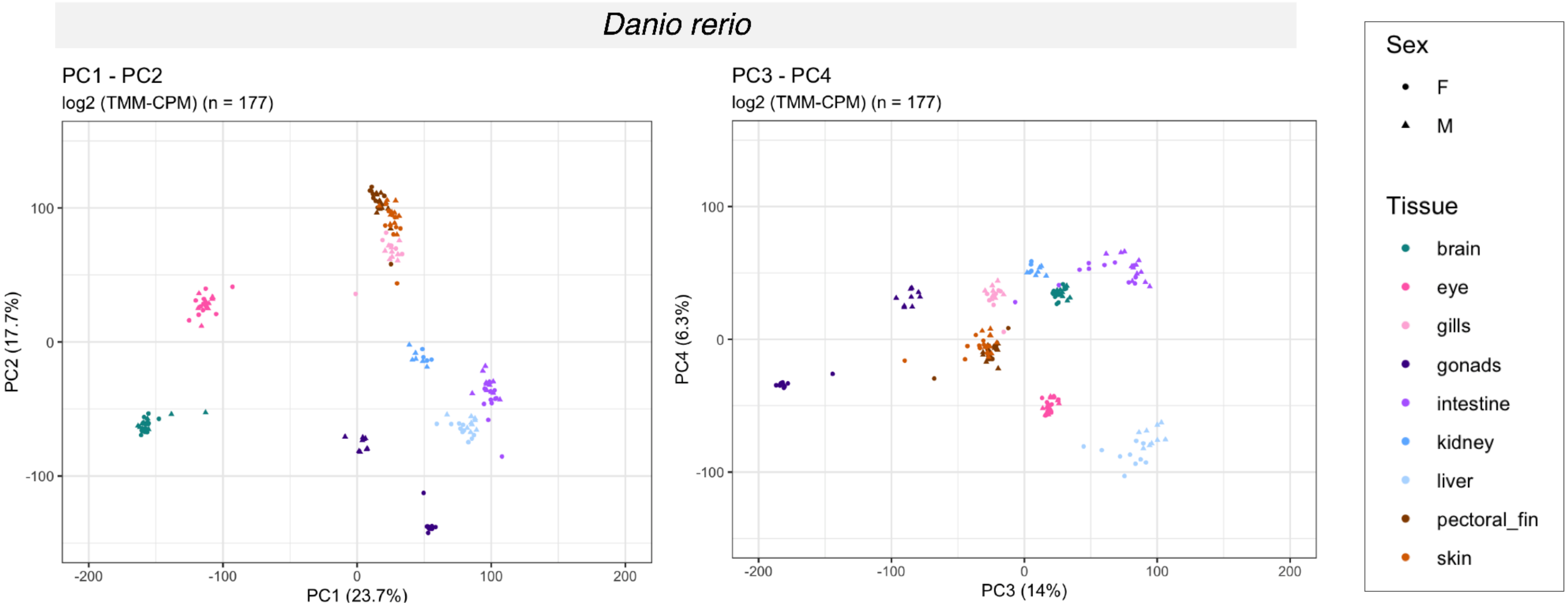


221 **Figure S9. Principal component analysis (PCA) of zebrafish organ-sex transcriptomes after two-step sample filtering.** Only conditions with at least 4 replicates are included (*n* = 177 samples). Left: PCA biplot of the first and second principal components. Right column: PCA biplot of the third and fourth principal components. The percentage of variance for each component is indicated on the respective axis. Expression data is in the form of $log_2$ TMM-CPM, normalized across all conditions.

A

| | blood | | brain | | eye | | gills | | gonads | | heart | | intestine | | kidney | | liver | | muscle | | pectoral fin | | skin | | swim bladder | | Σ | |
|---|---|---|---|---|---|---|---|---|---|---|---|---|---|---|---|---|---|---|---|---|---|---|---|---|---|---|---|---|
| | ♀ | ♂ | ♀ | ♂ | ♀ | ♂ | ♀ | ♂ | ♀ | ♂ | ♀ | ♂ | ♀ | ♂ | ♀ | ♂ | ♀ | ♂ | ♀ | ♂ | ♀ | ♂ | ♀ | ♂ | ♀ | ♂ | ♀ | ♂ |
| *D. rerio* | | | 13 | 11 | 11 | 11 | 7 | 9 | 11 | 9 | 1 | 2 | 11 | 12 | 4 | 6 | 10 | 9 | 1 | 1 | 10 | 12 | 10 | 11 | | | 89 | 93 |
| *E. lucius* | 2 | 3 | 10 | 9 | 6 | 5 | 9 | 9 | 7 | 5 | 10 | 9 | 8 | 8 | 9 | 8 | 4 | 10 | 7 | 8 | 10 | 10 | | | 3 | 5 | 85 | 89 |
| *L. oculatus* | 6 | | 11 | | 10 | | 6 | | 5 | | 9 | | 15 | | | | 8 | | 7 | | 8 | | 12 | | 7 | | 104 | |

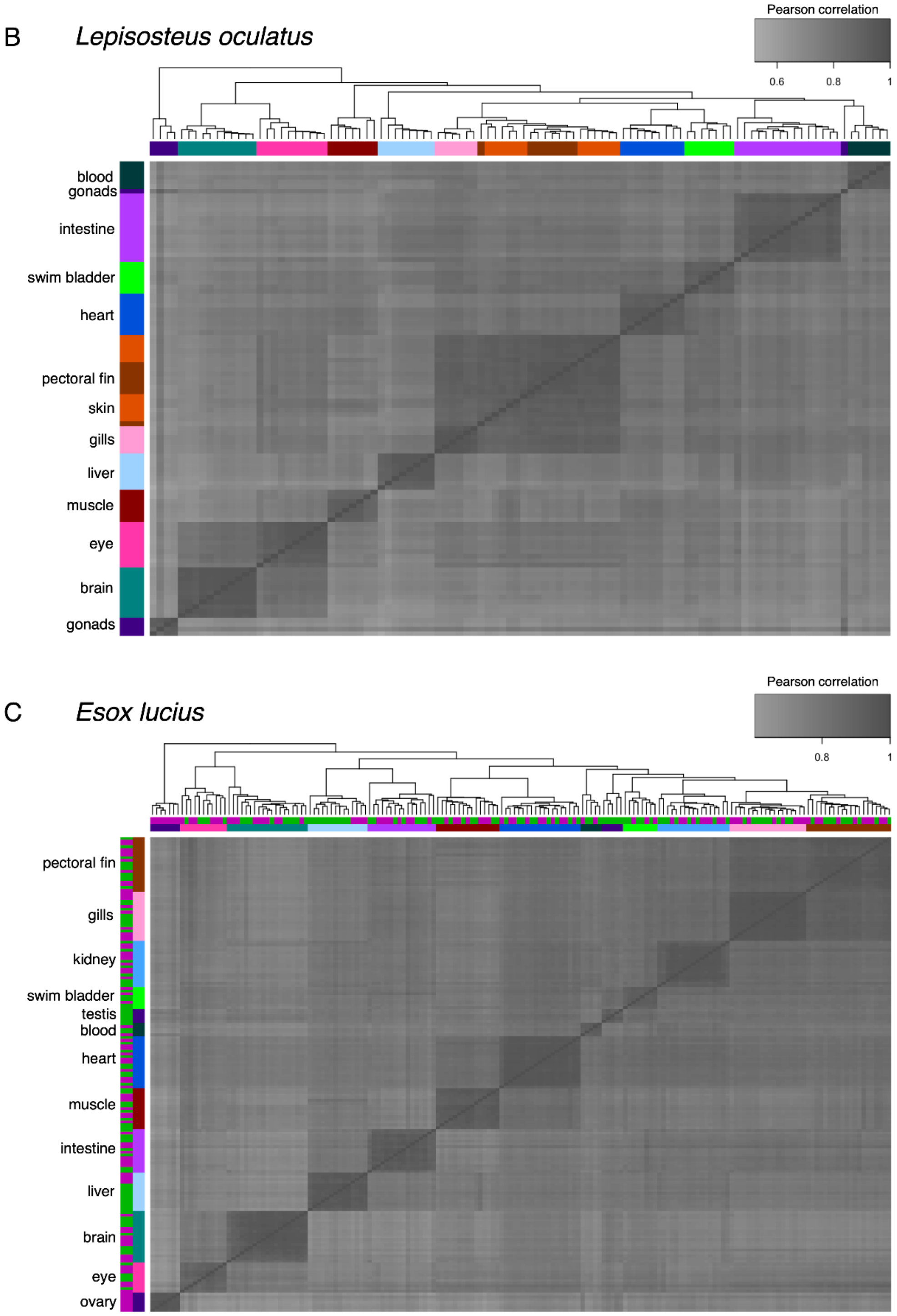


**Figure S10. Hierarchical clustering of organ transcriptomes after two-step sample filtering. (A)** Summary of BRB-seq dataset. Table shows number of sampled individuals per organ or organ-sex condition (columns) in each species (rows). Conditions with fewer than 4 replicates (light gray) were excluded from expression variability analyses. Also shown in **Figure 1B**. **(B-C)** Cluster heatmaps for **(B)** spotted gar and **(C)** Northern pike. Pairwise distances between samples are based on Pearson's correlation of expression levels ($log_2$ TMM-CPM, normalized across all conditions within each species). Only protein-coding genes were considered after expression normalization (spotted gar: $n$ = 11873 genes, Northern pike: $n$ = 17297). The color keys for organ (B and C) and sex (C only, outer color bars) are shown in (A). See **Figure 1C** for the corresponding heatmap in zebrafish.

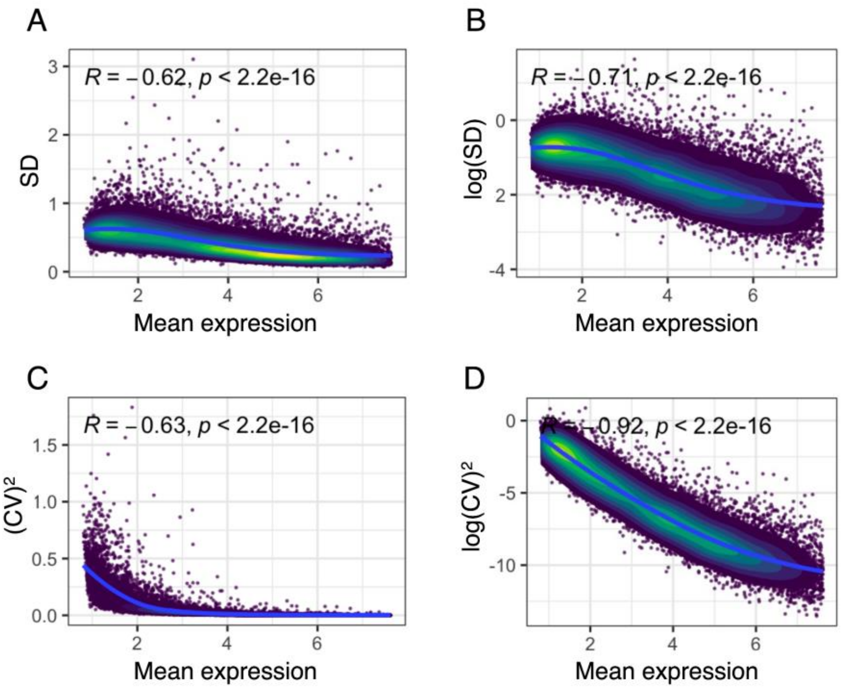


**Figure S11. Standard statistical measures of expression dispersion are strongly inversely correlated with expression level.** Data from female zebrafish brain (*n* = 13 samples). Mean expression refers to log-transformed, TMM-normalized counts per million ($log_2$ TMM-CPM) without jackknife (*n* - 1) resampling. **(A)** Standard deviation (SD). **(B)** $Log_2$-transformed standard deviation. **(C)** (Squared) coefficient of variation (CV, defined as the ratio of standard deviation to the mean). **(D**) $Log_2$-transformed (squared) coefficient of variation. Blue lines are LOESS regression curves and *R* is Pearson correlation coefficient.

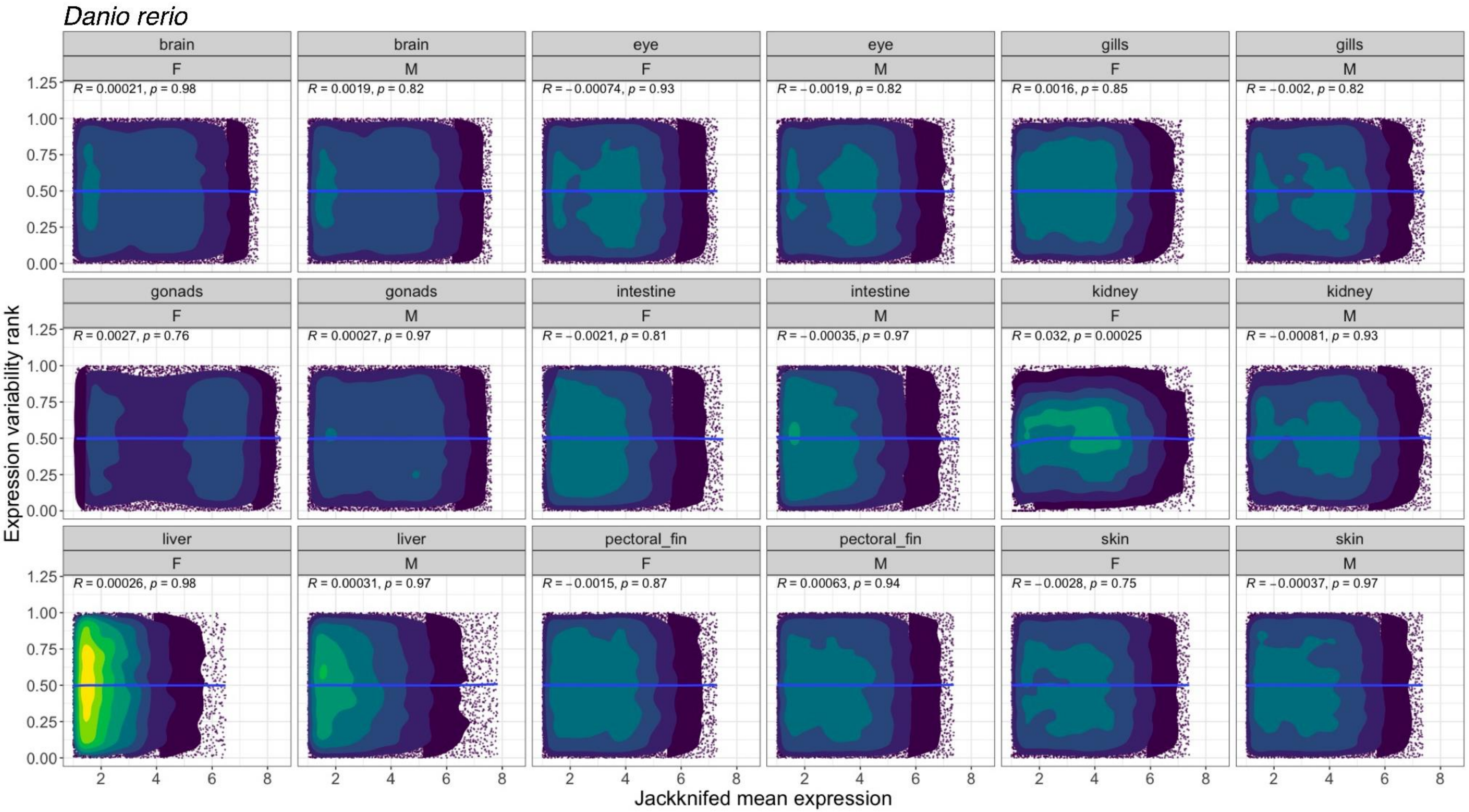


**Figure S12. Estimated expression variability is independent of mean expression for each organ-sex condition in zebrafish.** All values are means estimated from jackknife (*n* - 1) resampling. Mean expression refers to log-transformed, TMM-normalized counts per million ($\log_2$ TMM-CPM). Expression variability rank refers to the local percentile rank of residual $\log_2$-transformed (squared) coefficient of variation ($\log(CV)^2$) within sliding windows of 100 neighboring genes sorted by expression level. Blue lines are LOESS regression curves and *R* is Pearson correlation coefficient.

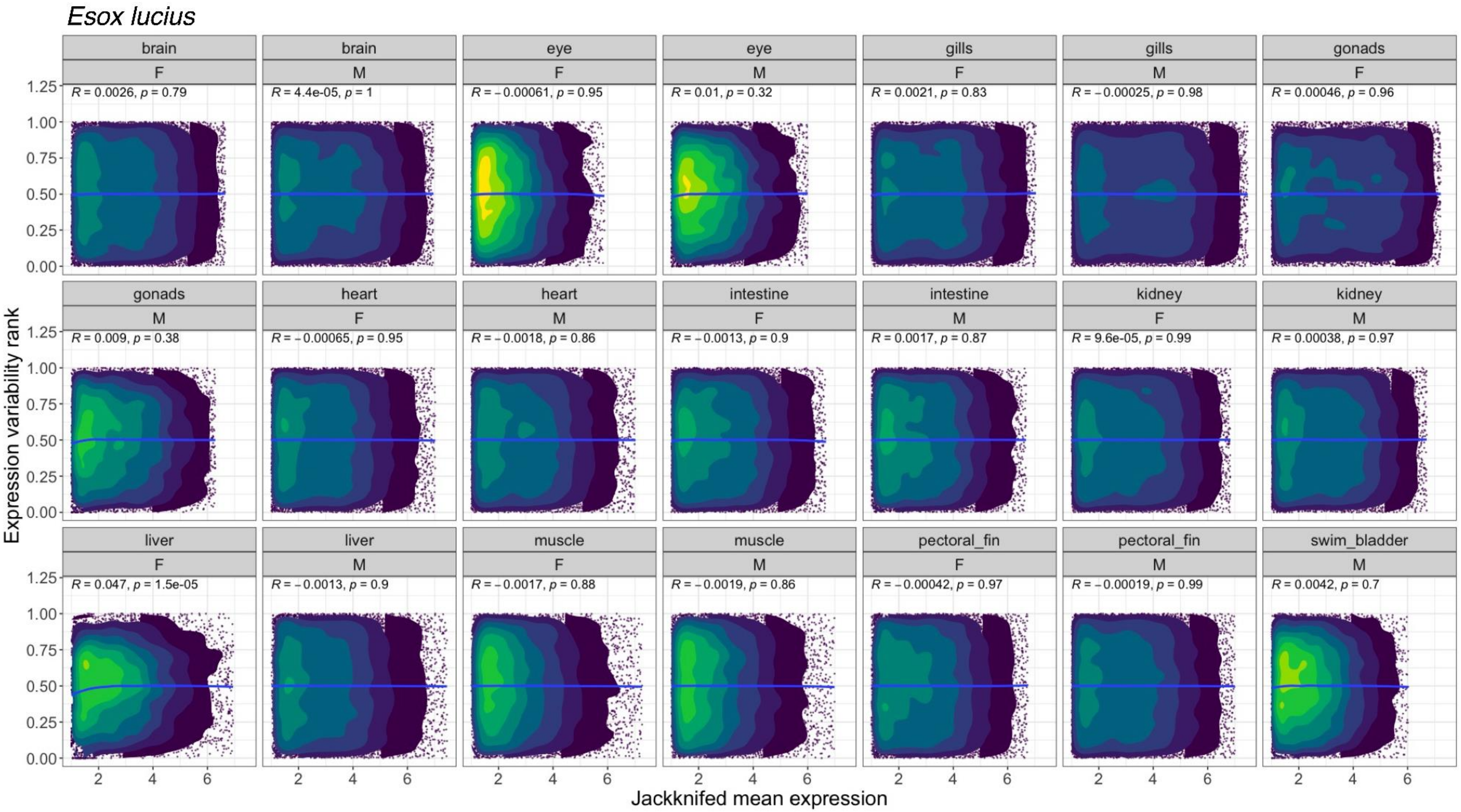


**Figure S13. Estimated expression variability is independent of mean expression for each organ-sex condition in Northern pike.** All values are means estimated from jackknife (*n* - 1) resampling. Mean expression refers to log-transformed, TMM-normalized counts per million ($log_2$ TMM-CPM). Expression variability rank refers to the local percentile rank of residual $log_2$-transformed (squared) coefficient of variation ($log(CV)^2$) within sliding windows of 100 neighboring genes sorted by expression level. Blue lines are LOESS regression curves and *R* is Pearson correlation coefficient.

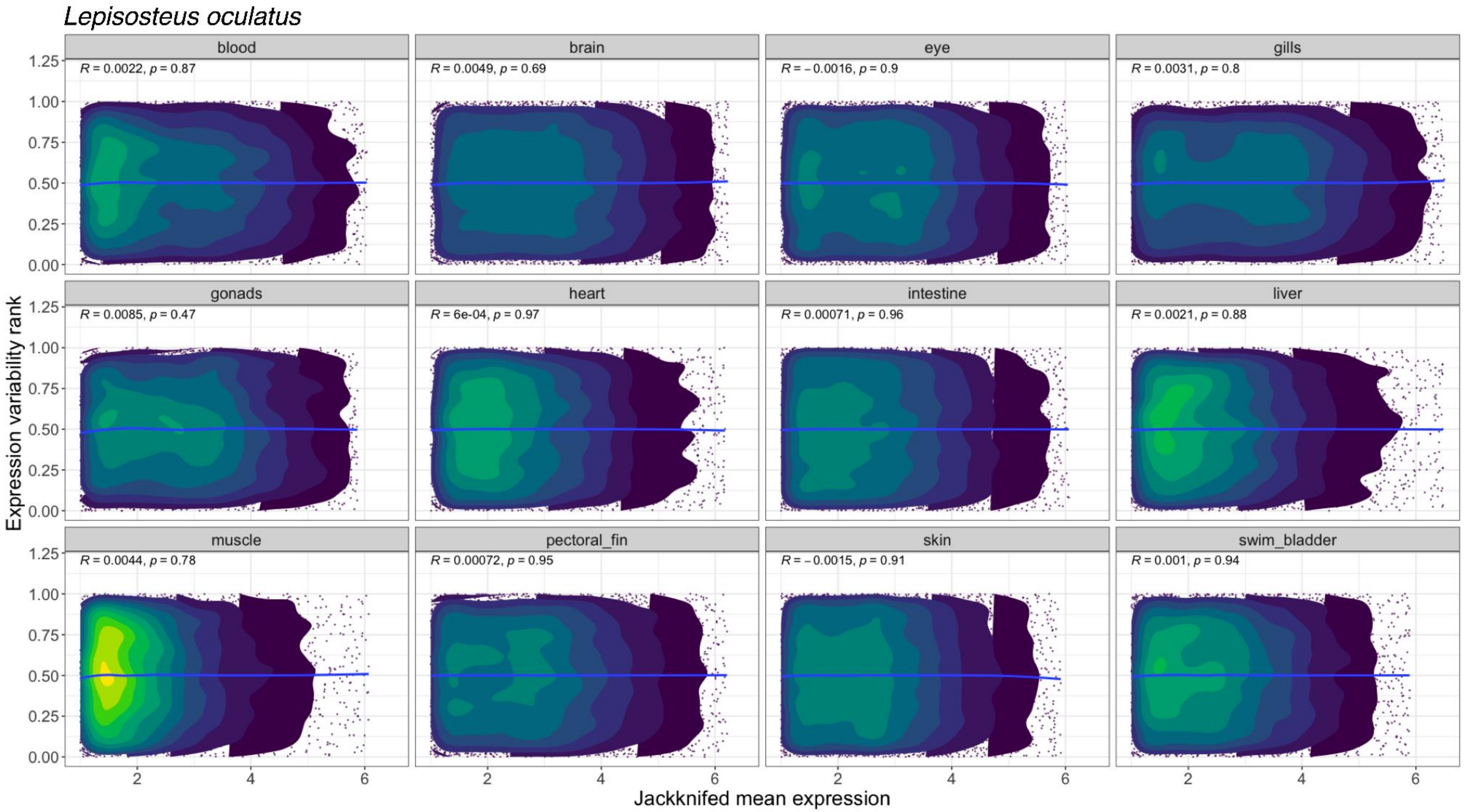


**Figure S14. Estimated expression variability is independent of mean expression for each organ in spotted gar.** All values are means estimated from jackknife (*n* - 1) resampling. Mean expression refers to log-transformed, TMM-normalized counts per million ($log_2$ TMM-CPM). Expression variability rank refers to the local percentile rank of residual $log_2$-transformed (squared) coefficient of variation ($log(CV)^2$) within sliding windows of 100 neighboring genes sorted by expression level. Blue lines are LOESS regression curves and *R* is Pearson correlation coefficient.

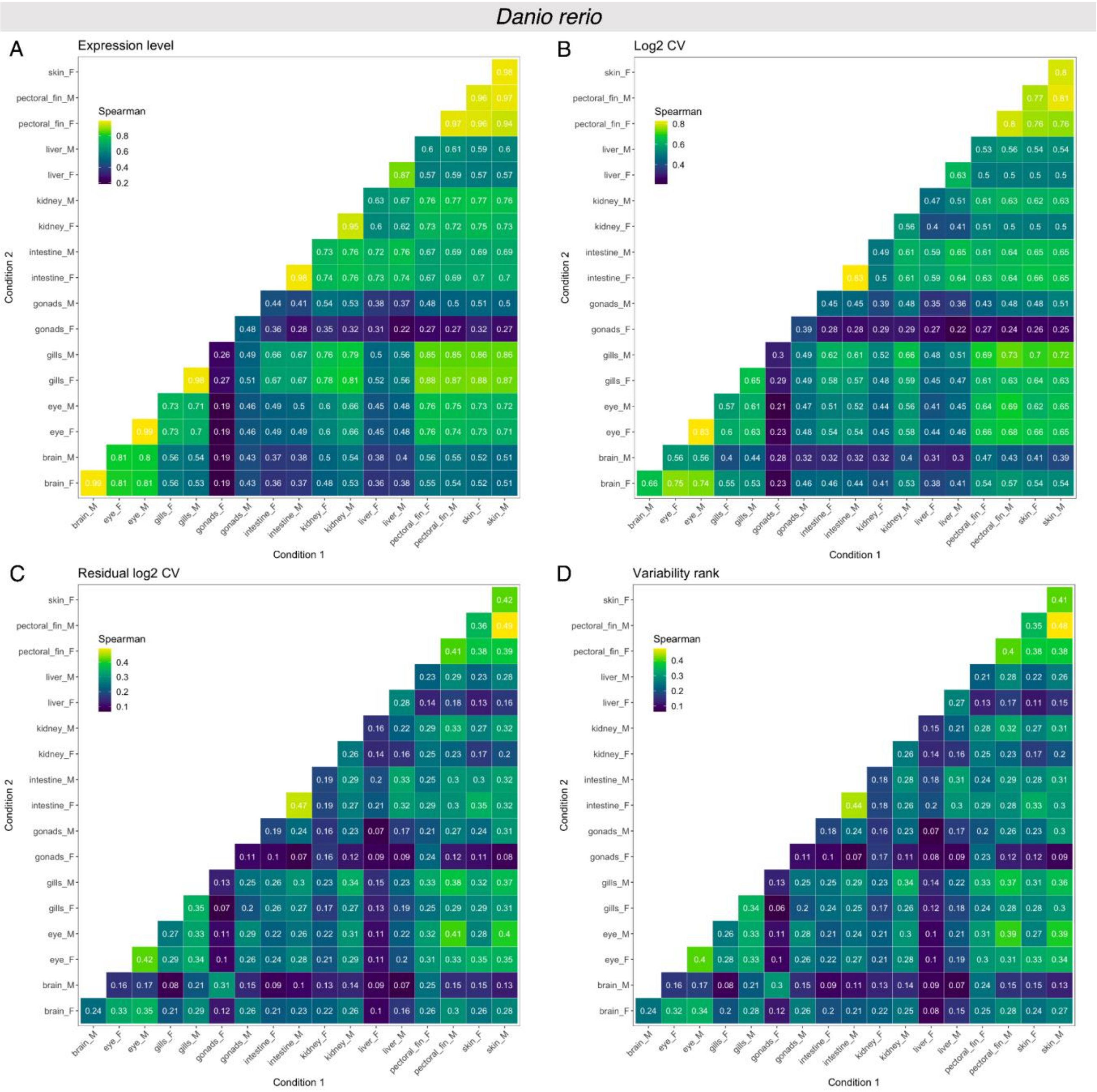


**Figure S15. Spearman rank correlation of expression level and expression variability measures between organ-sex conditions in zebrafish.** Only protein-coding genes without missing variability rank estimates across all organ-sex conditions ($n$ = 5599 genes) are included. All input values are means estimated from jackknife resampling. **(A)** Log-transformed, TMM-normalized counts per million ($\log_2$ TMM-CPM). **(B)** Log-transformed (squared) coefficient of variation (CV). **(C)** Residuals of $\log_2$ (squared) CV. **(D)** Variability rank, defined as the local percentile rank of residual $\log_2(CV)^2$ within sliding windows of 100 neighboring genes. Note that color scale ranges are not identical across (A)-(D).

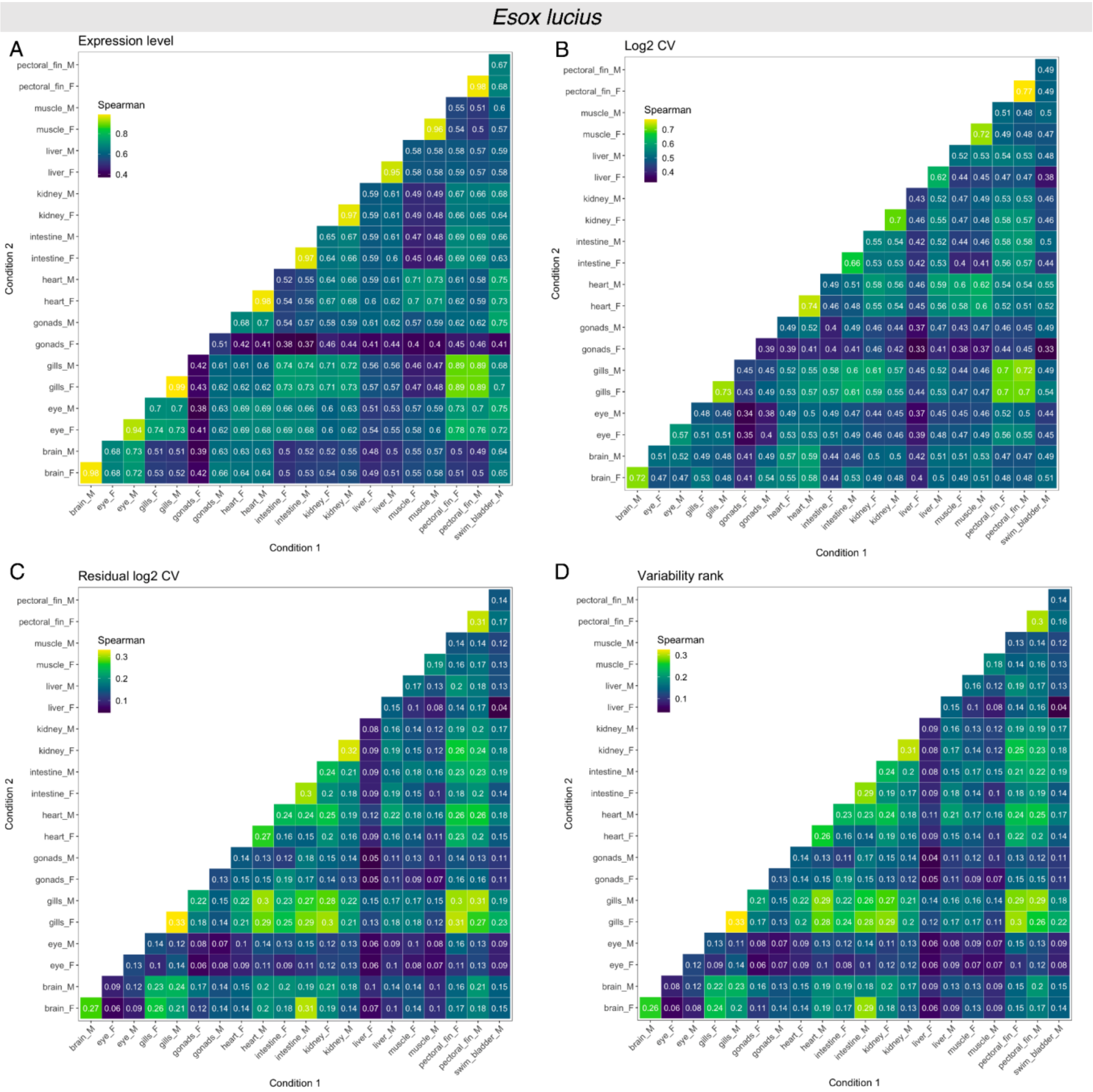


**Figure S16. Spearman rank correlation of expression level and expression variability measures between organ-sex conditions in Northern pike.** Only protein-coding genes without missing variability rank estimates across all organ-sex conditions ($n$ = 4567 genes) are included. All input values are means estimated from jackknife resampling. **(A)** Log-transformed, TMM-normalized counts per million ($\log_2$ TMM-CPM). **(B)** Log-transformed (squared) coefficient of variation (CV). **(C)** Residuals of $\log_2$ (squared) CV. **(D)** Variability rank, defined as the local percentile rank of residual $\log_2(CV)^2$ within sliding windows of 100 neighboring genes. Note that color scale ranges are not identical across (A)-(D).

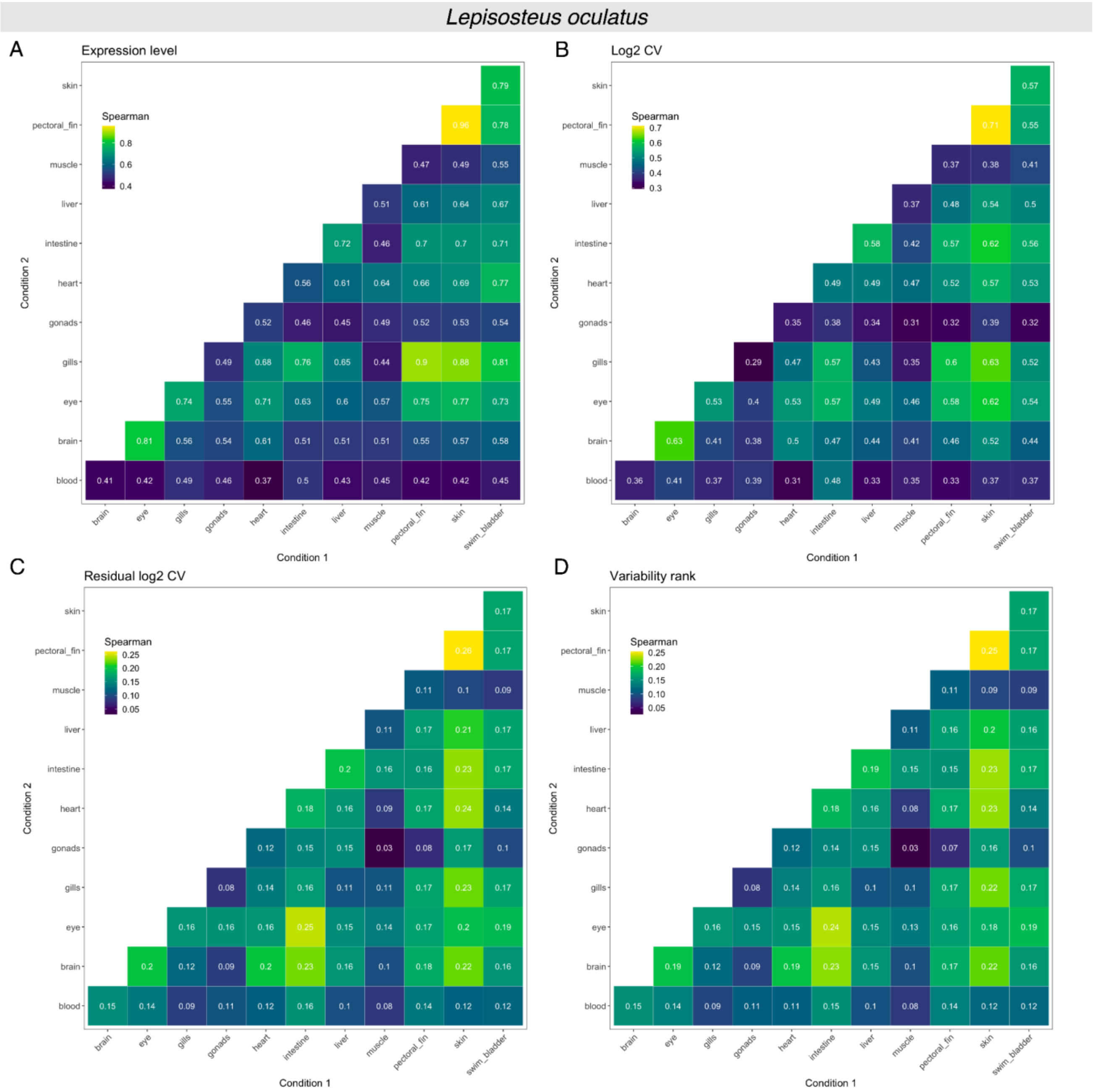


**Figure S17. Spearman rank correlation of expression level and expression variability measures between organ-sex conditions in spotted gar.** Only protein-coding genes without missing variability rank estimates across all organ-sex conditions (*n* = 2607 genes) are included. All input values are means estimated from jackknife resampling. **(A)** Log-transformed, TMM-normalized counts per million ($log_2$ TMM-CPM). **(B)** Log-transformed (squared) coefficient of variation (CV). **(C)** Residuals of $log_2$ (squared) CV. **(D)** Variability rank, defined as the local percentile rank of residual $log_2(CV)^2$ within sliding windows of 100 neighboring genes. Note that color scale ranges are not identical across (A)-(D).

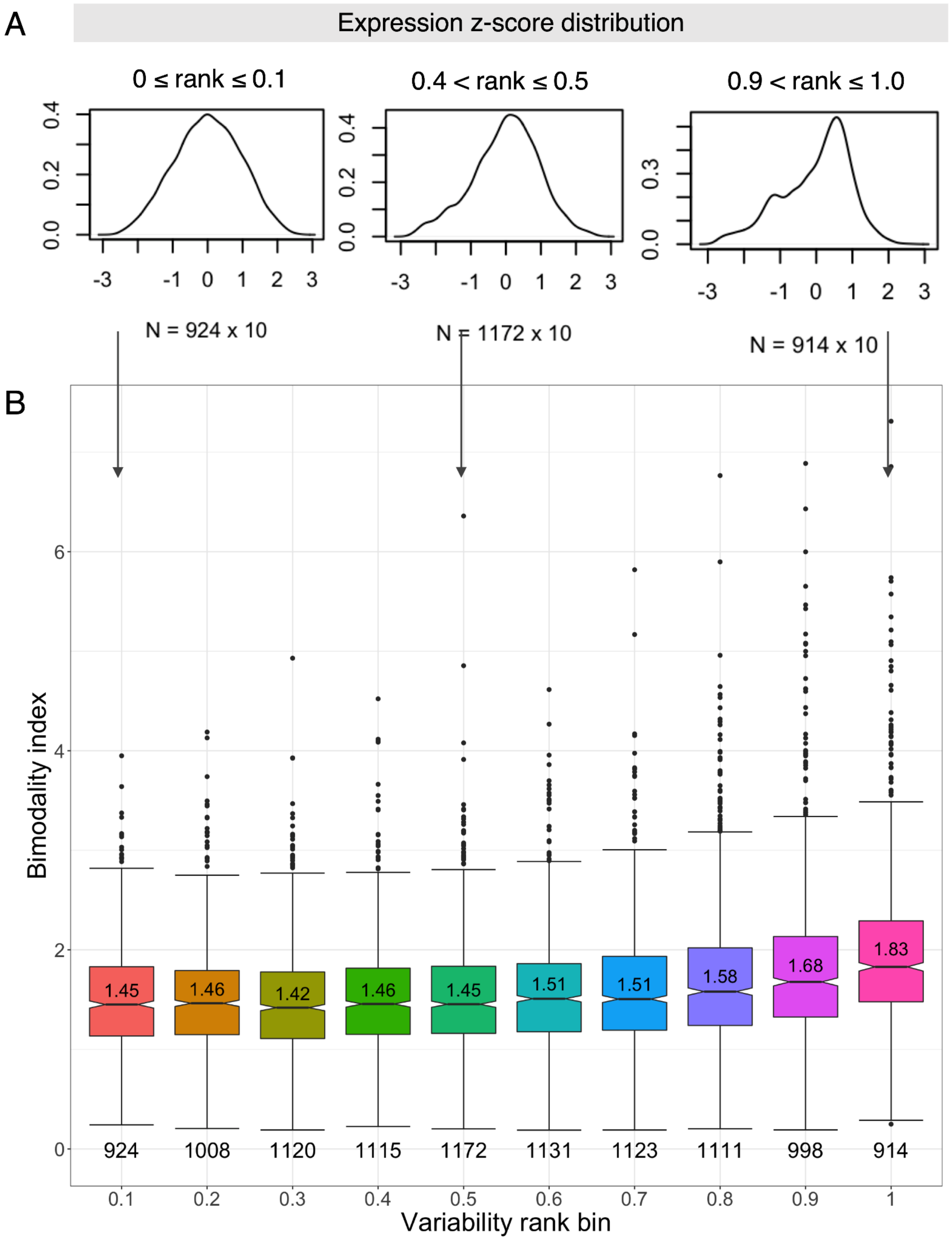


**Figure S18. Bimodality tests for protein-coding genes grouped by expression variability rank.** Data from female Northern pike brain ($n$ = 10 samples). Genes were sorted into ten variability rank bins of size 0.1. **(A)** Kernel density distributions of expression level z-scores for three variability rank bins. The number of data points $N$ includes the number of genes within the bin times the number of sampled individuals for the given condition. See **Figure S19A** for the z-score distributions for all 11 variability rank bins ranging from from 0.0 to 1.0. **(B)** Boxplots showing distributions of the bimodality index (BI) (Wang et al. 2009) for genes binned by variability rank. For each bin, the median BI and number of genes is indicated.

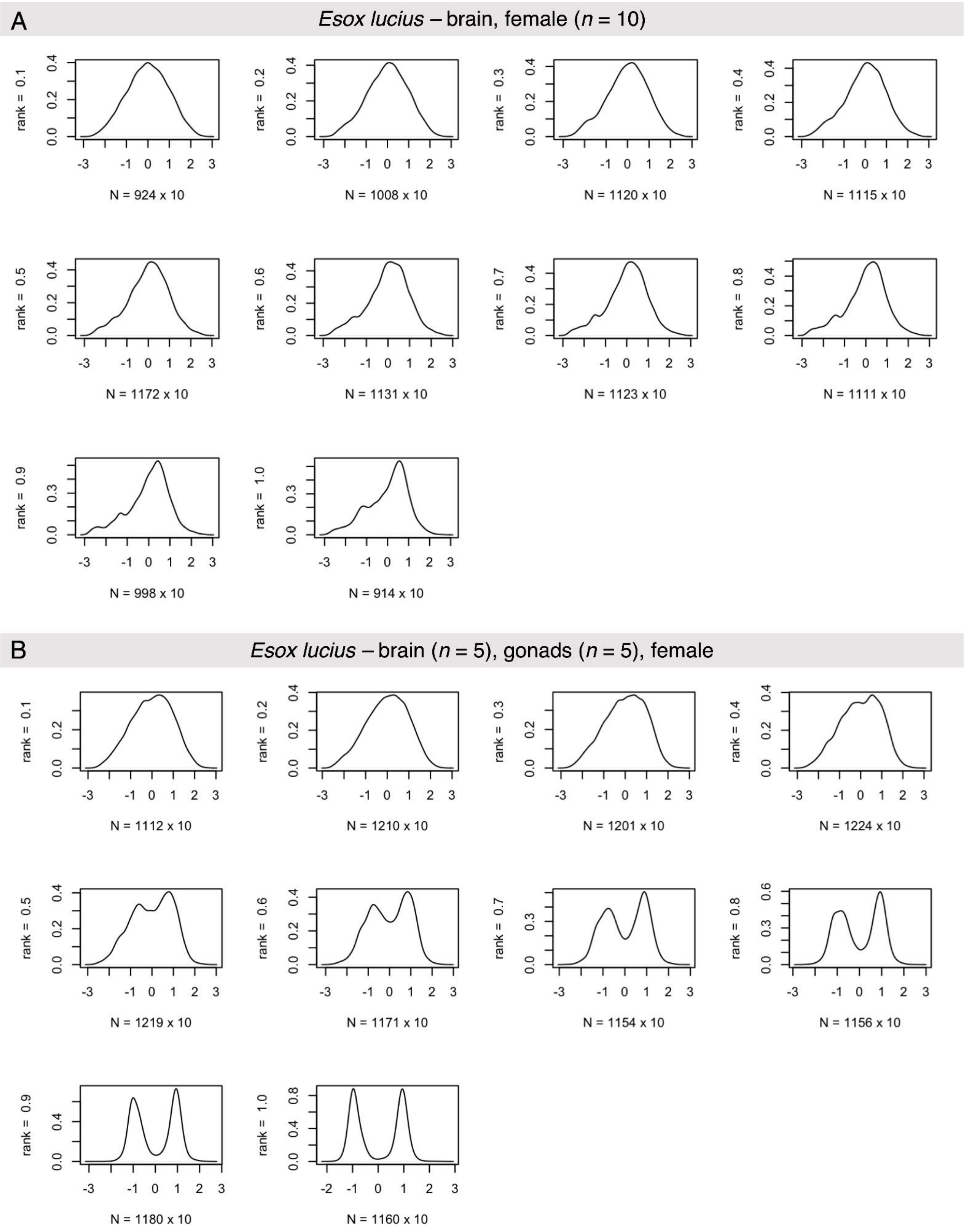


**Figure S19. Kernel density distribution of expression level z-scores for genes sorted by variability rank.** Protein-coding genes were sorted into ten variability rank bins of size 0.1. The number of data points *N* includes the number of genes within the bin times the number of sampled individuals for the given condition. **(A)** Data from female Northern pike brain (*n* = 10 samples). **(B)** Pseudo-condition based on data from female Northern pike brain and ovary (*n* = 5 for each organ), with the same total sample size as (A). Replicates from two organs were combined to generate strongly bimodal expression profiles.

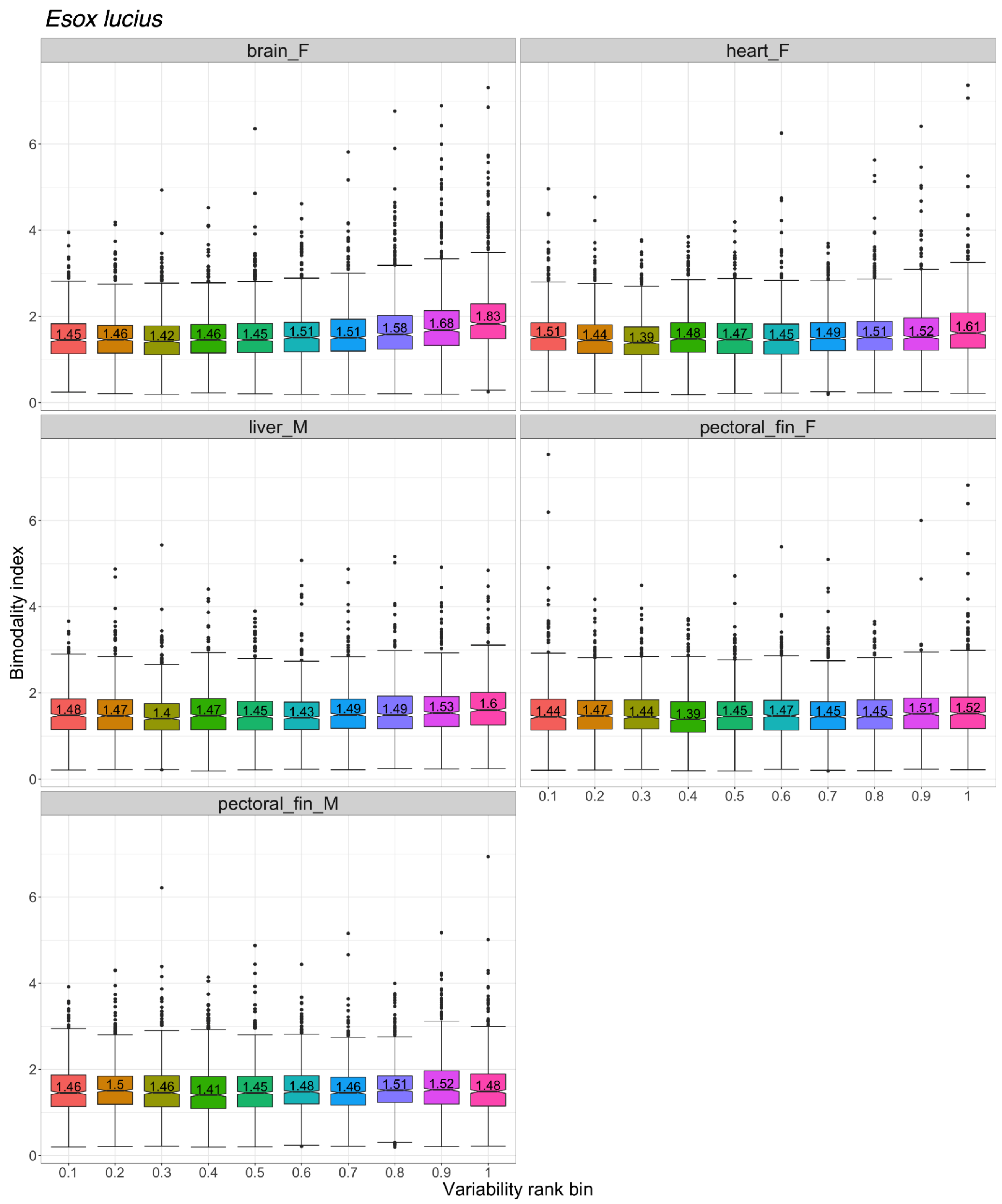


**Figure S20. Bimodality index (BI) distribution of genes sorted by variability rank for organ-sex conditions in Northern pike.** Only conditions with $n \geq 10$ replicates were considered. Protein-coding genes were sorted into ten variability rank bins of size 0.1. Median BI is indicated for each bin.

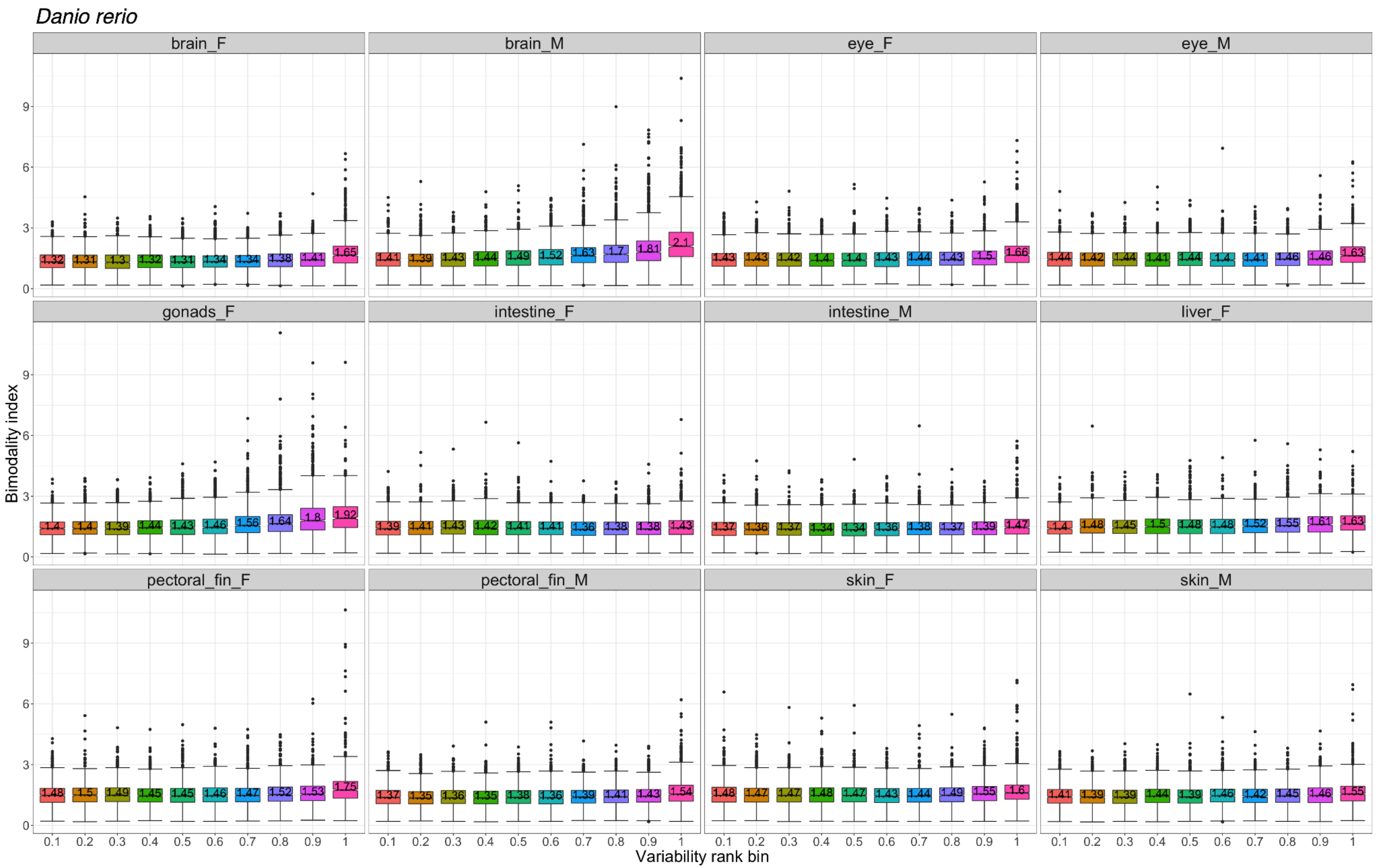


**Figure S21. Bimodality index (BI) distribution of genes sorted by variability rank for organ-sex conditions in zebrafish.** Only conditions with $n \geq 10$ replicates were considered. Protein-coding genes were sorted into ten variability rank bins of size 0.1. Median BI is indicated for each bin.

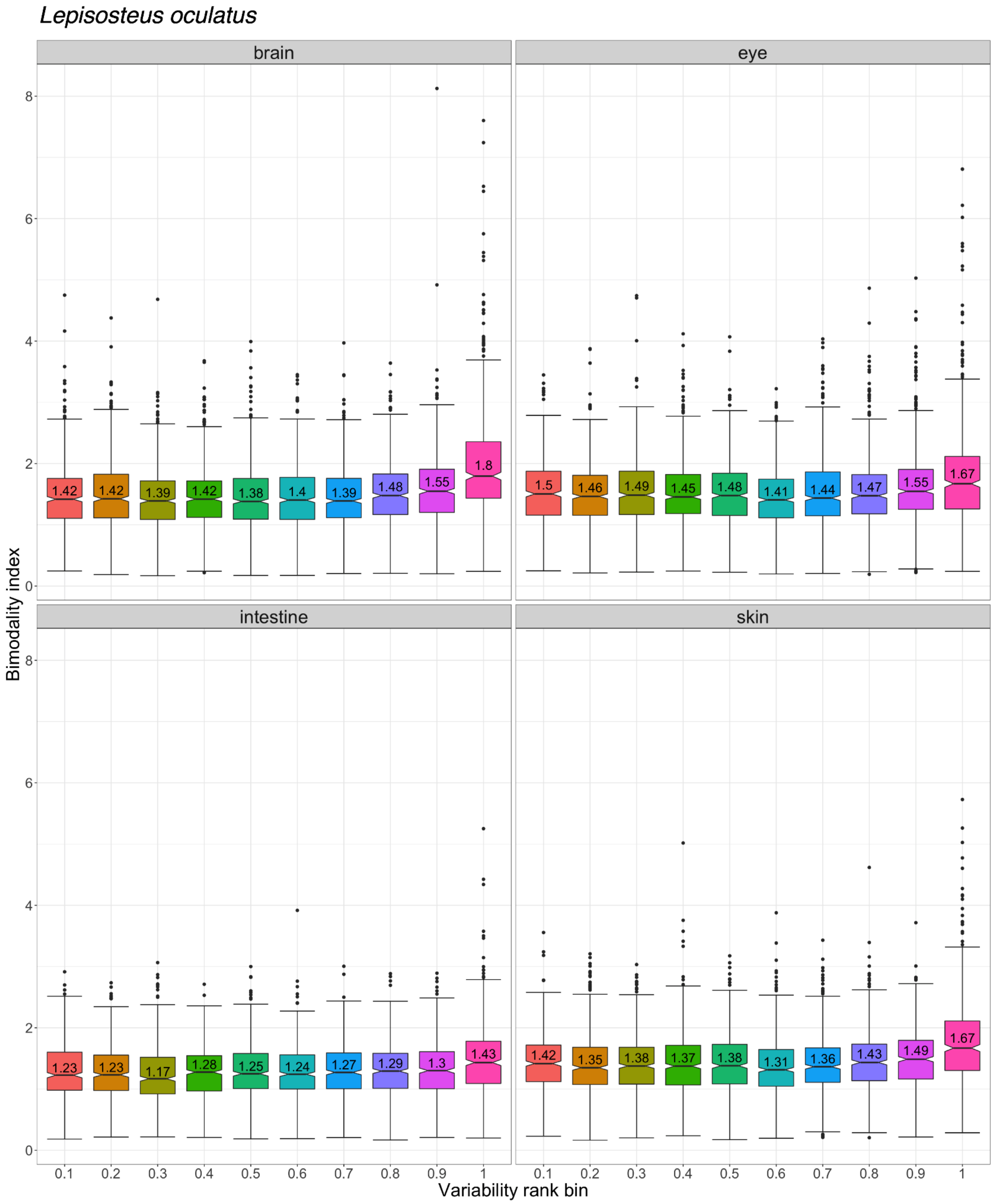


**Figure S22. Bimodality index (BI) distribution of genes sorted by variability rank for organs in spotted gar.** Only conditions with $n \geq 10$ replicates were considered. Protein-coding genes were sorted into ten variability rank bins of size 0.1. Median BI is indicated for each bin.

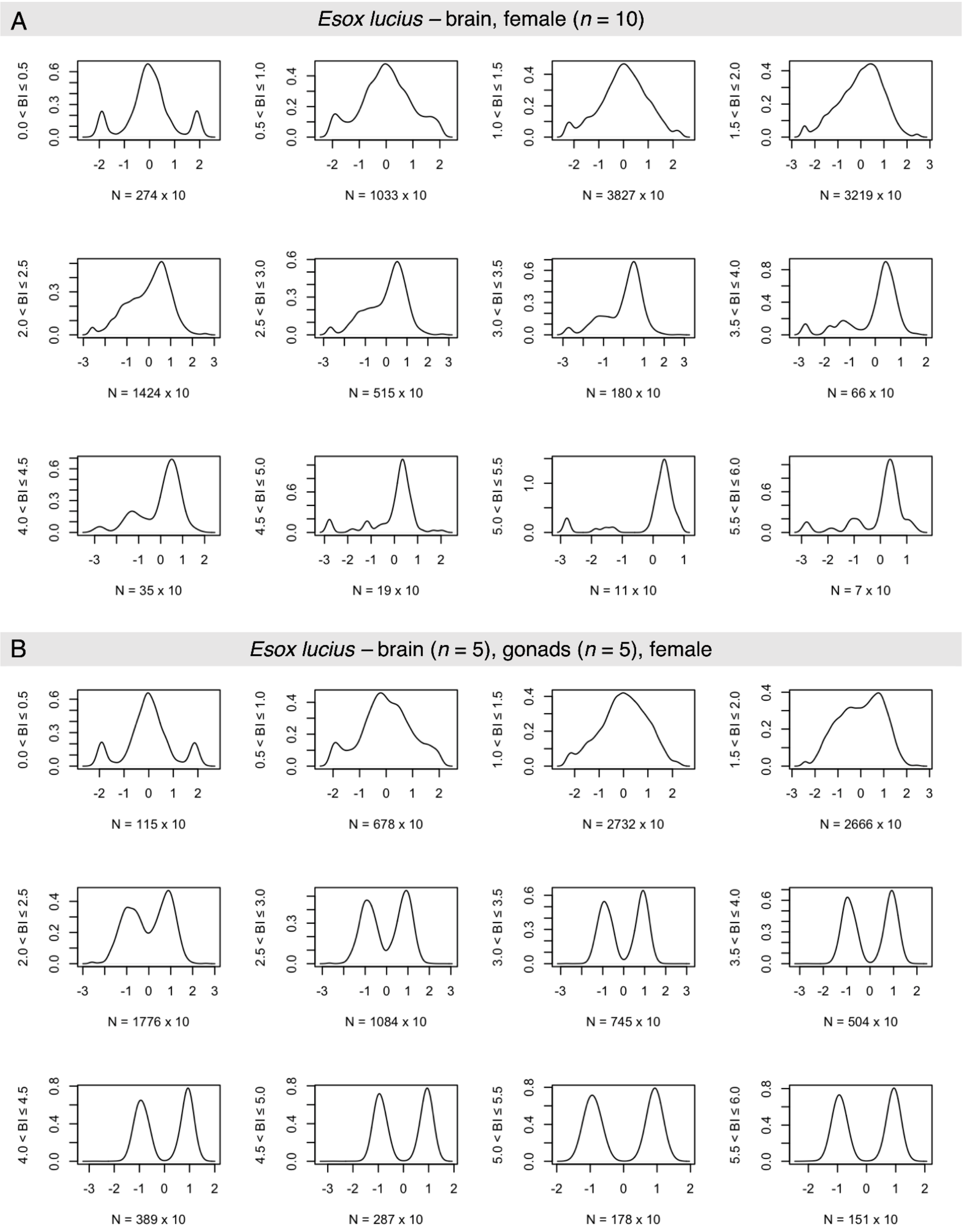


**Figure S23. Kernel density distribution of expression level z-scores for genes sorted by bimodality index (BI).** Protein-coding genes were grouped into 12 BI bins of size 0.5. The number of data points *N* includes the number of genes within the bin times the number of individuals for the given condition. **(A)** Data from female Northern pike brain (*n* = 10 samples). **(B)** Pseudo-condition based on data from female Northern pike brain and ovary (*n* = 5 for each organ), with the same total sample size as (A). Replicates from two organs were combined to generate strongly bimodal expression profiles.

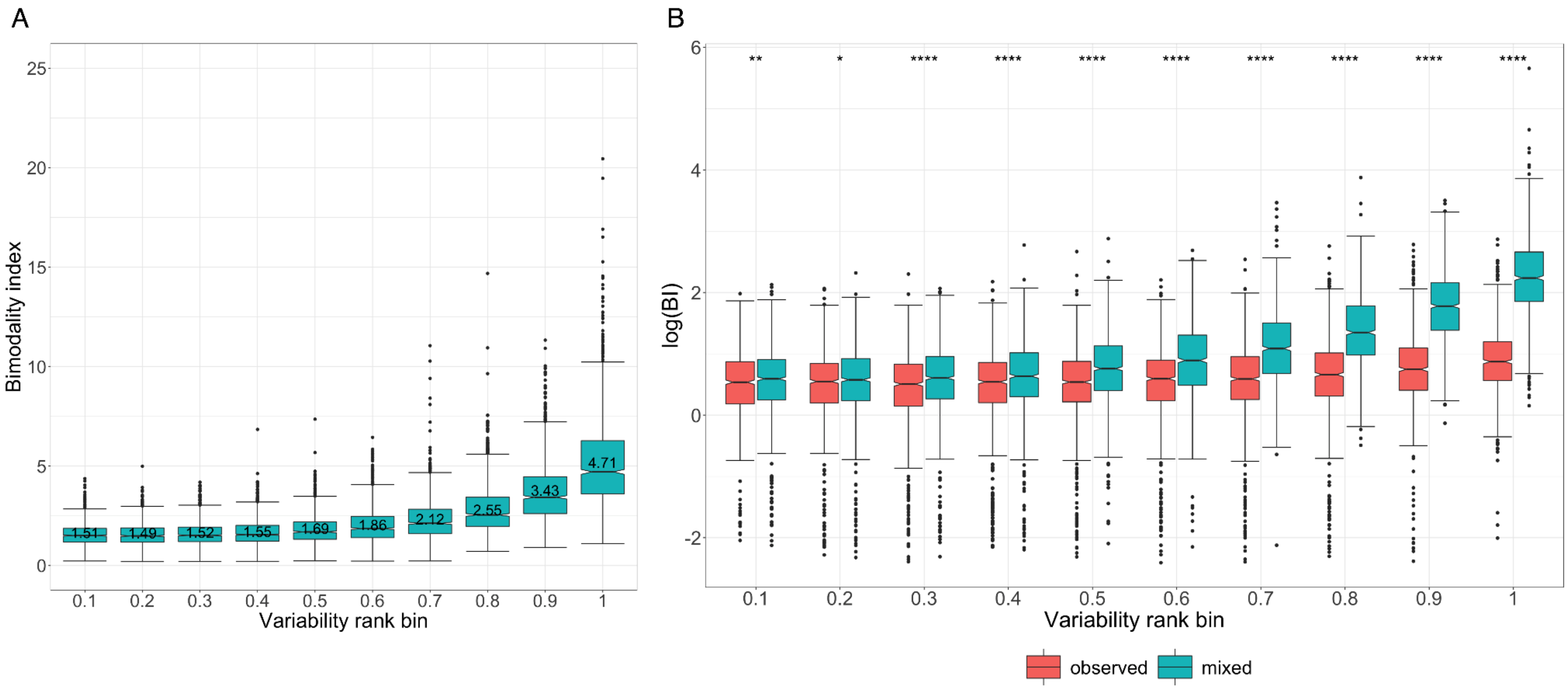


**Figure S24. Bimodality index (BI) distribution for genes sorted by variability rank given observed and mixed conditions.** Protein-coding genes were sorted into ten variability rank bins of size 0.1. **(A)** Mixed pseudo-condition combining data from female Northern pike brain ($n$ = 5) and ovary ($n$ = 5). **(B)** $Log_2$-transformed bimodality index (BI) distributions given data from female Northern pike brain ($n$ = 10) ('observed', red boxplots) and the pseudo-condition in (A) ('mixed', blue boxplots). Between-group comparisons were performed using Wilcoxon rank-sum tests ($p$-value significance levels: '****': $p \leq 0.0001$, '***': $p \leq 0.001$, '**': $p \leq 0.01$, '*': $p \leq 0.05$, 'ns': $p > 0.05$).

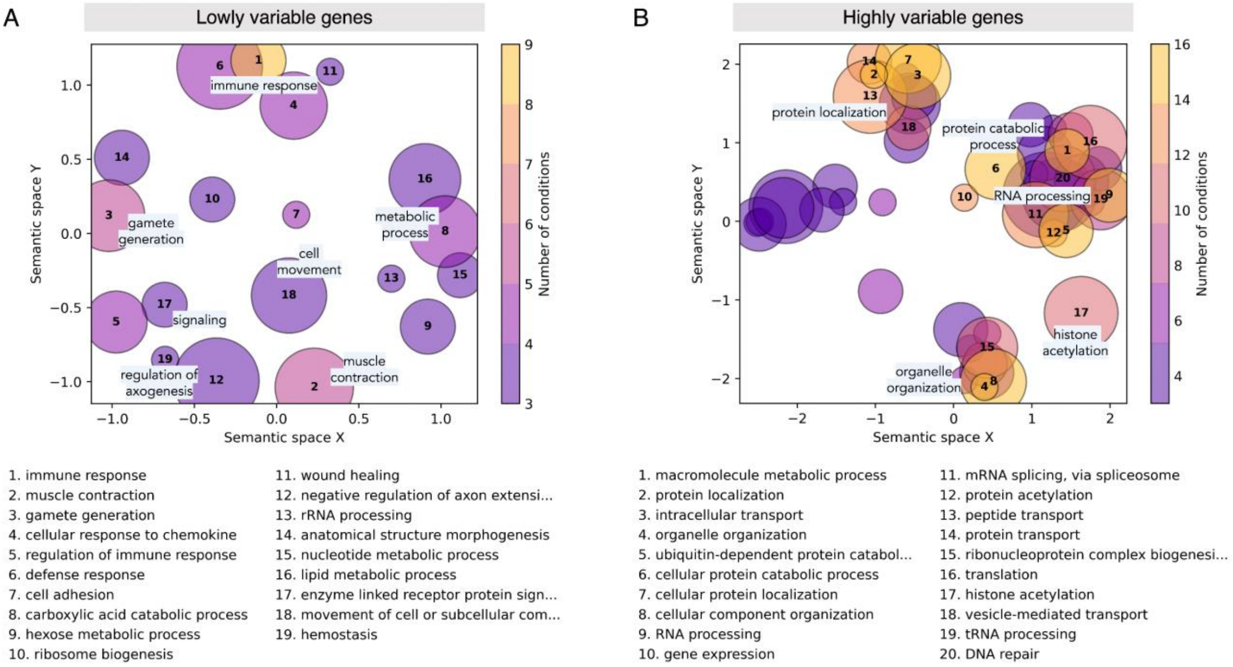


**Figure S25. Semantic similarity bubble plots of depleted Gene Ontology (GO) biological process terms for (A) underdispersed (variability rank ≤ 0.20) and (B) overdispersed (variability rank ≥ 0.80) genes in zebrafish organs.** Under-representation analysis of GO terms was performed separately for the set of under- and overdispersed genes in each organ-sex condition using the conditional hypergeometric test in *GOstats* (Falcon and Gentleman 2007). Terms that are significantly depleted (unadjusted *p*-value < 0.01) in at least three organ-sex conditions were retained. Redundancy reduction and visualization of terms was performed using *GO-Figure!* (Reijnders and Waterhouse 2021). Each circle represents a group of similar GO terms collapsed into a representative term. Representative terms are plotted in a semantic similarity space such that more similar terms are positioned closer together. The top 20 representative terms are listed and sorted by the number of organ-sex conditions in which the representative is significant. Circle color indicates the number of conditions in which a representative term is significant while circle size is scaled by the number of related terms summarized in a cluster.

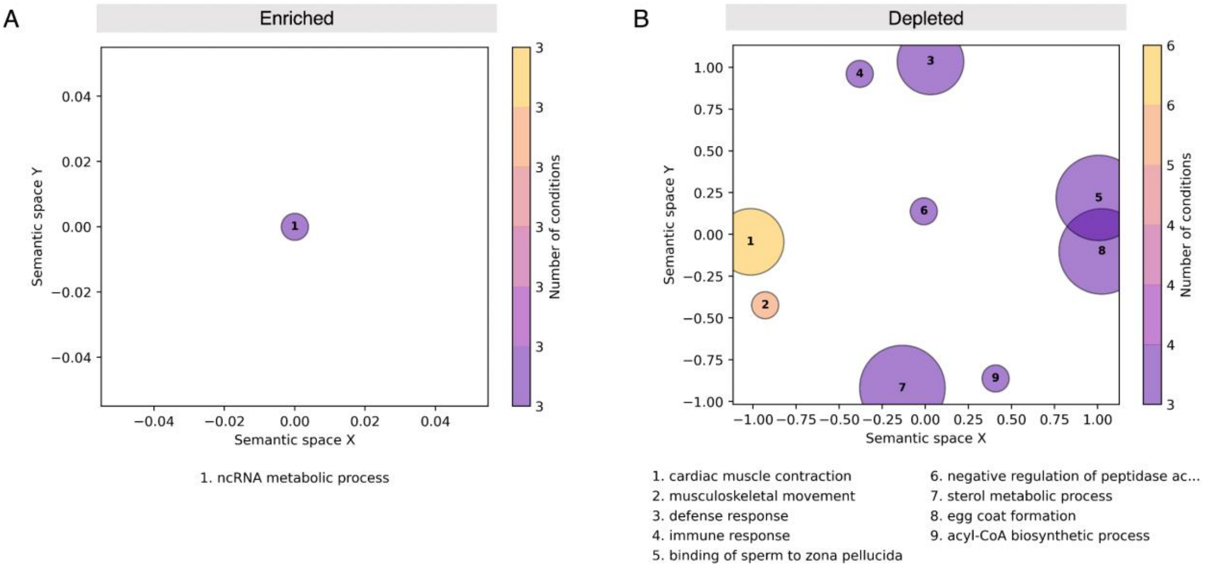


**Figure S26. Semantic similarity bubble plots of (A) enriched and (B) depleted GO biological process terms for moderately variable (0.40 ≤ variability rank ≤ 0.60) genes in zebrafish organs.** Over- and under-representation analysis of GO terms were performed separately for the set of moderately variable genes in each organ-sex condition using the conditional hypergeometric test in *GOstats* (Falcon and Gentleman 2007). Terms that are significant (unadjusted *p*-value < 0.01) in at least three organ-sex conditions were retained. Redundancy reduction and visualization of terms was performed using *GO-Figure!* (Reijnders and Waterhouse 2021), as previously described (**Figure 3, Figure S25**).

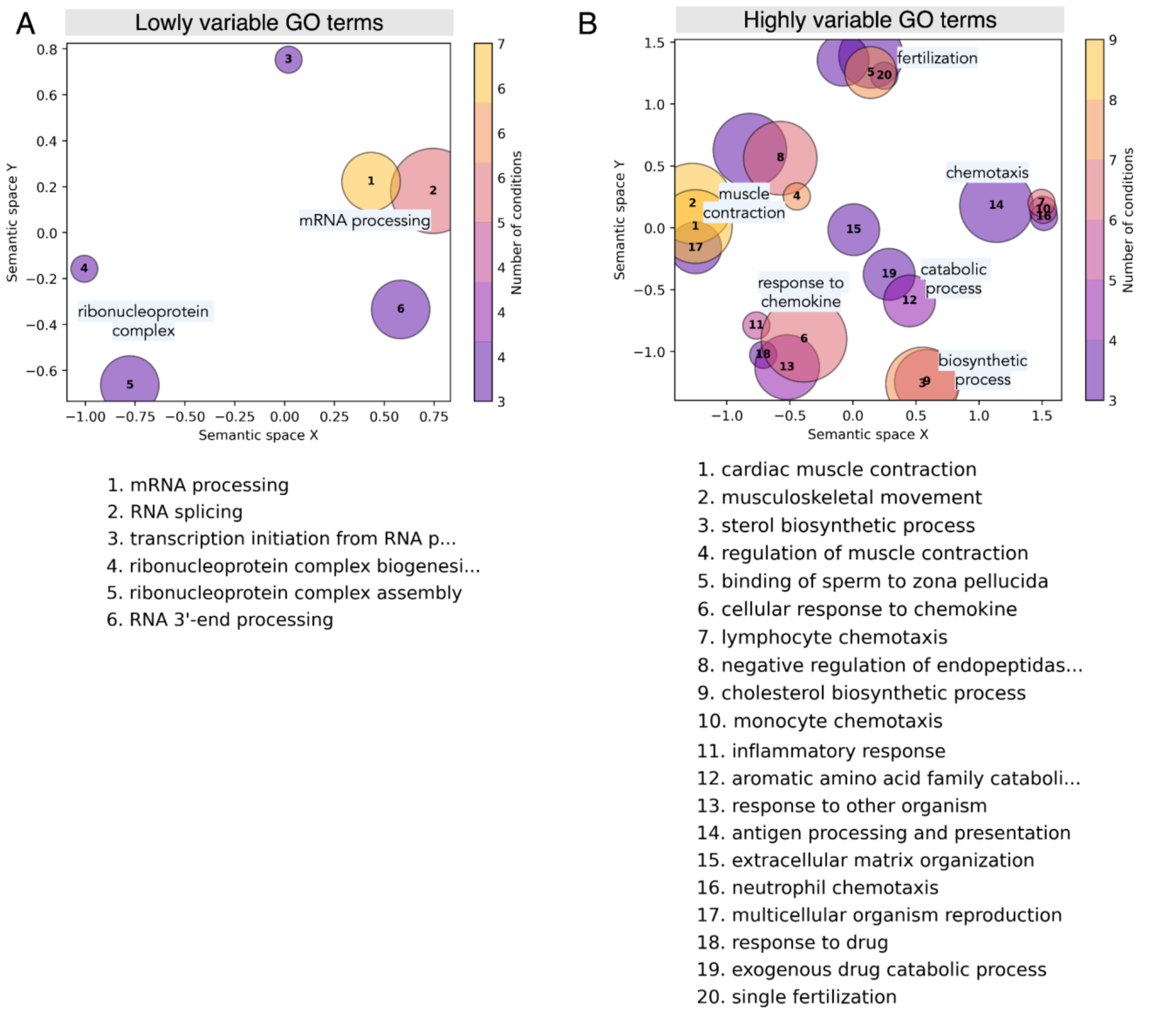


**Figure S27. Semantic similarity bubble plots of (A) lowly variable and (B) highly variable Gene Ontology (GO) biological process terms as measured in zebrafish organs.** GO terms consisting of at least 20 genes and no more than 1000 genes were used as input to a competitive gene set analysis implemented using *cameraPR* in *limma* (Wu and Smyth 2012), with variability rank as the genewise statistic. Terms that are significantly lowly or highly variable (FDR < 0.05) in at least three organ-sex conditions were retained. Redundancy reduction and visualization of terms was performed using *GO-Figure!* (Reijnders and Waterhouse 2021). Each circle represents a group of similar GO terms collapsed into a representative term. Representative terms are plotted in a semantic similarity space such that more similar terms are positioned closer together. The top 20 representative terms are listed and sorted by the number of organ-sex conditions in which the representative is significant. Circle color indicates the number of conditions in which a representative term is significant while circle size is scaled by the number of related terms summarized in a cluster.

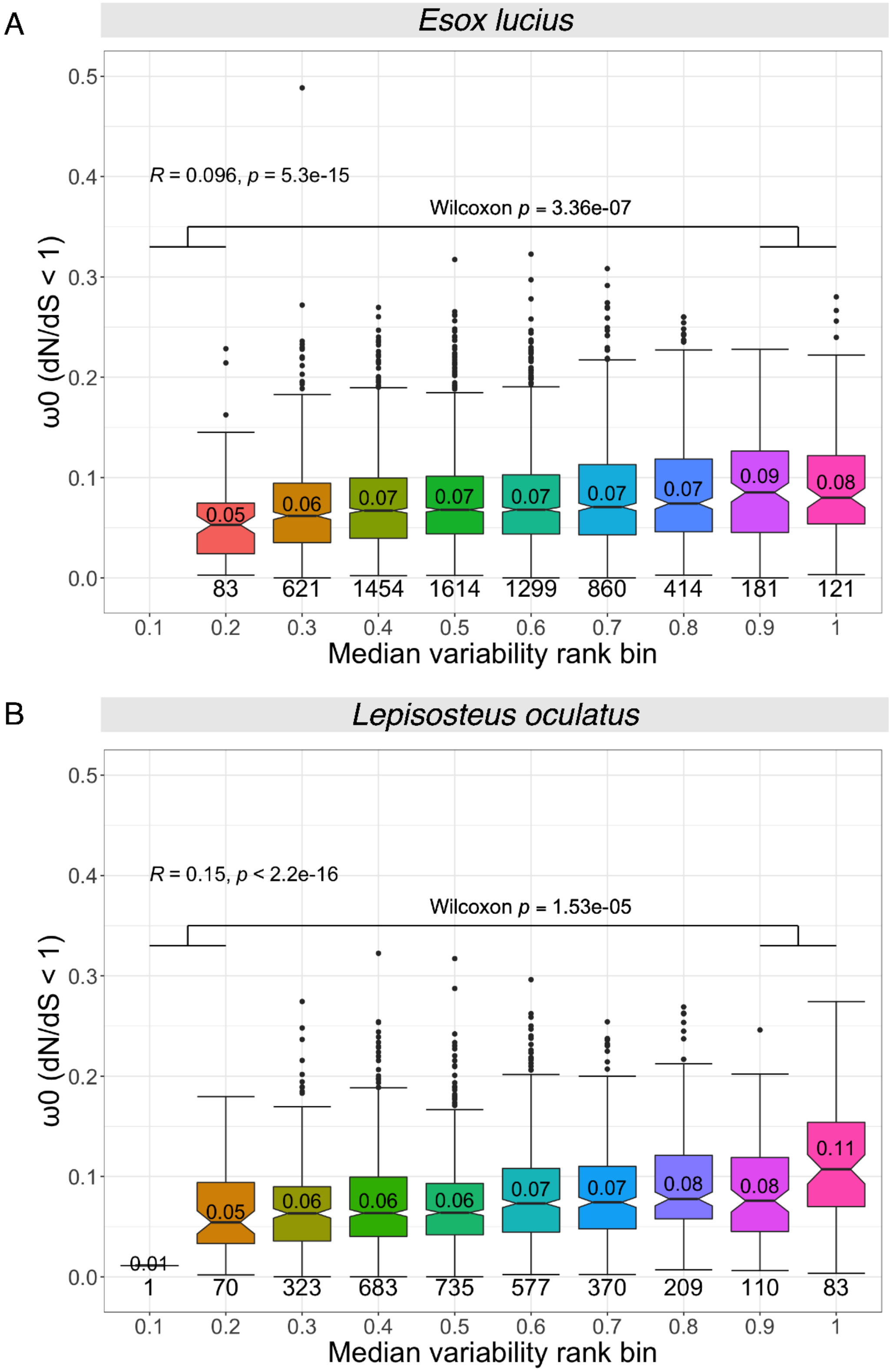


**Figure S28. Purifying selection of (A) Northern pike and (B) spotted gar genes grouped by median expression variability across conditions.** Boxplots show the distribution of ratios of nonsynonymous substitutions ($d_N$) to synonymous substitutions ($d_S$) of sites that have evolved under purifying selection ($\omega_0$ ratio, for $d_N/d_S < 1$). Selection statistics based on conservative branch-site likelihood tests were retrieved from the Selectome database (Moretti et al. 2014; Proux et al. 2009), with tested branches as shown in **Figure 4A**. Protein-coding genes were grouped into 10 bins of size 0.1 ([0,0.1), (0.1,0.2], …, (0.8,0.9], (0.9,1.0]) based on their median variability rank across conditions. The median $\omega_0$ is indicated within each boxplot and the number of genes included are annotated below each. *R* is the Spearman's correlation coefficient between $\omega_0$ and the unbinned median variability across conditions, with the corresponding *p*-value. The Wilcoxon *p*-value corresponds to a Wilcoxon rank-sum test comparing the least variable genes (variability rank ≤ 0.2) against the most variable genes (variability rank > 0.8).

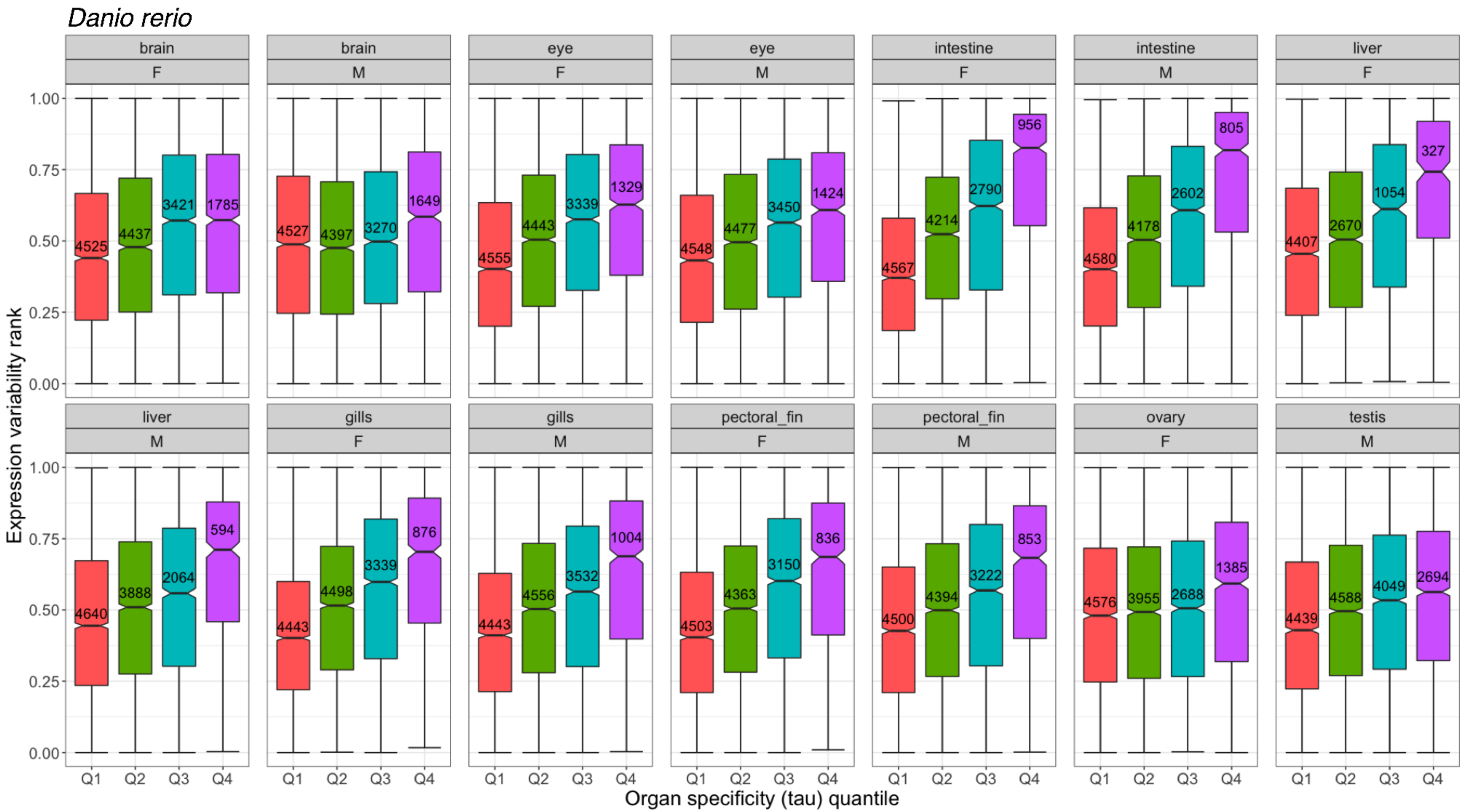


**Figure S29. Relationship between expression variability rank and organ specificity index across organ-sex conditions in zebrafish.** Boxplots show the distribution of variability ranks grouped by organ expression specificity (tau, $\tau$) quartiles. Quartiles were computed considering only the set of genes with both organ specificity and expression variability measurements ($n$ = 19358 genes). The $\tau$ cutoff for the first to third quartiles are as follows: $Q_1$ (25th percentile): $\tau = 0.320$; $Q_2$ (50th percentile): $\tau = 0.539$, $Q_3$ (75th percentile): $\tau = 0.751$. Each quartile is annotated by the number of genes included.

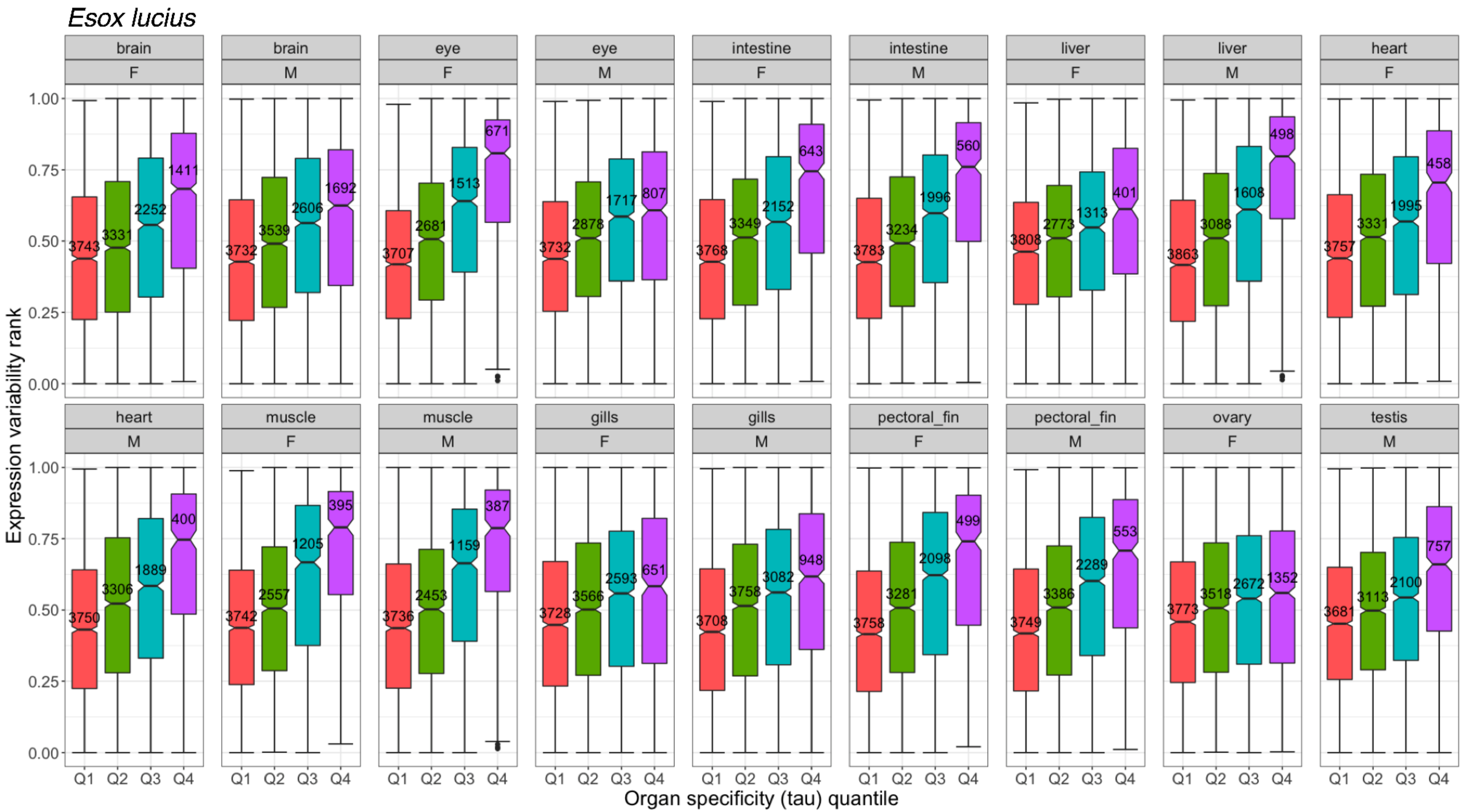


**Figure S30. Relationship between expression variability rank and organ specificity index across organ-sex conditions in Northern pike.** Boxplots show the distribution of variability ranks grouped by organ expression specificity (tau, $\tau$) quartiles. Quartiles were computed considering only the set of genes with both organ specificity and expression variability measurements (*n* = 16025 genes). The $\tau$ cutoff for the first to third quartiles are as follows: $Q_1$ (25th percentile): $\tau = 0.301$; $Q_2$ (50th percentile): $\tau = 0.517$, $Q_3$ (75th percentile): $\tau = 0.750$. Each quartile is annotated by the number of genes included.

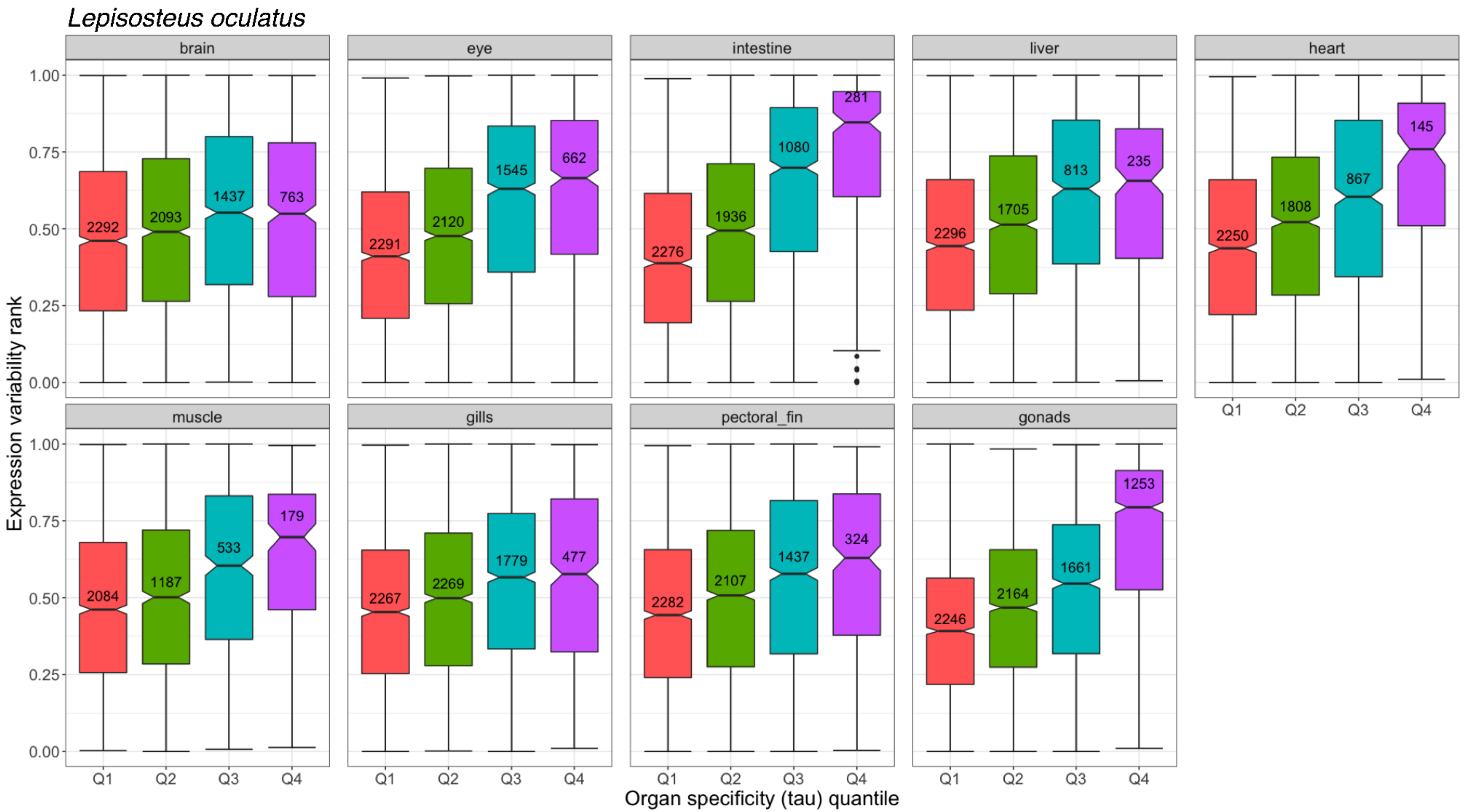


**Figure S31. Relationship between expression variability rank and organ specificity index across organs in spotted gar.** Boxplots show the distribution of variability ranks grouped by organ expression specificity (tau, $\tau$) quartiles. Quartiles were computed considering only the set of genes with both organ specificity and expression variability measurements (*n* = 9681 genes). The $\tau$ cutoff for the first to third quartiles are as follows: $Q_1$ (25th percentile): $\tau = 0.283$; $Q_2$ (50th percentile): $\tau = 0.503$, $Q_3$ (75th percentile): $\tau = 0.766$. Each quartile is annotated by the number of genes included.

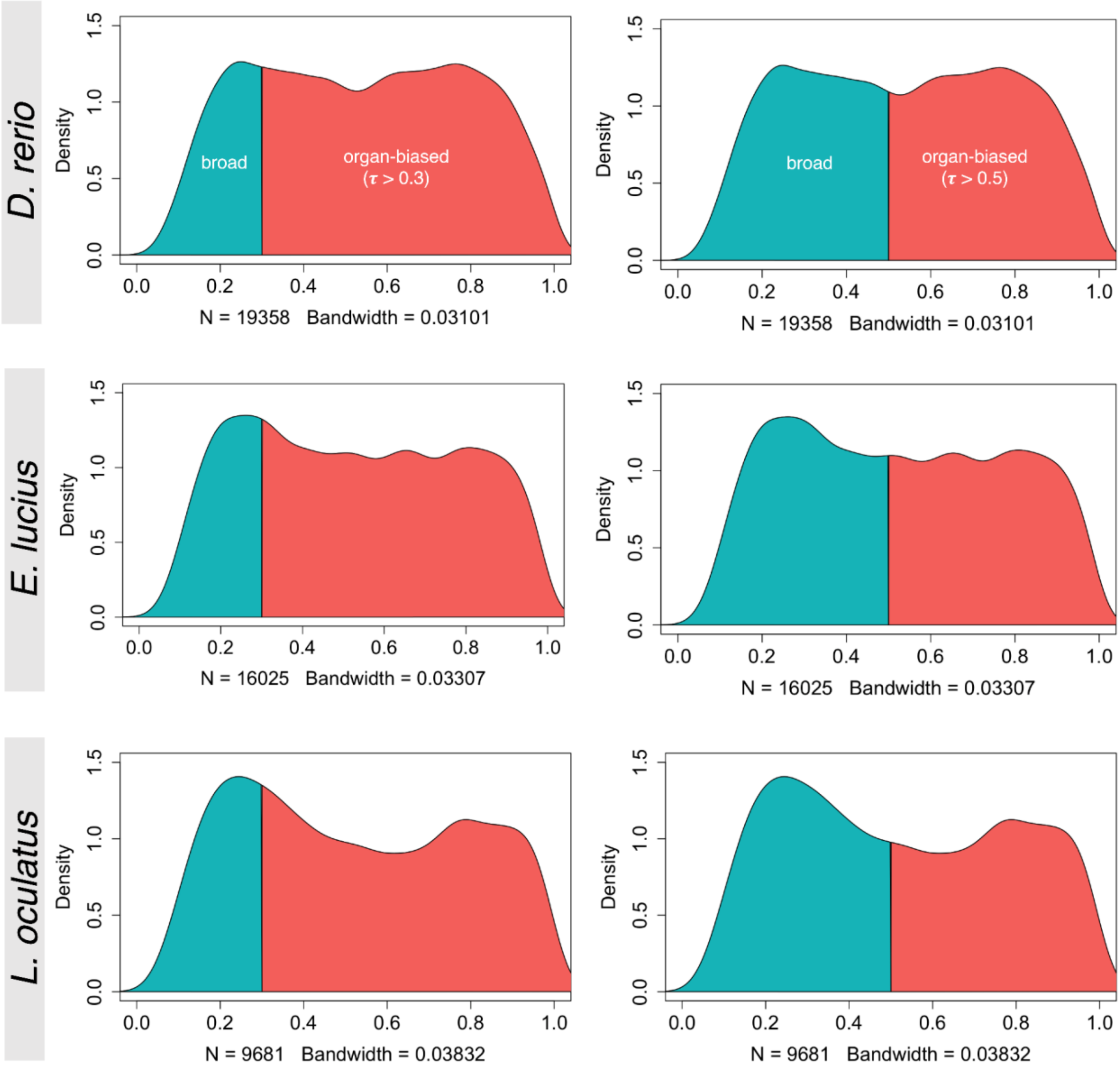


**Figure S32. Organ expression specificity distributions.** Kernel density distribution plots of tau ($\tau$) for zebrafish (top row), Northern pike (middle row), and spotted gar (bottom row). Only genes with both organ specificity and expression variability measurements are included (*N*). Left column: A cutoff of $\tau > 0.3$ distinguishes broadly expressed genes (turquoise) from organ-biased genes (salmon). Right column: A higher cutoff of $\tau > 0.50$ is applied to distinguish the two gene categories.

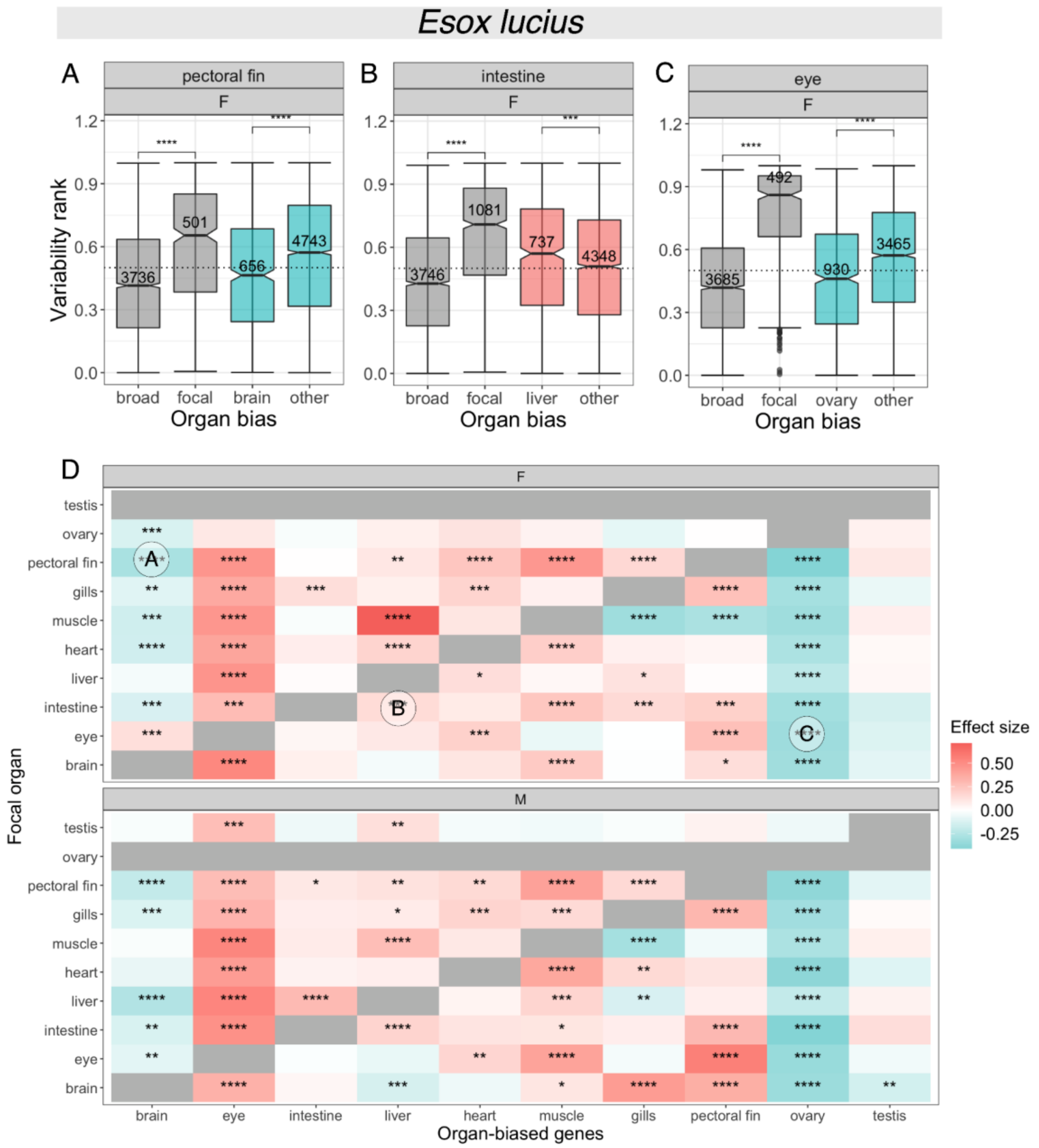


**Figure S33. Relationship between expression variability and organ-biased expression for Northern pike. (A)** Boxplot of expression variability ranks of genes expressed in female ('F') Northern pike pectoral fin, for sets of genes classified by organ bias and with an emphasis on brain-biased genes. Broadly expressed genes ('broad') have a measured organ expression specificity index (tau) of $\tau \leq 0.3$; genes with $\tau > 0.3$ are considered organ-biased. Among the set of organ-biased genes, genes are further classified into three categories: focal-biased genes ('focal') are preferentially expressed in the focal organ (i.e., pectoral fin), brain-biased genes ('brain') are preferentially expressed in the brain, and the remaining ('other') are preferentially expressed in other organs. The number of genes included are annotated for each category. Comparisons between broadly expressed and focal-biased genes, and between brain-biased and other organ-biased genes were performed using Wilcoxon rank-sum tests (*p*-value significance levels: '****': $p \leq 0.0001$, '***': $p \leq 0.001$, '**': $p \leq 0.01$, '*': $p \leq 0.05$, 'ns': $p > 0.05$). The dashed line indicates a variability rank of 0.5. **(B)** Boxplot of variability ranks of genes expressed in female Northern pike intestine, with genes classified by organ bias and with an emphasis on liver-biased genes. **(C)** Boxplot of variability ranks of genes expressed in female Northern pike eye, with genes classified by organ bias and with an emphasis on ovary-biased genes. **(D)** Heatmap summarizing pairwise comparisons of expression variability ranks between subsets of organ-biased genes for female ('F', top) and male ('M', bottom) individuals. Rows correspond to focal organ whereas columns correspond to subsets of organ-biased genes**.** Pairwise comparisons were performed using Wilcoxon rank-sum tests with Benjamini-Hochberg ('BH') correction for multiple testing (adjusted *p*-value < 0.05). Effect size was computed using Glass's delta (δ), with color scale shown in (L). Cells in gray are not included in the dataset. Three cells are annotated, corresponding to the following comparisons in boxplots (A-C): (A) brain-biased vs. other organ-biased genes expressed in female Northern pike pectoral fin, (B) liver-biased vs. other organ-biased biased genes expressed in female intestine, and (C) ovary-biased vs. other organ-biased genes expressed in female eye. A negative effect size entails lower expression variability relative to other organ-biased genes (A,C) (turquoise), whereas a positive effect size entails higher variability (B) (salmon).

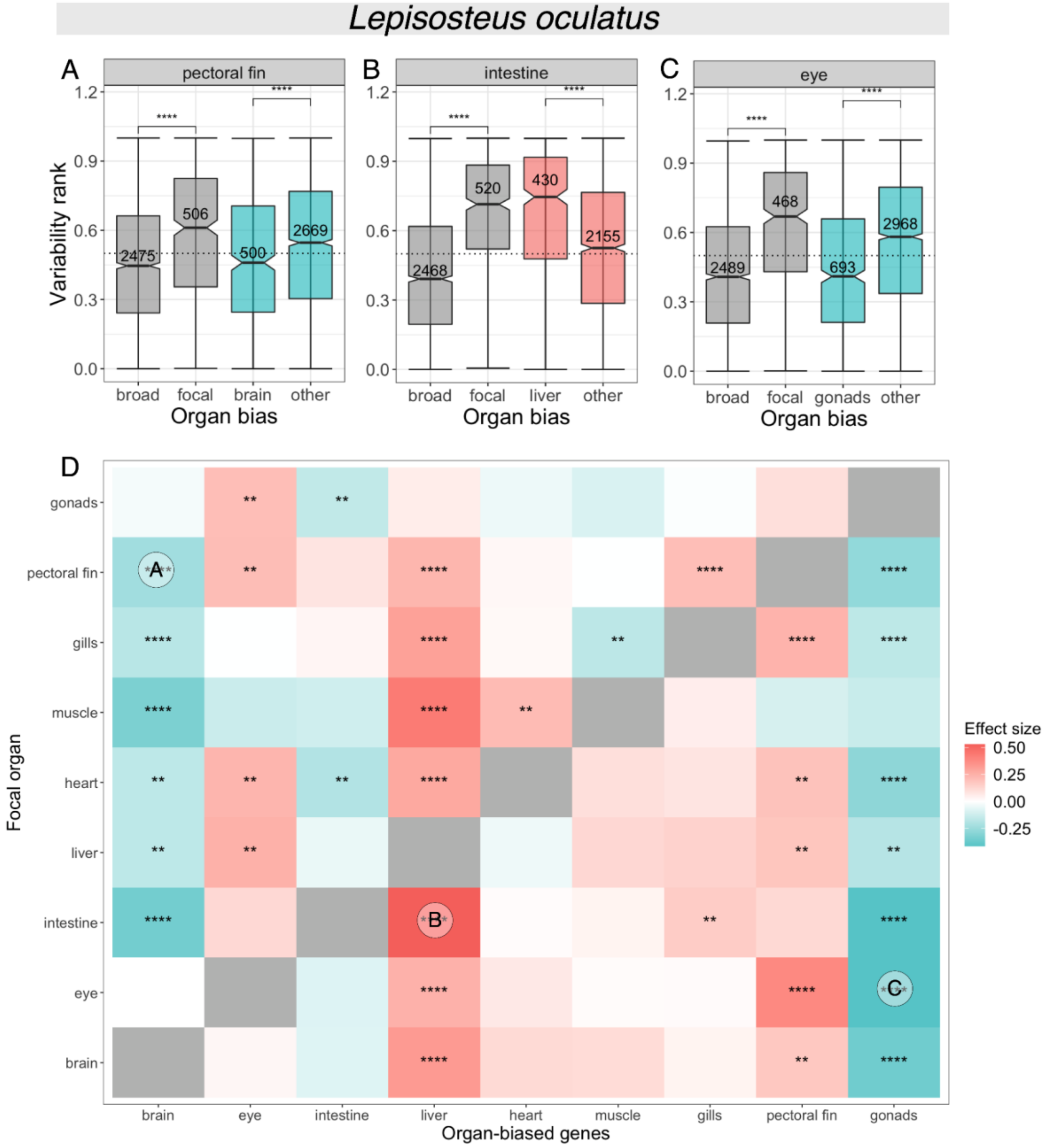


**Figure S34. Relationship between expression variability and organ-biased expression for spotted gar. (A)** Boxplot of expression variability ranks of genes expressed in spotted gar pectoral fin, for sets of genes classified by organ bias and with an emphasis on brain-biased genes. Broadly expressed genes ('broad') have a measured organ expression specificity index (tau) of $\tau \leq 0.3$; genes with $\tau > 0.3$ are considered organ-biased. Among the set of organ-biased genes, genes are further classified into three categories: focal-biased genes ('focal') are preferentially expressed in the focal organ (i.e., pectoral fin), brain-biased genes ('brain') are preferentially expressed in the brain, and the remaining ('other') are preferentially expressed in other organs. The number of genes included are annotated for each category. Comparisons between broadly expressed and focal-biased genes, and between brain-biased and other organ-biased genes were performed using Wilcoxon rank-sum tests ($p$-value significance levels: '****': $p \leq 0.0001$, '***': $p \leq 0.001$, '**': $p \leq 0.01$, '*': $p \leq 0.05$, 'ns': $p > 0.05$). The dashed line indicates a variability rank of 0.5. **(B)** Boxplot of variability ranks of genes expressed in spotted gar intestine, with genes classified by organ bias and with an emphasis on liver-biased genes. **(C)** Boxplot of variability ranks of genes expressed in spotted gar eye, with genes classified by organ bias and with an emphasis on gonads-biased genes. **(D)** Heatmap summarizing pairwise comparisons of expression variability ranks between subsets of organ-biased genes. Rows correspond to focal organ whereas columns correspond to subsets of organ-biased genes**.** Pairwise comparisons were performed using Wilcoxon rank-sum tests with Benjamini-Hochberg ('BH') correction for multiple testing (adjusted $p$-value < 0.05). Effect size was computed using Glass's delta ($\delta$), with color scale shown in (L). Cells in gray are not included in the dataset. Three cells are annotated, corresponding to the following comparisons in boxplots (A-C): (A) brain-biased vs. other organ-biased genes expressed in the pectoral fin, (B) liver-biased vs. other organ-biased biased genes expressed in the intestine, and (C) ovary-biased vs. other organ-biased genes expressed in the eye. A negative effect size entails lower expression variability relative to other organ-biased genes (A,C) (turquoise), whereas a positive effect size entails higher variability (B) (salmon).

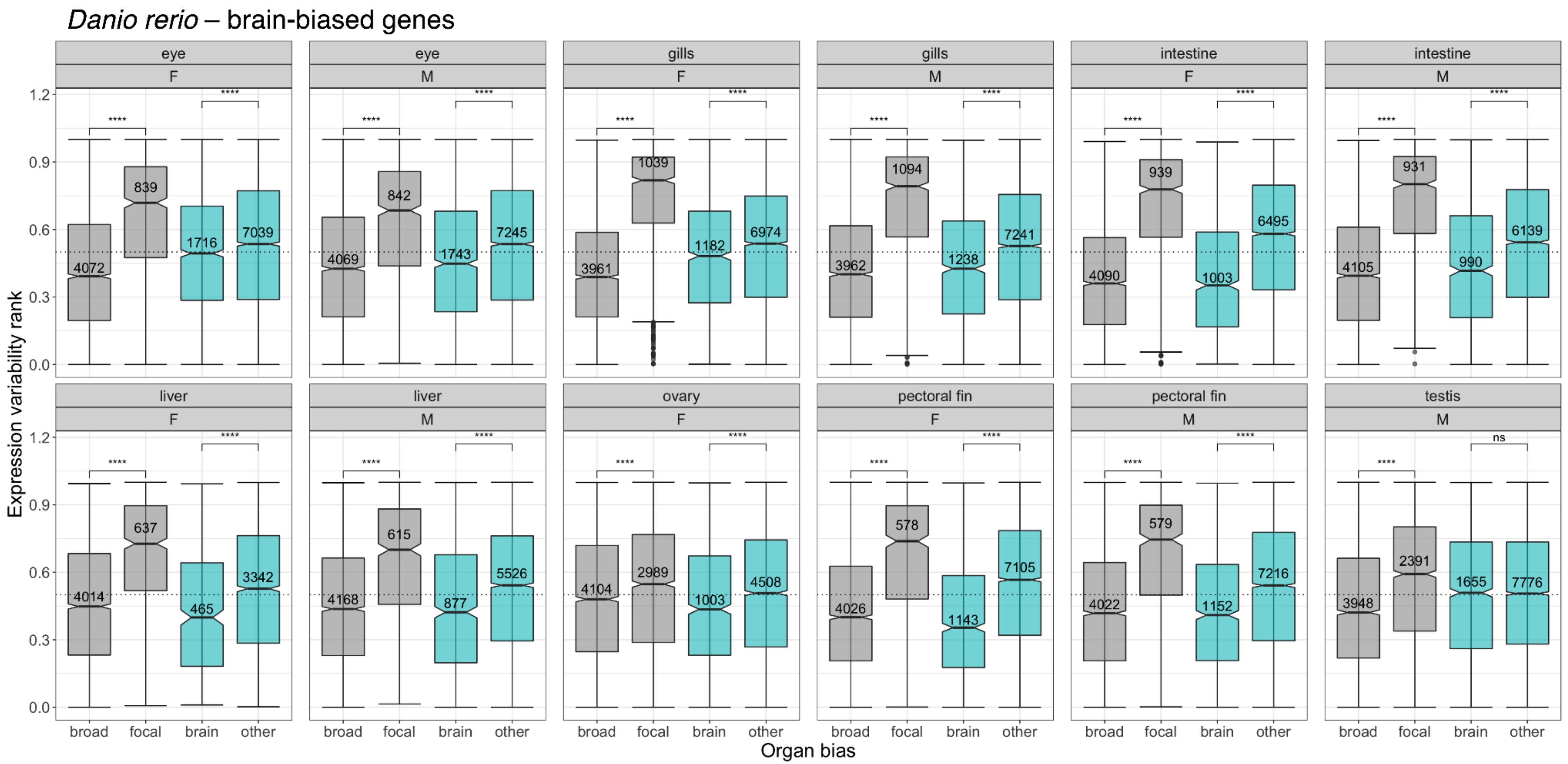


**Figure S35. Distribution of expression variability ranks across organ-sex conditions in zebrafish, for genes classified by organ bias and with an emphasis on brain-biased genes.** Broadly expressed genes ('broad') have a measured organ expression specificity index (tau) of $\tau \leq 0.3$; genes with $\tau > 0.3$ are considered organ-biased. Among the set of organ-biased genes, genes are further classified into three categories: focal-biased genes ('focal') are preferentially expressed in the indicated focal organ, brain-biased genes ('brain') are preferentially expressed in the brain, and the remaining ('other') are preferentially expressed in other organs. The number of genes included are annotated for each category. Comparisons between broadly expressed and focal-biased genes (gray boxplots), and between brain-biased and other organ-biased genes (turquoise boxplots) were performed using Wilcoxon rank-sum tests (*p*-value significance levels: '****': $p \leq 0.0001$, '***': $p \leq 0.001$, '**': $p \leq 0.01$, '*': $p \leq 0.05$, 'ns': $p > 0.05$). The dashed line indicates a variability rank of 0.5.

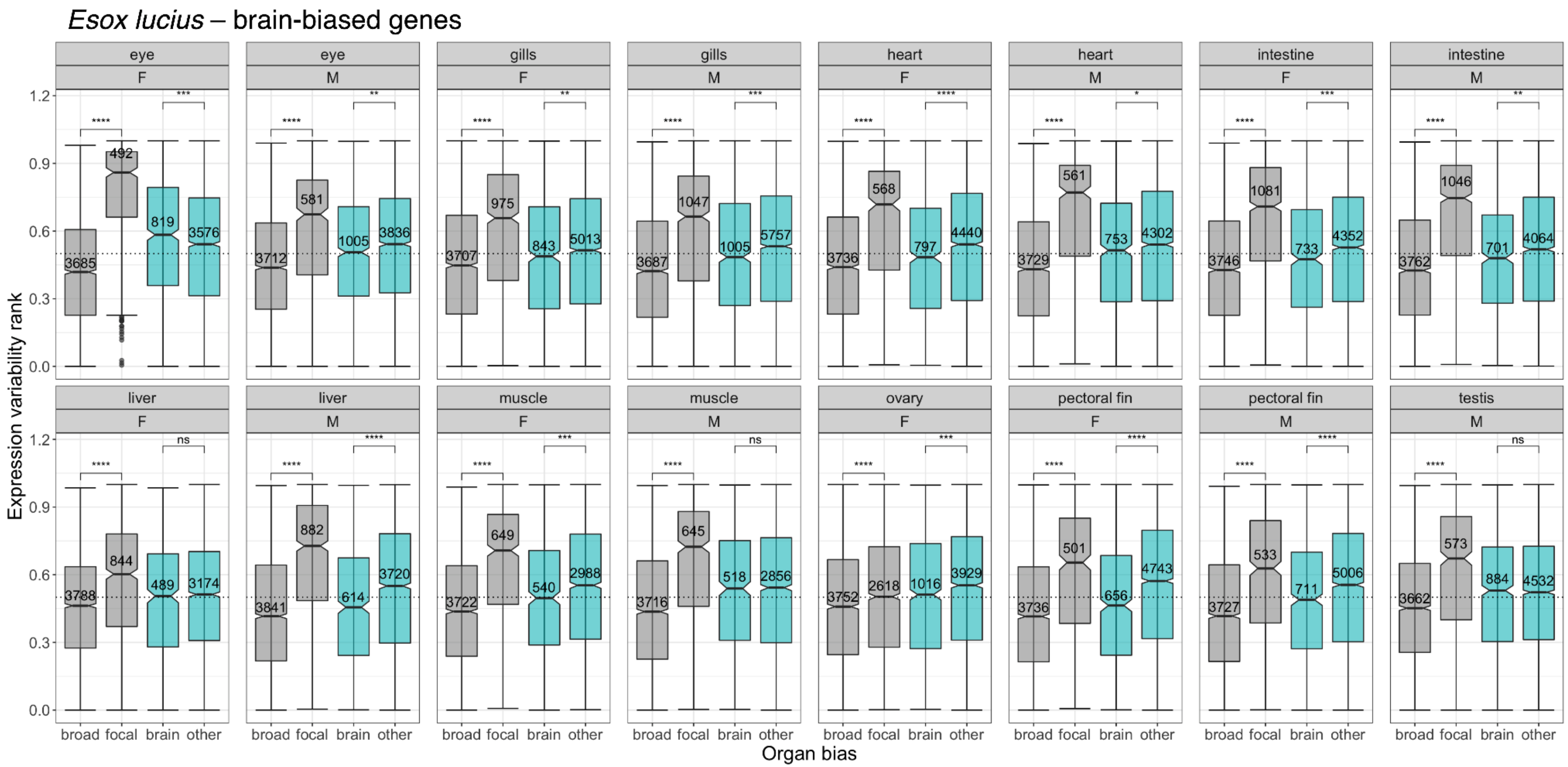


**Figure S36. Distribution of expression variability ranks across organ-sex conditions in Northern pike, for genes classified by organ bias and with an emphasis on brain-biased genes.** Broadly expressed genes ('broad') have a measured organ expression specificity index (tau) of $\tau \leq 0.3$; genes with $\tau > 0.3$ are considered organ-biased. Among the set of organ-biased genes, genes are further classified into three categories: focal-biased genes ('focal') are preferentially expressed in the indicated focal organ, brain-biased genes ('brain') are preferentially expressed in the brain, and the remaining ('other') are preferentially expressed in other organs. The number of genes included are annotated for each category. Comparisons between broadly expressed and focal-biased genes (gray boxplots), and between brain-biased and other organ-biased genes (turquoise boxplots) were performed using Wilcoxon rank-sum tests (*p*-value significance levels: '****': $p \leq 0.0001$, '***': $p \leq 0.001$, '**': $p \leq 0.01$, '*': $p \leq 0.05$, 'ns': $p > 0.05$). The dashed line indicates a variability rank of 0.5.

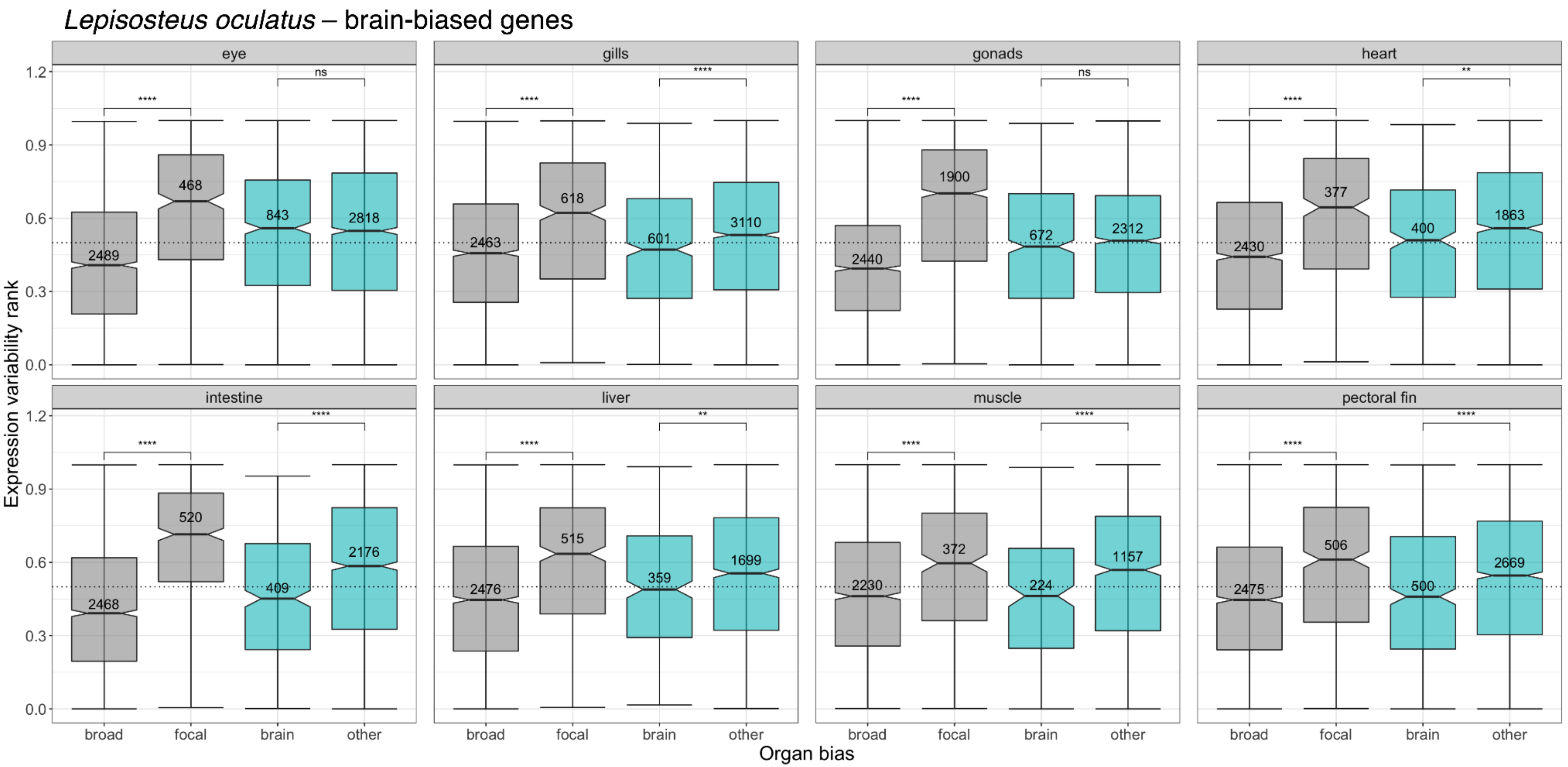


**Figure S37. Distribution of expression variability ranks across organs in spotted gar, for genes classified by organ bias and with an emphasis on brain-biased genes.** Broadly expressed genes ('broad') have a measured organ expression specificity index (tau) of $\tau \leq 0.3$; genes with $\tau > 0.3$ are considered organ-biased. Among the set of organ-biased genes, genes are further classified into three categories: focal-biased genes ('focal') are preferentially expressed in the indicated focal organ, brain-biased genes ('brain') are preferentially expressed in the brain, and the remaining ('other') are preferentially expressed in other organs. The number of genes included are annotated for each category. Comparisons between broadly expressed and focal-biased genes (gray boxplots), and between brain-biased and other organ-biased genes (turquoise boxplots) were performed using Wilcoxon rank-sum tests (*p*-value significance levels: '****': $p \leq 0.0001$, '***': $p \leq 0.001$, '**': $p \leq 0.01$, '*': $p \leq 0.05$, 'ns': $p > 0.05$). The dashed line indicates a variability rank of 0.5.

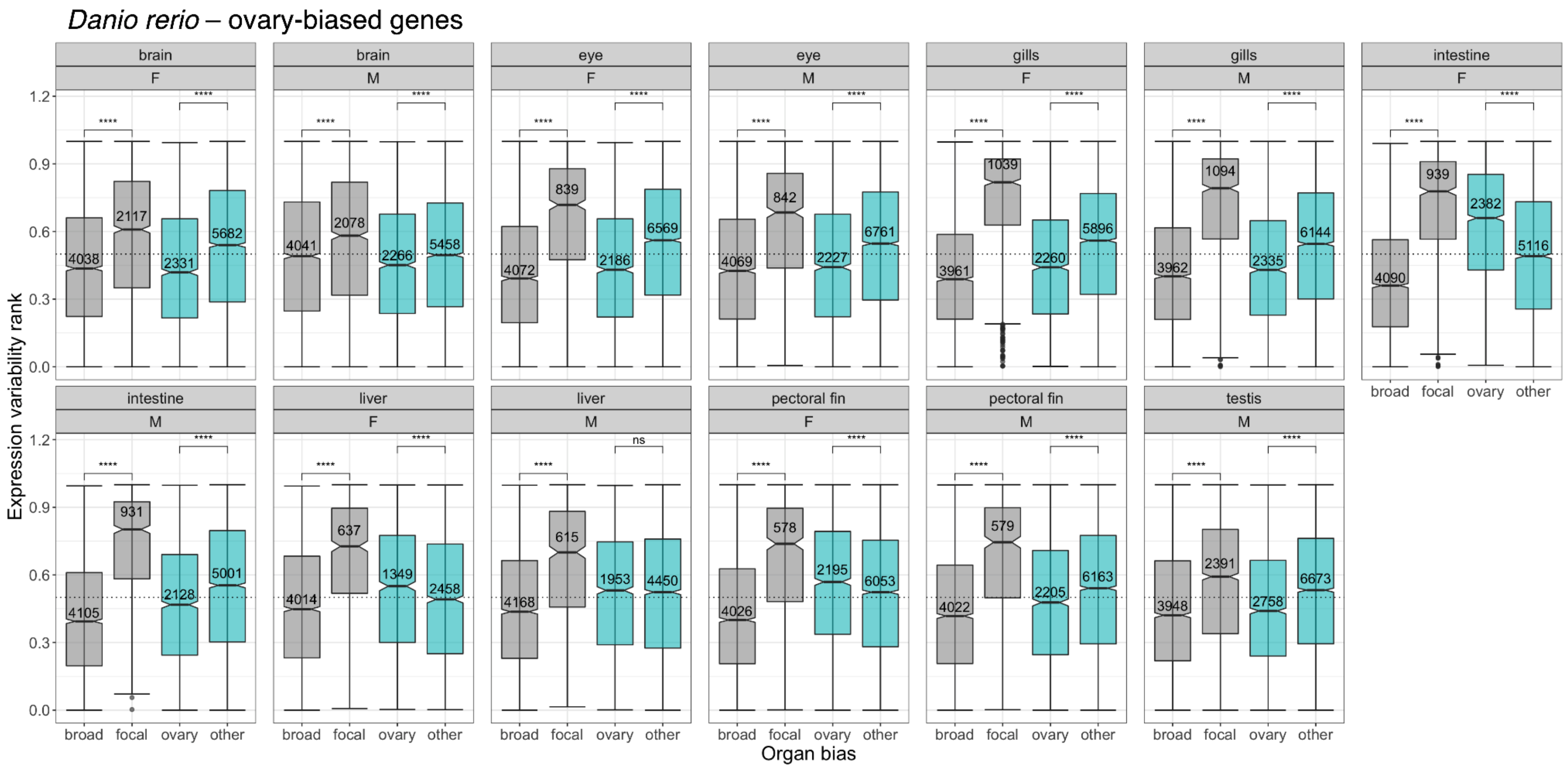


**Figure S38. Distribution of expression variability ranks across organ-sex conditions in zebrafish, for genes classified by organ bias and with an emphasis on ovary-biased genes.** Broadly expressed genes ('broad') have a measured organ expression specificity index (tau) of $\tau \leq 0.3$; genes with $\tau > 0.3$ are considered organ-biased. Among the set of organ-biased genes, genes are further classified into three categories: focal-biased genes ('focal') are preferentially expressed in the indicated focal organ, ovary-biased genes ('ovary') are preferentially expressed in the ovary, and the remaining ('other') are preferentially expressed in other organs. The number of genes included are annotated for each category. Comparisons between broadly expressed and focal-biased genes (gray boxplots), and between ovary-biased and other organ-biased genes (turquoise boxplots) were performed using Wilcoxon rank-sum tests ($p$-value significance levels: '****': $p \leq 0.0001$, '***': $p \leq 0.001$, '**': $p \leq 0.01$, '*': $p \leq 0.05$, 'ns': $p > 0.05$). The dashed line indicates a variability rank of 0.5.

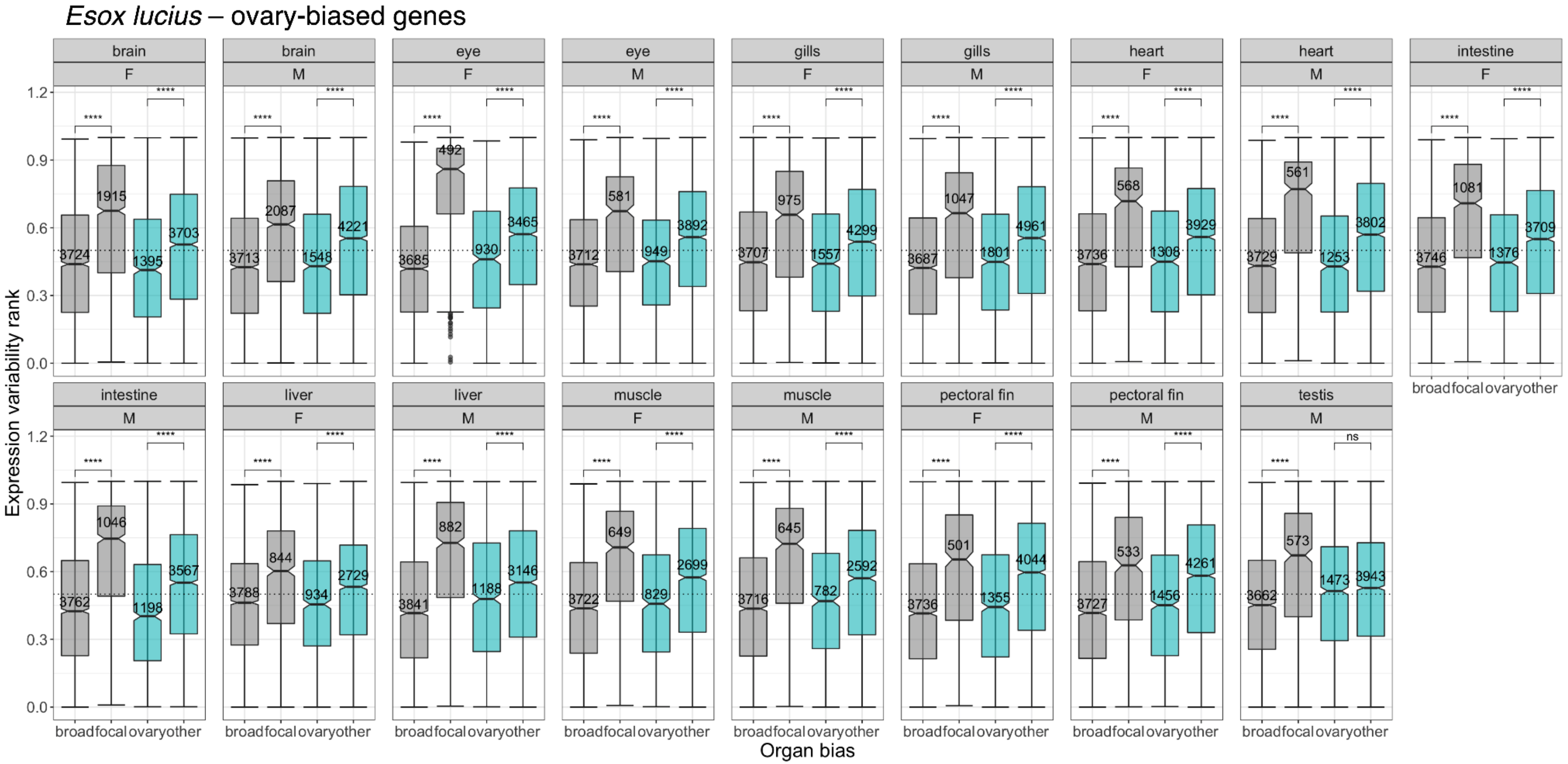


**Figure S39. Distribution of expression variability ranks across organ-sex conditions in Northern pike, for genes classified by organ bias and with an emphasis on ovary-biased genes.** Broadly expressed genes ('broad') have a measured organ expression specificity index (tau) of $\tau \leq 0.3$; genes with $\tau > 0.3$ are considered organ-biased. Among the set of organ-biased genes, genes are further classified into three categories: focal-biased genes ('focal') are preferentially expressed in the indicated focal organ, ovary-biased genes ('ovary') are preferentially expressed in the ovary, and the remaining ('other') are preferentially expressed in other organs. The number of genes included are annotated for each category. Comparisons between broadly expressed and focal-biased genes (gray boxplots), and between ovary-biased and other organ-biased genes (turquoise boxplots) were performed using Wilcoxon rank-sum tests ($p$-value significance levels: '****': $p \leq 0.0001$, '***': $p \leq 0.001$, '**': $p \leq 0.01$, '*': $p \leq 0.05$, 'ns': $p > 0.05$). The dashed line indicates a variability rank of 0.5.

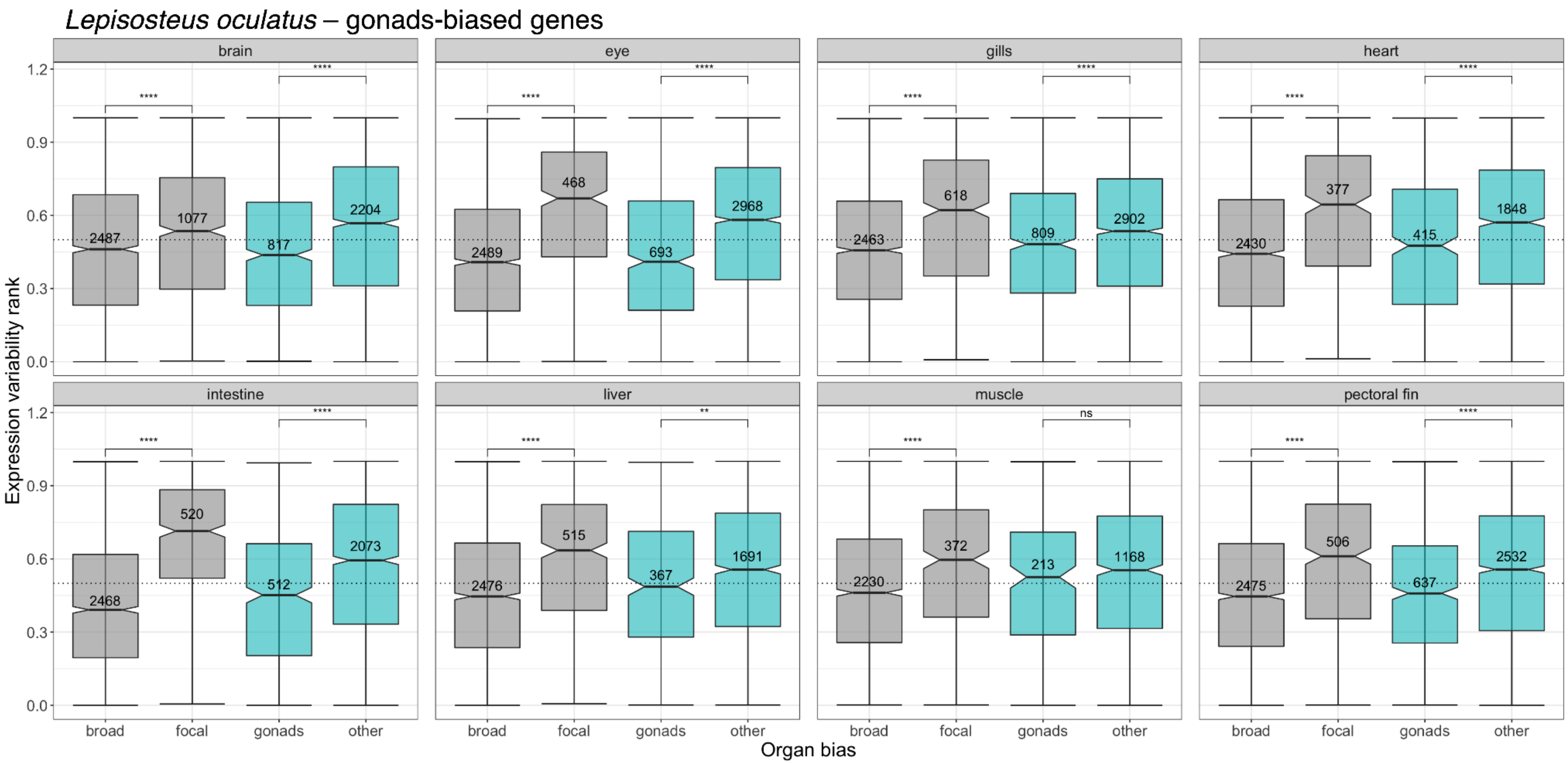


**Figure S40. Distribution of expression variability ranks across organs in spotted gar, for genes classified by organ bias and with an emphasis on gonads-biased genes.** Broadly expressed genes ('broad') have a measured organ expression specificity index (tau) of $\tau \leq 0.3$; genes with $\tau > 0.3$ are considered organ-biased. Among the set of organ-biased genes, genes are further classified into three categories: focal-biased genes ('focal') are preferentially expressed in the indicated focal organ, gonads-biased genes ('gonads') are preferentially expressed in the gonads, and the remaining ('other') are preferentially expressed in other organs. The number of genes included are annotated for each category. Comparisons between broadly expressed and focal-biased genes (gray boxplots), and between gonads-biased and other organ-biased genes (turquoise boxplots) were performed using Wilcoxon rank-sum tests ($p$-value significance levels: '****': $p \leq 0.0001$, '***': $p \leq 0.001$, '**': $p \leq 0.01$, '*': $p \leq 0.05$, 'ns': $p > 0.05$). The dashed line indicates a variability rank of 0.5.

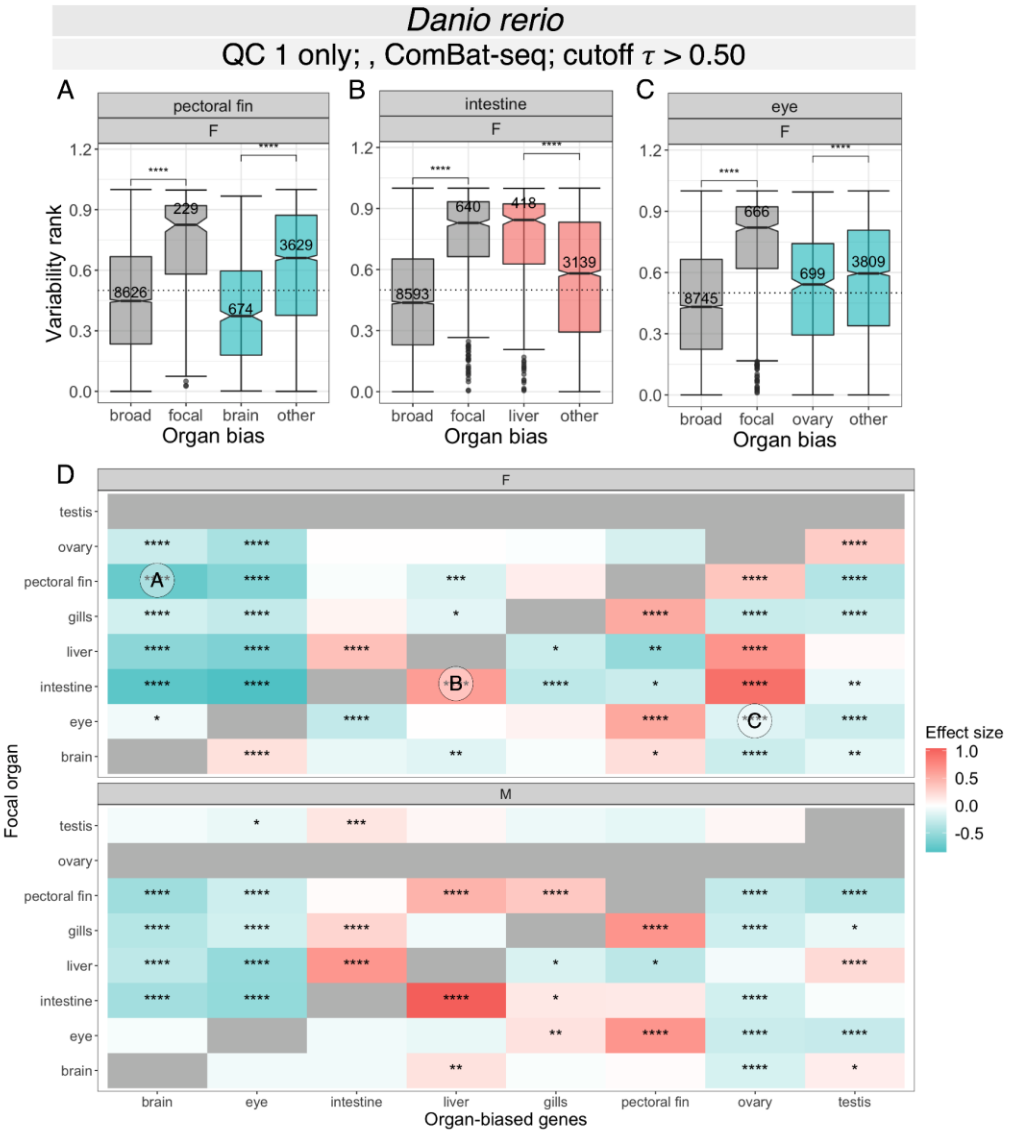


**Figure S41. Relationship between expression variability and organ-biased expression for zebrafish.** To assess the influence of sample filtering criteria on the analysis, samples were filtered based only on sequencing quality ("QC 1 only") (**Figure S3**) rather than the two-step filtering protocol described in Materials and Methods. Batch correction with *ComBat-seq* (Zhang et al. 2020) was performed prior to gene filtering, normalization, expression specificity and variability analysis. **(A)** Boxplot of expression variability ranks of genes expressed in female ('F') zebrafish pectoral fin, for sets of genes classified by organ bias and with an emphasis on brain-biased genes. Broadly expressed genes ('broad') have a measured organ expression specificity index (tau) of $\tau \leq 0.5$; genes with $\tau > 0.5$ are considered organ-biased. Among the set of organ-biased genes, genes are further classified into three categories: focal-biased genes ('focal') are preferentially expressed in the focal organ (i.e., pectoral fin), brain-biased genes ('brain') are preferentially expressed in the brain, and the remaining ('other') are preferentially expressed in other organs. The number of genes included are annotated for each category. Comparisons between broadly expressed and focal-biased genes, and between brain-biased and other organ-biased genes were performed using Wilcoxon rank-sum tests (*p*-value significance levels: '****': $p \leq 0.0001$, '***': $p \leq 0.001$, '**': $p \leq 0.01$, '*': $p \leq 0.05$, 'ns': $p > 0.05$). The dashed line indicates a variability rank of 0.5. **(B)** Boxplot of variability ranks of genes expressed in female zebrafish intestine, with genes classified by organ bias and with an emphasis on liver-biased genes. **(C)** Boxplot of variability ranks of genes expressed in female zebrafish eye, with genes classified by organ bias and with an emphasis on ovary-biased genes. **(D)** Heatmap summarizing pairwise comparisons of expression variability ranks between subsets of organ-biased genes for female ('F', top) and male ('M', bottom) zebrafish. Rows correspond to focal organ whereas columns correspond to subsets of organ-biased genes**.** Pairwise comparisons were performed using Wilcoxon rank-sum tests with Benjamini-Hochberg ('BH') correction for multiple testing (adjusted *p*-value < 0.05). Effect size was computed using Glass's delta (δ), with color scale shown in (L). Cells in gray are not included in the dataset. Three cells are annotated, corresponding to the following comparisons in boxplots (A-C): (A) brain-biased vs. other organ-biased genes expressed in female zebrafish pectoral fin, (B) liver-biased vs. other organ-biased biased genes expressed in female zebrafish intestine, and (C) ovary-biased vs. other organ-biased genes expressed in female zebrafish eye. A negative effect size entails lower expression variability relative to other organ-biased genes (A,C) (turquoise), whereas a positive effect size entails higher variability (B) (salmon).

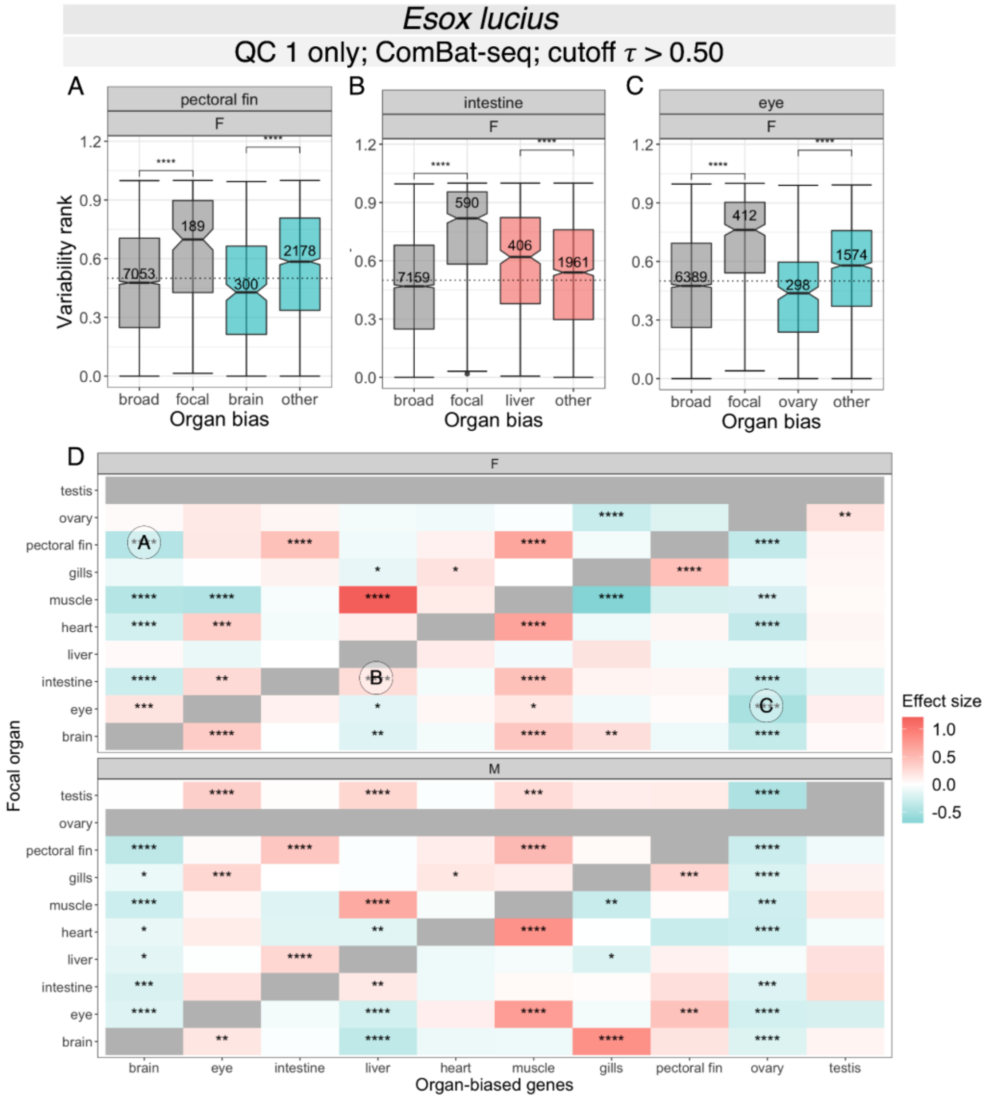


**Figure S42. Relationship between expression variability and organ-biased expression for Northern pike.** To assess the influence of sample filtering criteria on the analysis, samples were filtered based only on sequencing quality ("QC 1 only") (**Figure S2**) rather than the two-step filtering protocol described in Materials and Methods. Batch correction with *ComBat-seq* (Zhang et al. 2020) was performed prior to gene filtering, normalization, expression specificity and variability analysis. **(A)** Boxplot of expression variability ranks of genes expressed in female ('F') Northern pike pectoral fin, for sets of genes classified by organ bias and with an emphasis on brain-biased genes. Broadly expressed genes ('broad') have a measured organ expression specificity index (tau) of $\tau \leq 0.5$; genes with $\tau > 0.5$ are considered organ-biased. Among the set of organ-biased genes, genes are further classified into three categories: focal-biased genes ('focal') are preferentially expressed in the focal organ (i.e., pectoral fin), brain-biased genes ('brain') are preferentially expressed in the brain, and the remaining ('other') are preferentially expressed in other organs. The number of genes included are annotated for each category. Comparisons between broadly expressed and focal-biased genes, and between brain-biased and other organ-biased genes were performed using Wilcoxon rank-sum tests (*p*-value significance levels: '****': $p \leq 0.0001$, '***': $p \leq 0.001$, '**': $p \leq 0.01$, '*': $p \leq 0.05$, 'ns': $p > 0.05$). The dashed line indicates a variability rank of 0.5. **(B)** Boxplot of variability ranks of genes expressed in female Northern pike intestine, with genes classified by organ bias and with an emphasis on liver-biased genes. **(C)** Boxplot of variability ranks of genes expressed in female Northern pike eye, with genes classified by organ bias and with an emphasis on ovary-biased genes. **(D)** Heatmap summarizing pairwise comparisons of expression variability ranks between subsets of organ-biased genes for female ('F', top) and male ('M', bottom) individuals. Rows correspond to focal organ whereas columns correspond to subsets of organ-biased genes**.** Pairwise comparisons were performed using Wilcoxon rank-sum tests with Benjamini-Hochberg ('BH') correction for multiple testing (adjusted *p*-value < 0.05). Effect size was computed using Glass's delta (δ), with color scale shown in (L). Cells in gray are not included in the dataset. Three cells are annotated, corresponding to the following comparisons in boxplots (A-C): (A) brain-biased vs. other organ-biased genes expressed in female Northern pike pectoral fin, (B) liver-biased vs. other organ-biased biased genes expressed in female intestine, and (C) ovary-biased vs. other organ-biased genes expressed in female eye. A negative effect size entails lower expression variability relative to other organ-biased genes (A,C) (turquoise), whereas a positive effect size entails higher variability (B) (salmon).

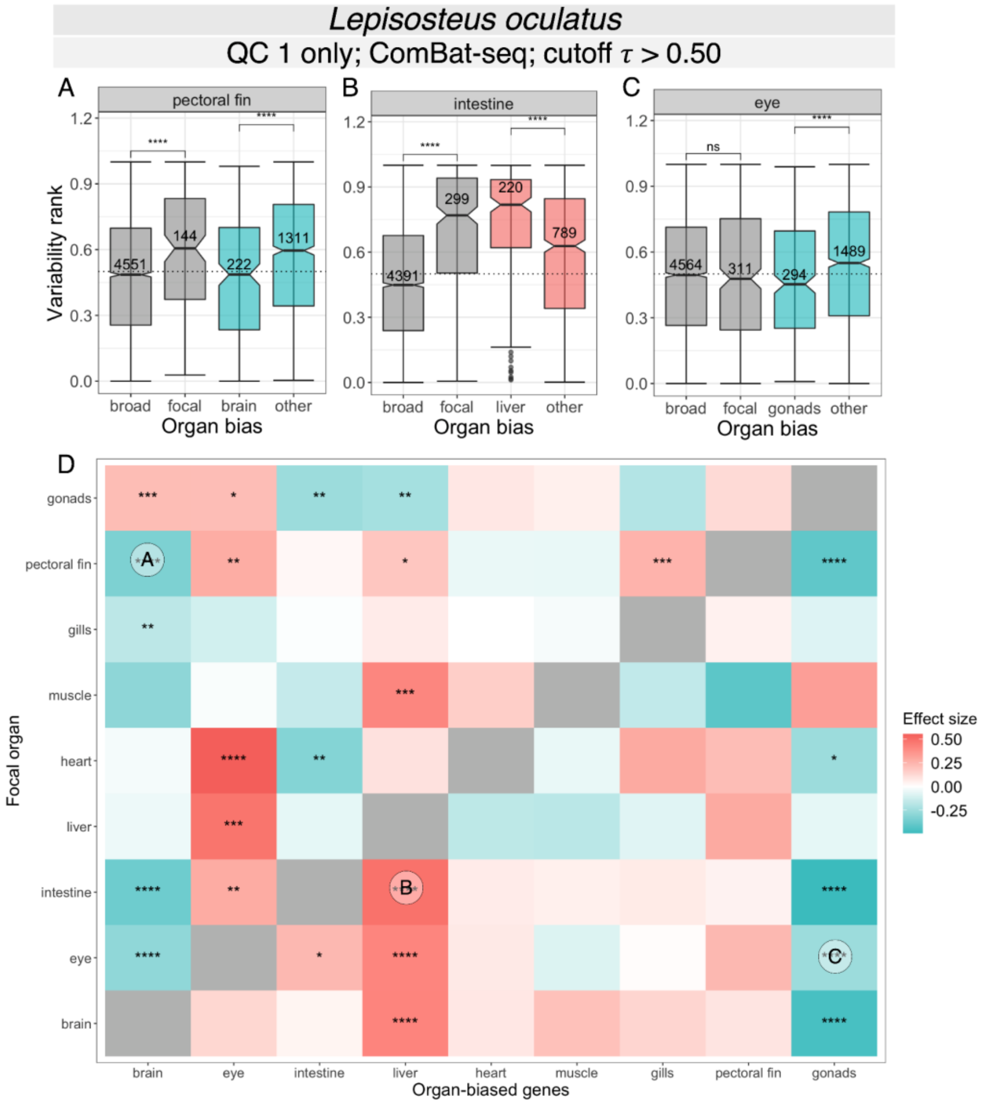


**Figure S43. Relationship between expression variability and organ-biased expression for spotted gar.** To assess the influence of sample filtering criteria on the analysis, samples were filtered based only on sequencing quality ("QC 1 only") (**Figure S1**) rather than the two-step filtering protocol described in Materials and Methods. Batch correction with *ComBat-seq* (Zhang et al. 2020) was performed prior to gene filtering, normalization, expression specificity and variability analysis. **(A)** Boxplot of expression variability ranks of genes expressed in spotted gar pectoral fin, for sets of genes classified by organ bias and with an emphasis on brain-biased genes. Broadly expressed genes ('broad') have a measured organ expression specificity index (tau) of $\tau \leq 0.5$; genes with $\tau > 0.5$ are considered organ-biased. Among the set of organ-biased genes, genes are further classified into three categories: focal-biased genes ('focal') are preferentially expressed in the focal organ (i.e., pectoral fin), brain-biased genes ('brain') are preferentially expressed in the brain, and the remaining ('other') are preferentially expressed in other organs. The number of genes included are annotated for each category. Comparisons between broadly expressed and focal-biased genes, and between brain-biased and other organ-biased genes were performed using Wilcoxon rank-sum tests (*p*-value significance levels: '****': $p \leq 0.0001$, '***': $p \leq 0.001$, '**': $p \leq 0.01$, '*': $p \leq 0.05$, 'ns': $p > 0.05$). The dashed line indicates a variability rank of 0.5. **(B)** Boxplot of variability ranks of genes expressed in spotted gar intestine, with genes classified by organ bias and with an emphasis on liver-biased genes. **(C)** Boxplot of variability ranks of genes expressed in spotted gar eye, with genes classified by organ bias and with an emphasis on gonads-biased genes. **(D)** Heatmap summarizing pairwise comparisons of expression variability ranks between subsets of organ-biased genes. Rows correspond to focal organ whereas columns correspond to subsets of organ-biased genes**.** Pairwise comparisons were performed using Wilcoxon rank-sum tests with Benjamini-Hochberg ('BH') correction for multiple testing (adjusted *p*-value < 0.05). Effect size was computed using Glass's delta (δ), with color scale shown in (L). Cells in gray are not included in the dataset. Three cells are annotated, corresponding to the following comparisons in boxplots (A-C): (A) brain-biased vs. other organ-biased genes expressed in the pectoral fin, (B) liver-biased vs. other organ-biased biased genes expressed in the intestine, and (C) ovary-biased vs. other organ-biased genes expressed in the eye. A negative effect size entails lower expression variability relative to other organ-biased genes (A,C) (turquoise), whereas a positive effect size entails higher variability (B) (salmon).

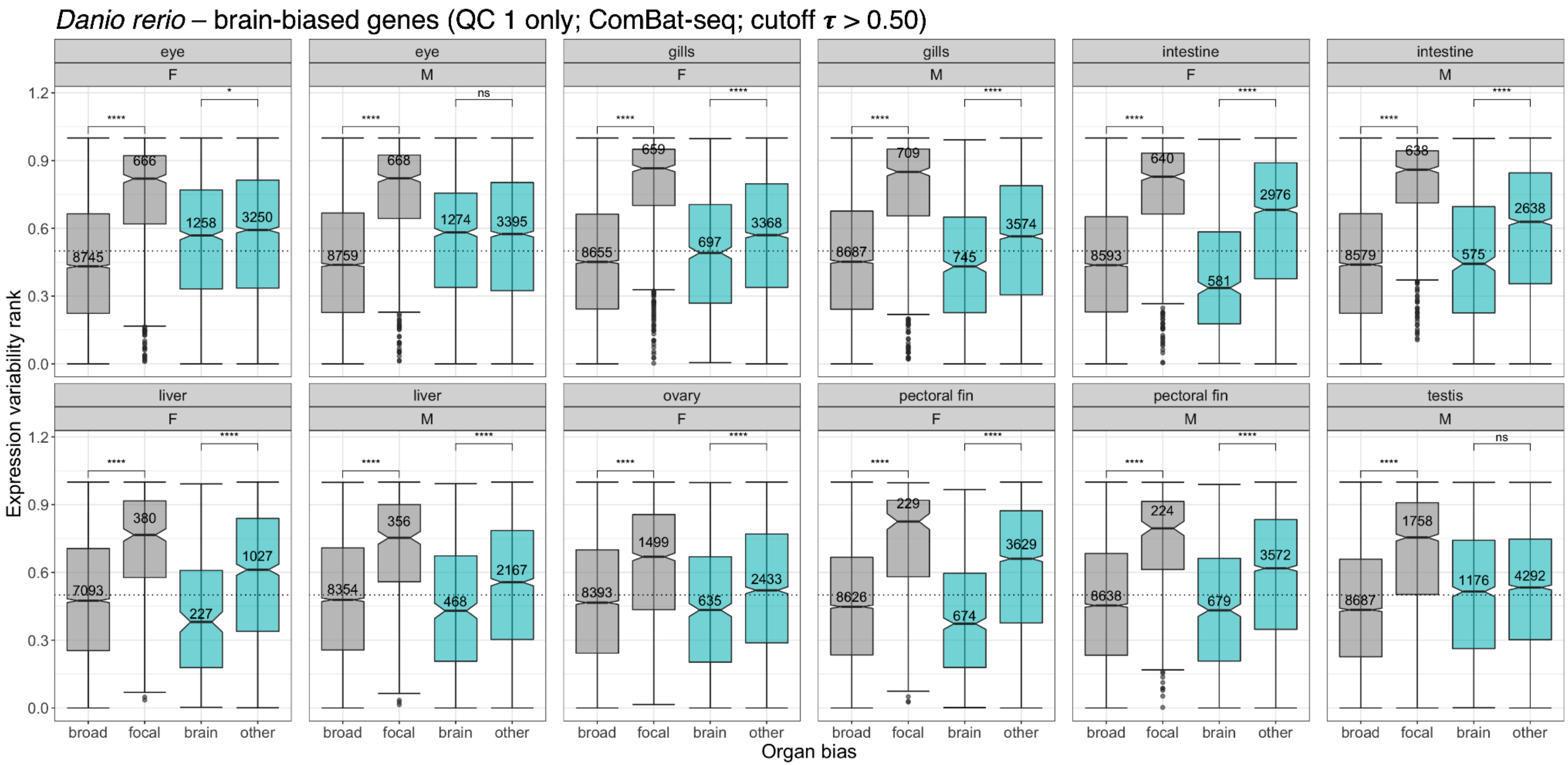


**Figure S44. Distribution of expression variability ranks across organ-sex conditions in zebrafish, for genes classified by organ bias and with an emphasis on brain-biased genes.** To assess the influence of sample filtering criteria on the analysis, samples were filtered based only on sequencing quality ("QC 1 only") (**Figure S3**) rather than the two-step filtering protocol. Batch correction with *ComBat-seq* **(Zhang et al. 2020)** was performed prior to gene filtering, normalization, expression specificity and variability analysis. Broadly expressed genes ('broad') have a measured organ expression specificity index (tau) of $\tau \leq 0.5$; genes with $\tau > 0.5$ are considered organ-biased. Among the set of organ-biased genes, genes are further classified into three categories: focal-biased genes ('focal') are preferentially expressed in the indicated focal organ, brain-biased genes ('brain') are preferentially expressed in the brain, and the remaining ('other') are preferentially expressed in other organs. The number of genes included are annotated for each category. Comparisons between broadly expressed and focal-biased genes (gray boxplots), and between brain-biased and other organ-biased genes (turquoise boxplots) were performed using Wilcoxon rank-sum tests ($p$-value significance levels: '****': $p \leq 0.0001$, '***': $p \leq 0.001$, '**': $p \leq 0.01$, '*': $p \leq 0.05$, 'ns': $p > 0.05$). The dashed line indicates a variability rank of 0.5.

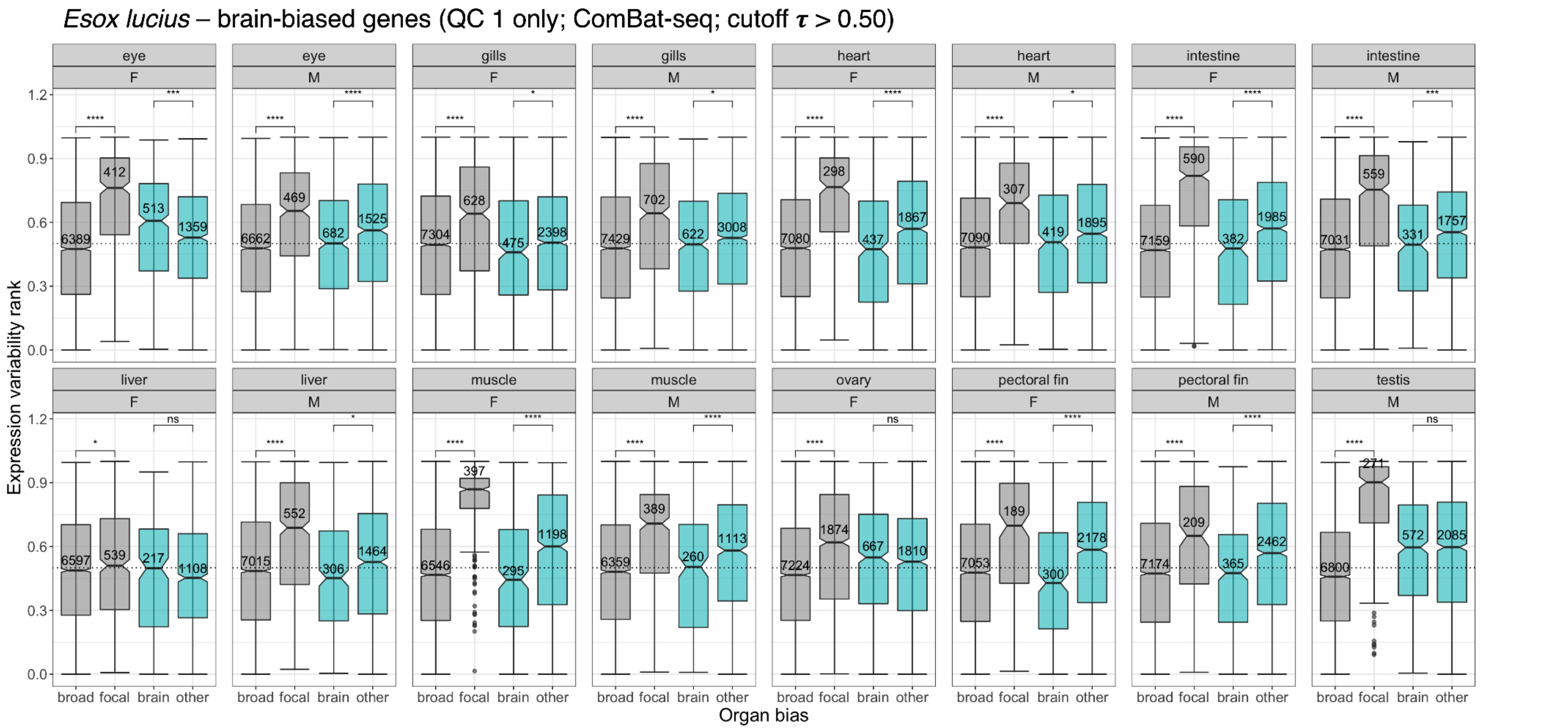


**Figure S45. Distribution of expression variability ranks across organ-sex conditions in Northern pike, for genes classified by organ bias and with an emphasis on brain-biased genes.** To assess the influence of sample filtering criteria on the analysis, samples were filtered based only on sequencing quality ("QC 1 only") (**Figure S2**) rather than the two-step filtering protocol. Batch correction with *ComBat-seq* (Zhang et al. 2020) was performed prior to gene filtering, normalization, expression specificity and variability analysis. Broadly expressed genes ('broad') have a measured organ expression specificity index (tau) of $\tau \leq 0.5$; genes with $\tau > 0.5$ are considered organ-biased. Among the set of organ-biased genes, genes are further classified into three categories: focal-biased genes ('focal') are preferentially expressed in the indicated focal organ, brain-biased genes ('brain') are preferentially expressed in the brain, and the remaining ('other') are preferentially expressed in other organs. The number of genes included are annotated for each category. Comparisons between broadly expressed and focal-biased genes (gray boxplots), and between brain-biased and other organ-biased genes (turquoise boxplots) were performed using Wilcoxon rank-sum tests (*p*-value significance levels: '****': $p \leq 0.0001$, '***': $p \leq 0.001$, '**': $p \leq 0.01$, '*': $p \leq 0.05$, 'ns': $p > 0.05$). The dashed line indicates a variability rank of 0.5.

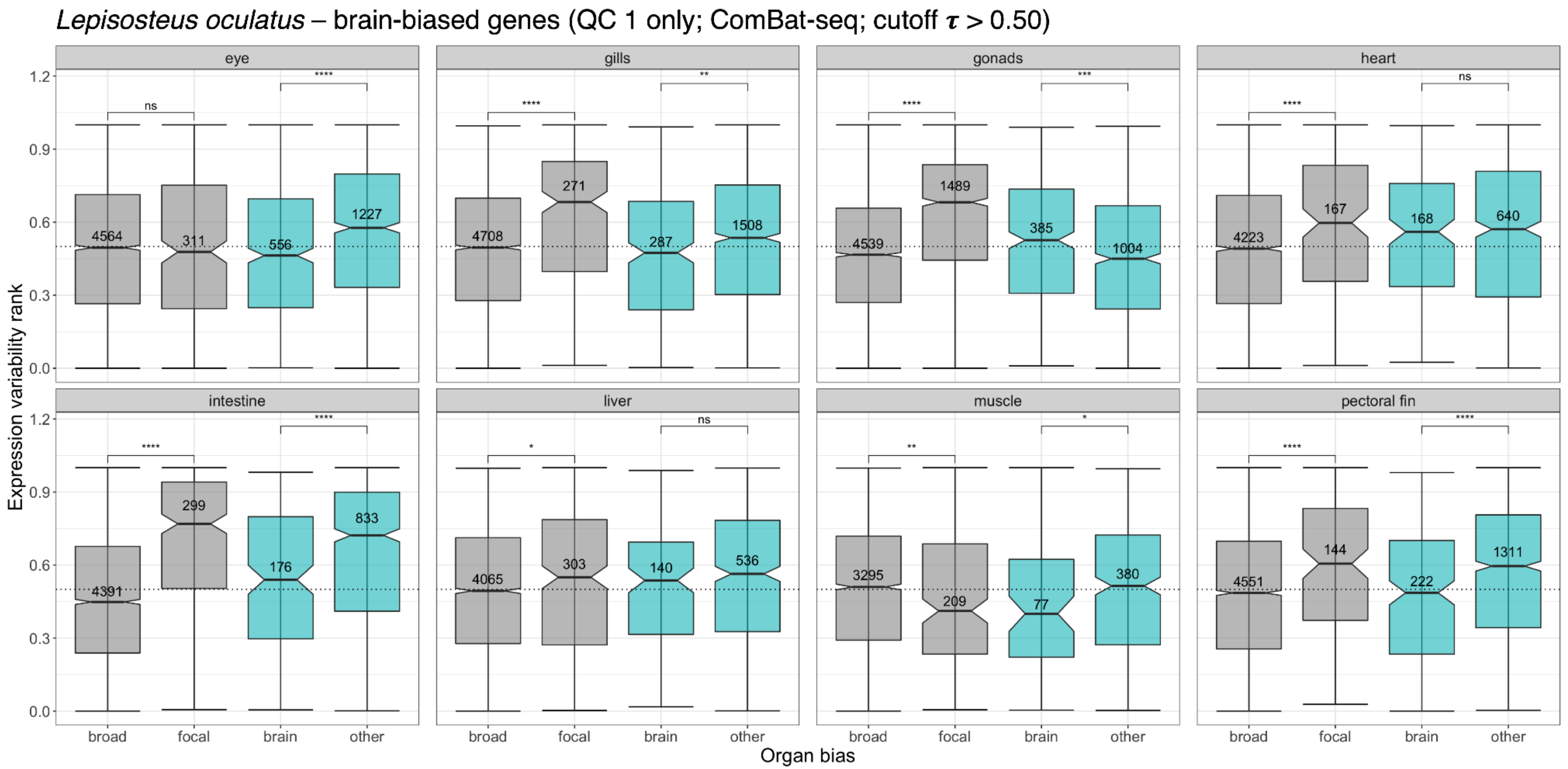


**Figure S46. Distribution of expression variability ranks across organs in spotted gar, for genes classified by organ bias and with an emphasis on brain-biased genes.** To assess the influence of sample filtering criteria on the analysis, samples were filtered based only on sequencing quality ("QC 1 only") (**Figure S1**) rather than the two-step filtering protocol. Batch correction with *ComBat-seq* (Zhang et al. 2020) was performed prior to gene filtering, normalization, expression specificity and variability analysis. Broadly expressed genes ('broad') have a measured organ expression specificity index (tau) of $\tau \leq 0.5$; genes with $\tau > 0.5$ are considered organ-biased. Among the set of organ-biased genes, genes are further classified into three categories: focal-biased genes ('focal') are preferentially expressed in the indicated focal organ, brain-biased genes ('brain') are preferentially expressed in the brain, and the remaining ('other') are preferentially expressed in other organs. The number of genes included are annotated for each category. Comparisons between broadly expressed and focal-biased genes (gray boxplots), and between brain-biased and other organ-biased genes (turquoise boxplots) were performed using Wilcoxon rank-sum tests (*p*-value significance levels: '****': $p \leq 0.0001$, '***': $p \leq 0.001$, '**': $p \leq 0.01$, '*': $p \leq 0.05$, 'ns': $p > 0.05$). The dashed line indicates a variability rank of 0.5.

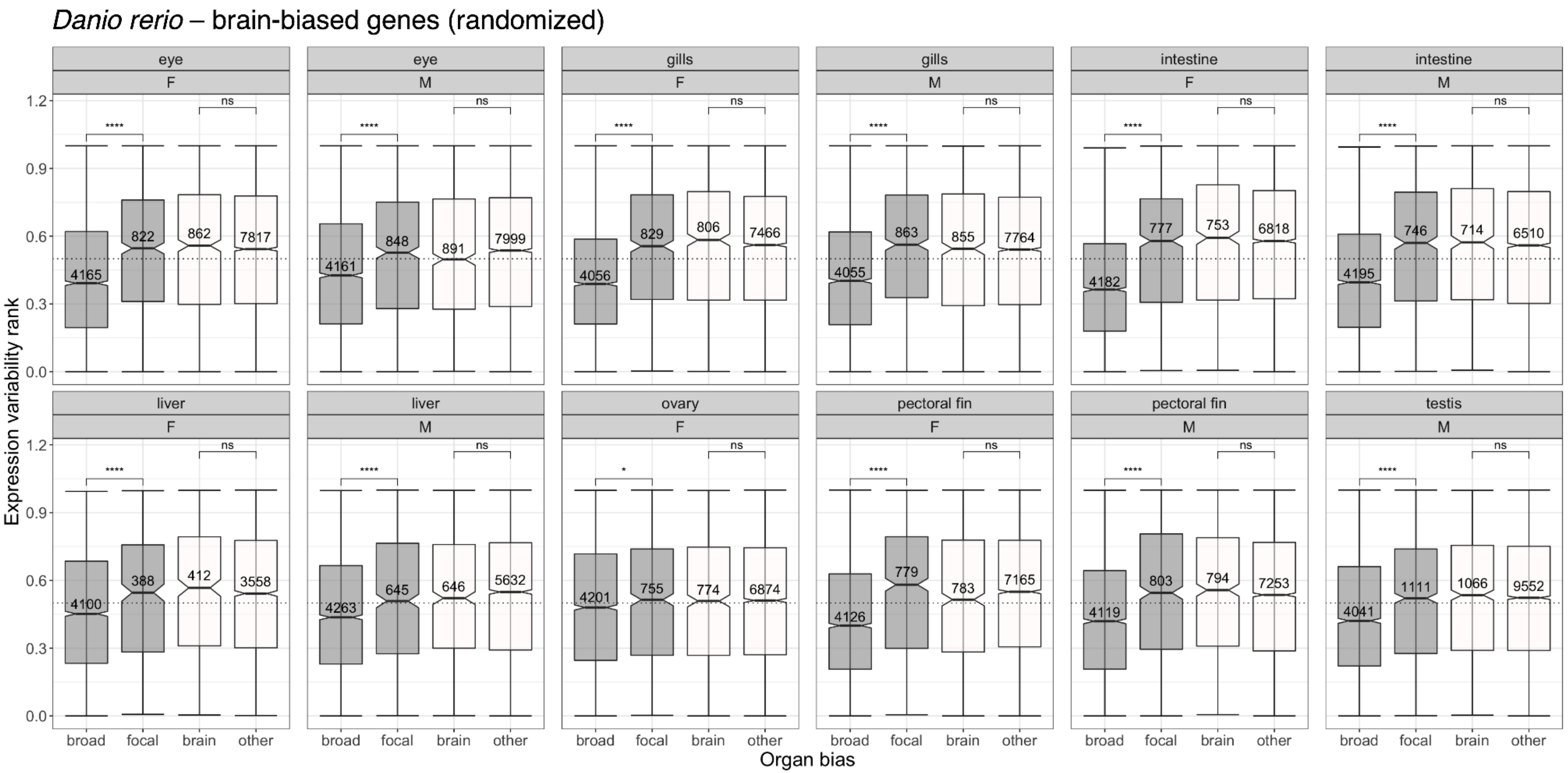


**Figure S47. Distribution of expression variability ranks across organ-sex conditions in zebrafish given randomized organ-biased genes and with an emphasis on 'brain-biased' genes.** For each gene, the top organ is assigned randomly with seed of 12345 without changing the expression specificity (tau) index. Genes with $\tau \leq 0.3$ are considered broadly expressed ('broad'), whereas genes with $\tau > 0.3$ are classified as organ-biased. Among the set of organ-biased genes, genes are further classified into three categories: focal-biased genes ('focal') are preferentially expressed in the indicated focal organ, brain-biased genes ('brain') are preferentially expressed in the brain, and the remaining ('other') are preferentially expressed in other organs. The number of genes included are annotated for each category. Comparisons between broadly expressed and focal-biased genes (gray boxplots), and between brain-biased and other organ-biased genes (white boxplots) were performed using Wilcoxon rank-sum tests (*p*-value significance levels: '****': $p \leq 0.0001$, '***': $p \leq 0.001$, '**': $p \leq 0.01$, '*': $p \leq 0.05$, 'ns': $p > 0.05$). The dashed line indicates a variability rank of 0.5.

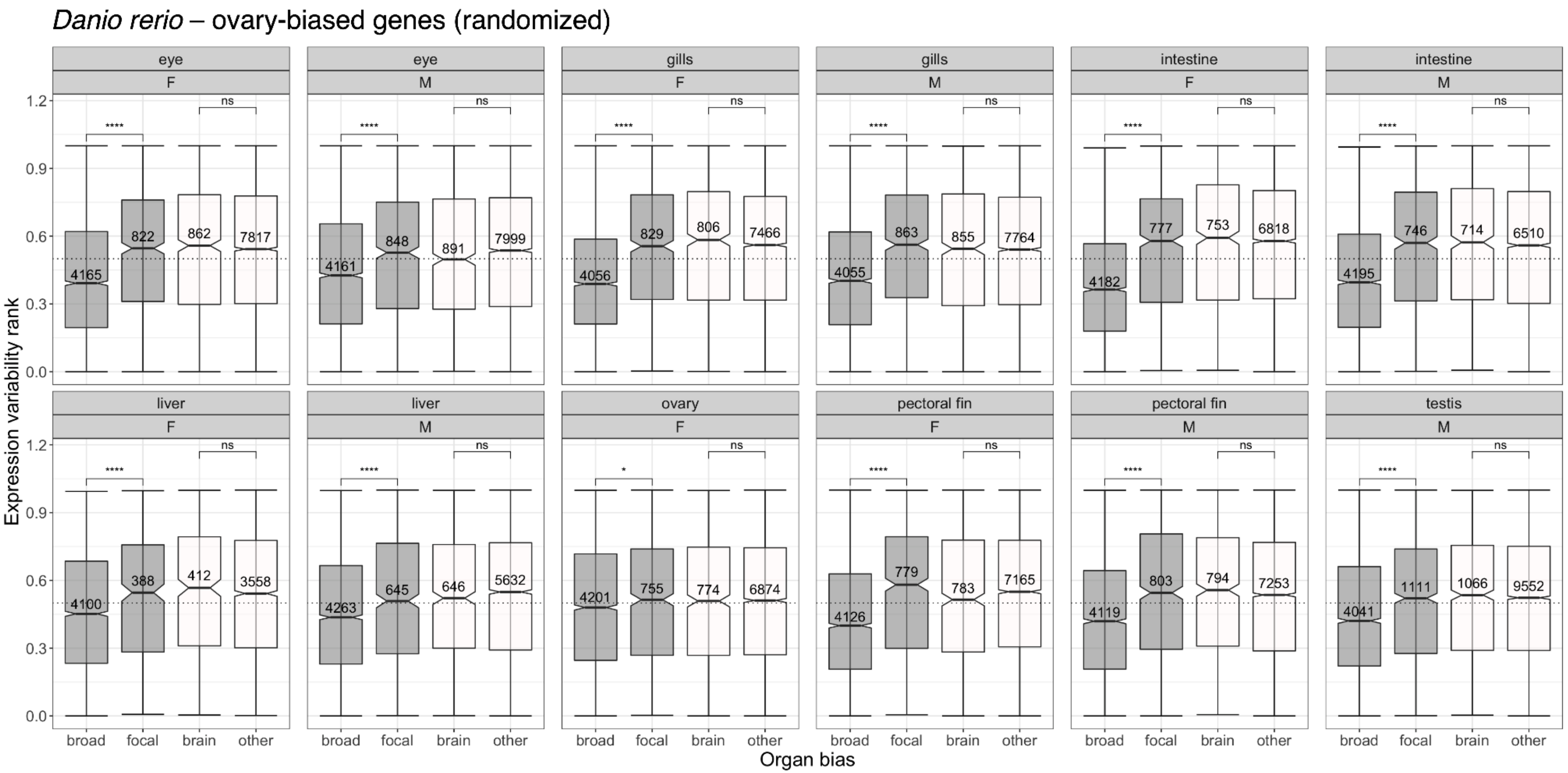


**Figure S48. Distribution of expression variability ranks across organ-sex conditions in zebrafish given randomized organ-biased genes and with an emphasis on 'ovary-biased' genes.** For each gene, the top organ is assigned randomly with seed of 12345 without changing the expression specificity (tau) index. Genes with $\tau \leq 0.3$ are considered broadly expressed ('broad'), whereas genes with $\tau > 0.3$ are classified as organ-biased. Among the set of organ-biased genes, genes are further classified into three categories: focal-biased genes ('focal') are preferentially expressed in the indicated focal organ, ovary-biased genes ('ovary') are preferentially expressed in the ovary, and the remaining ('other') are preferentially expressed in other organs. The number of genes included are annotated for each category. Comparisons between broadly expressed and focal-biased genes (gray boxplots), and between ovary-biased and other organ-biased genes (white boxplots) were performed using Wilcoxon rank-sum tests ($p$-value significance levels: '****': $p \leq 0.0001$, '***': $p \leq 0.001$, '**': $p \leq 0.01$, '*': $p \leq 0.05$, 'ns': $p > 0.05$). The dashed line indicates a variability rank of 0.5.

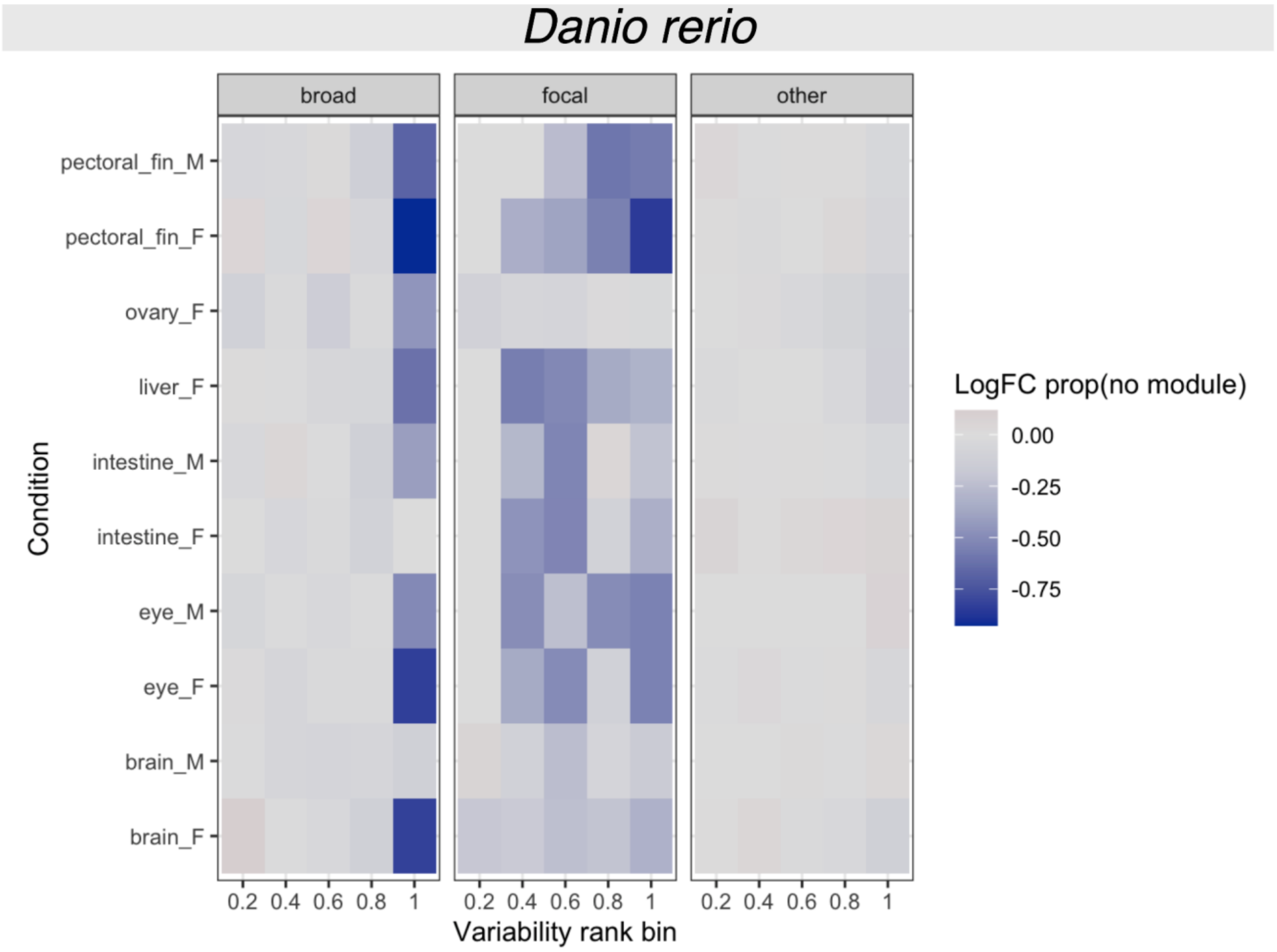


**Figure S49. Depletion of zebrafish genes not assigned to a co-expression module based on weighted gene co-expression network analysis (WGCNA).** Per organ-sex condition with at least 10 biological replicates, genes were grouped into subsets based on organ bias category ("broad", "focal", "other", based on a cutoff of τ ≤ 0.3) and variability rank within the organ in bins of width 0.2. For each gene subset, co-expression modules were inferred using *WGCNA* (Langfelder and Horvath 2008), in which genes without strong co-expression patterns are grouped into a 'dummy' pseudo-module. The proportion of genes assigned to the pseudo-module was compared between observed and randomized data. For each gene subset, a randomized subset was generated by rowwise permutation of the gene x sample matrix, which preserves the mean and variance of each gene while the removing covariance structure between genes. The color gradient shows log-fold enrichment relative to random, such that a logFC < 0 indicates depletion of genes without a proper module assignment.

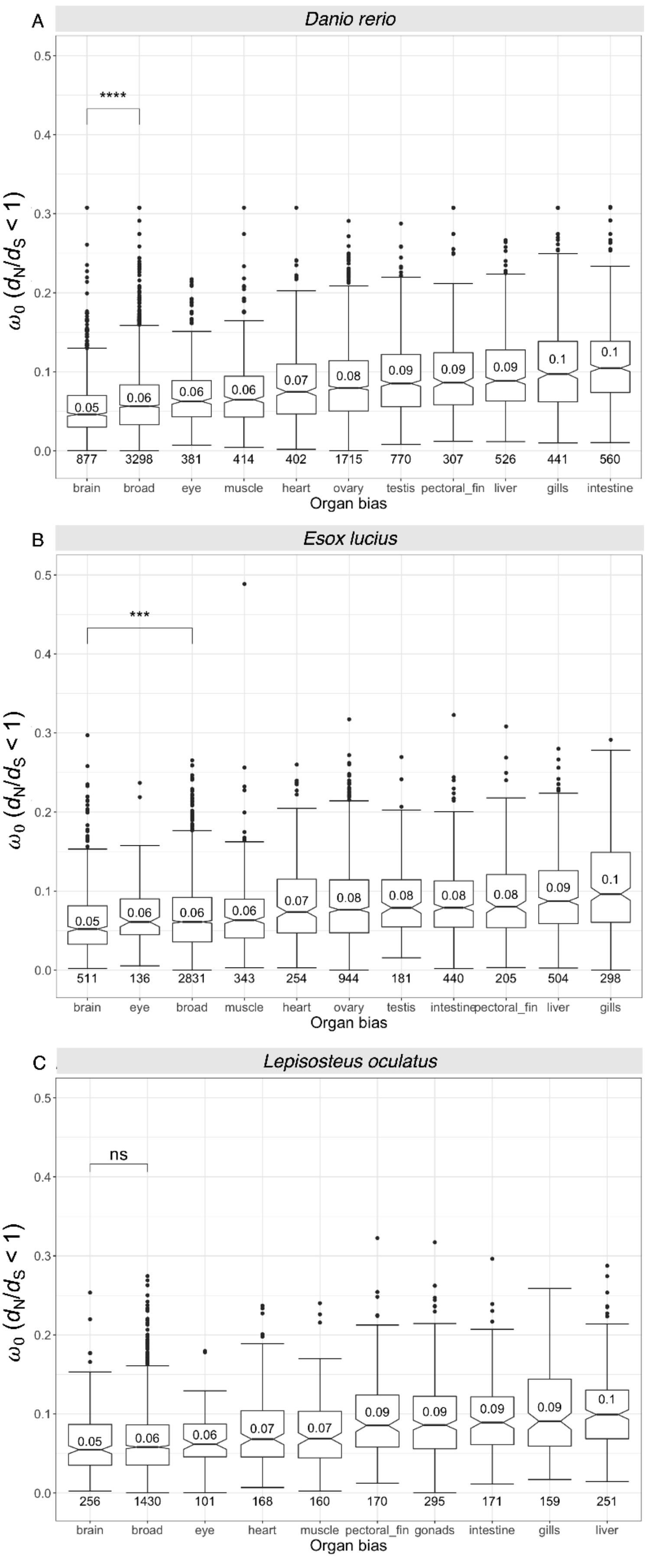
A
Danio rerio
B
Esox lucius
C
Lepisosteus oculatus
ω0 (dN/dS < 1)
Organ bias
brain
broad
eye
muscle
heart
ovary
testis
pectoral_fin
liver
gills
intestine
gonads
****
***
ns

**Figure S50. Strength of purifying selection of genes classified by organ bias, for (A) zebrafish, (B) Northern pike, and (C) spotted gar.** Boxplots show the distribution of ratios of nonsynonymous substitutions ($d_N$) to synonymous substitutions ($d_S$) of sites that have evolved under purifying selection ($\omega_0$ ratio, for $d_N/d_S < 1$) for each organ bias category. Categories are sorted from left to right by increasing median $\omega_0$. Broadly expressed genes ('broad') have a measured organ expression specificity index (tau) of $\tau \leq 0.3$, while genes with $\tau > 0.3$ are considered organ-biased and classified based on their top organ of expression. The median $\omega_0$ is indicated within each boxplot and the number of genes included are annotated below each. Tested branches included in all three species (A,B,C) are: Euteleostomi, Actinopterygii, Neopterygii; for teleosts only (A,B): Osteoglossocephalai, Clupeocephala; for zebrafish only (A): Otomorpha, Otophysi; and for Northern pike only (B): Euteleosteomorpha, Protacanthopterygii **(Figure 4A)**. Pairwise comparisons between brain-biased and broadly expressed genes were performed using Wilcoxon rank-sum tests (*p*-value significance levels: '****': $p \leq 0.0001$, '***': $p \leq 0.001$, '**': $p \leq 0.01$, '*': $p \leq 0.05$, 'ns': $p > 0.05$).

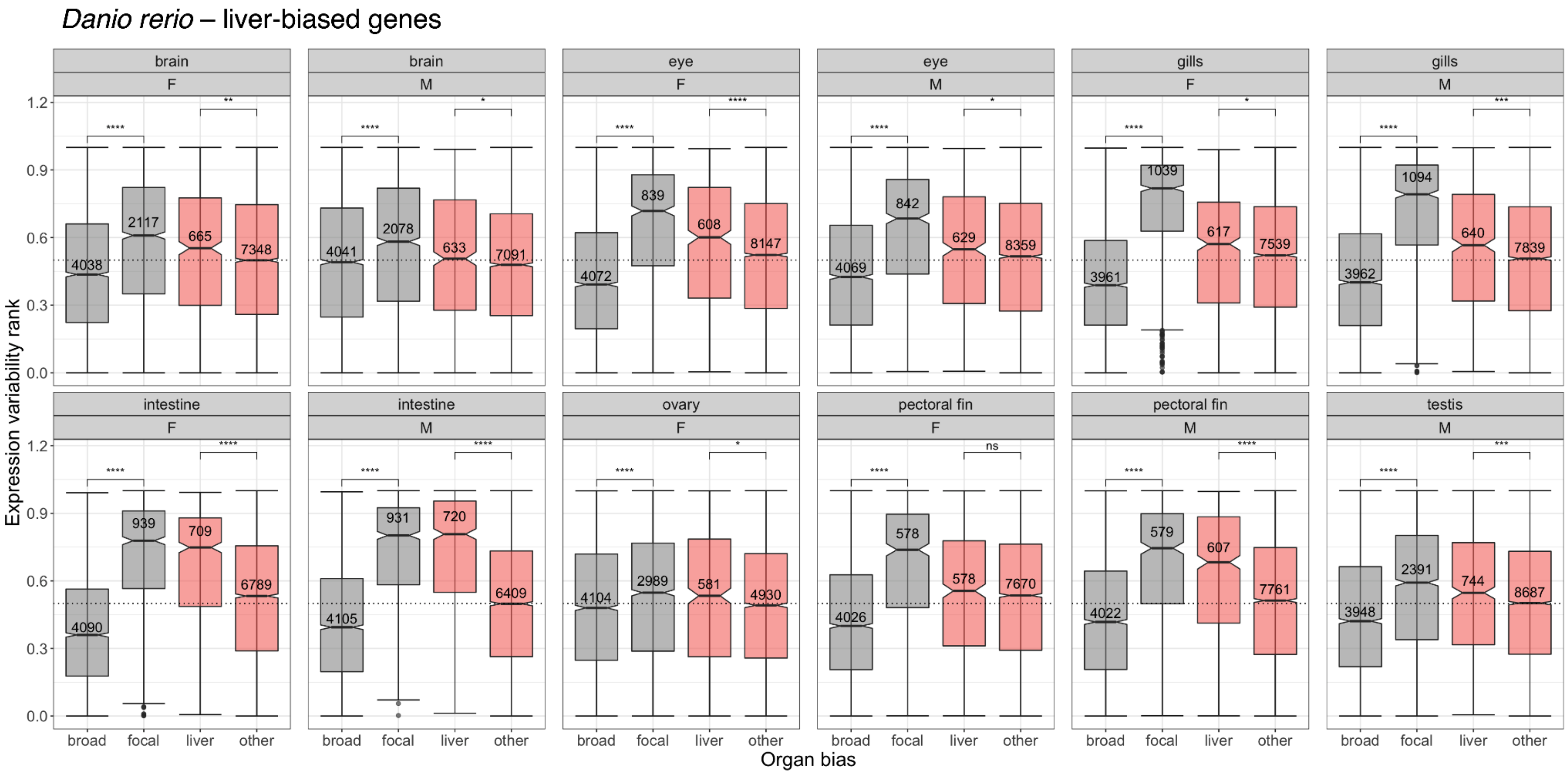


**Figure S51. Distribution of expression variability ranks across organ-sex conditions in zebrafish, for genes classified by organ bias and with an emphasis on liver-biased genes.** Broadly expressed genes ('broad') have a measured organ expression specificity index (tau) of $\tau \leq 0.3$; genes with $\tau > 0.3$ are considered organ-biased. Among the set of organ-biased genes, genes are further classified into three categories: focal-biased genes ('focal') are preferentially expressed in the indicated focal organ, liver-biased genes ('liver') are preferentially expressed in the liver, and the remaining ('other') are preferentially expressed in other organs. The number of genes included are annotated for each category. Comparisons between broadly expressed and focal-biased genes (gray boxplots), and between liver-biased and other organ-biased genes (salmon boxplots) were performed using Wilcoxon rank-sum tests ($p$-value significance levels: '****': $p \leq 0.0001$, '***': $p \leq 0.001$, '**': $p \leq 0.01$, '*': $p \leq 0.05$, 'ns': $p > 0.05$). The dashed line indicates a variability rank of 0.5.

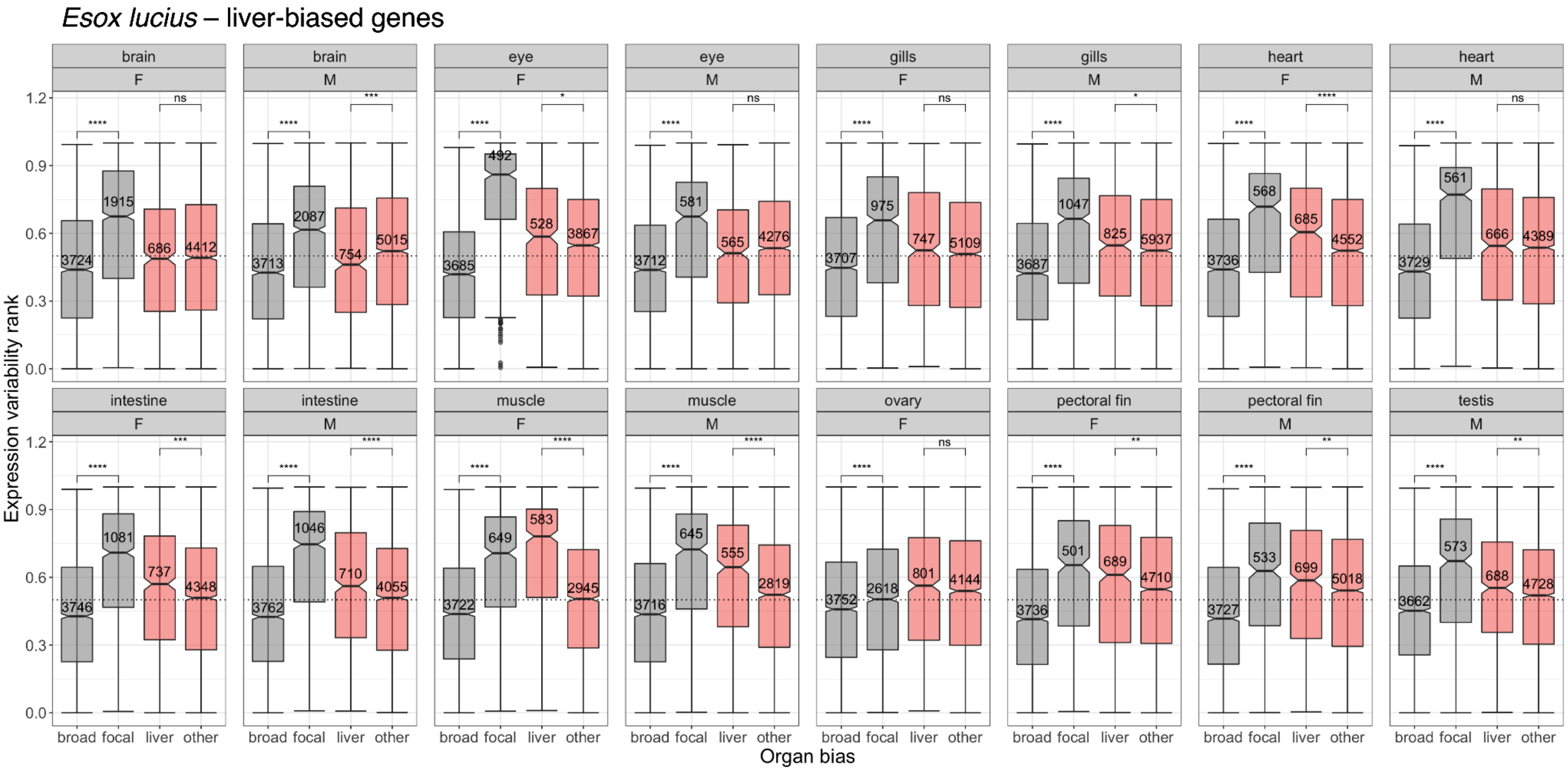


**Figure S52. Distribution of expression variability ranks across organ-sex conditions in Northern pike, for genes classified by organ bias and with an emphasis on liver-biased genes.** Broadly expressed genes ('broad') have a measured organ expression specificity index (tau) of $\tau \leq 0.3$; genes with $\tau > 0.3$ are considered organ-biased. Among the set of organ-biased genes, genes are further classified into three categories: focal-biased genes ('focal') are preferentially expressed in the indicated focal organ, liver-biased genes ('liver') are preferentially expressed in the liver, and the remaining ('other') are preferentially expressed in other organs. The number of genes included are annotated for each category. Comparisons between broadly expressed and focal-biased genes (gray boxplots), and between liver-biased and other organ-biased genes (salmon boxplots) were performed using Wilcoxon rank-sum tests ($p$-value significance levels: '****': $p \leq 0.0001$, '***': $p \leq 0.001$, '**': $p \leq 0.01$, '*': $p \leq 0.05$, 'ns': $p > 0.05$). The dashed line indicates a variability rank of 0.5.

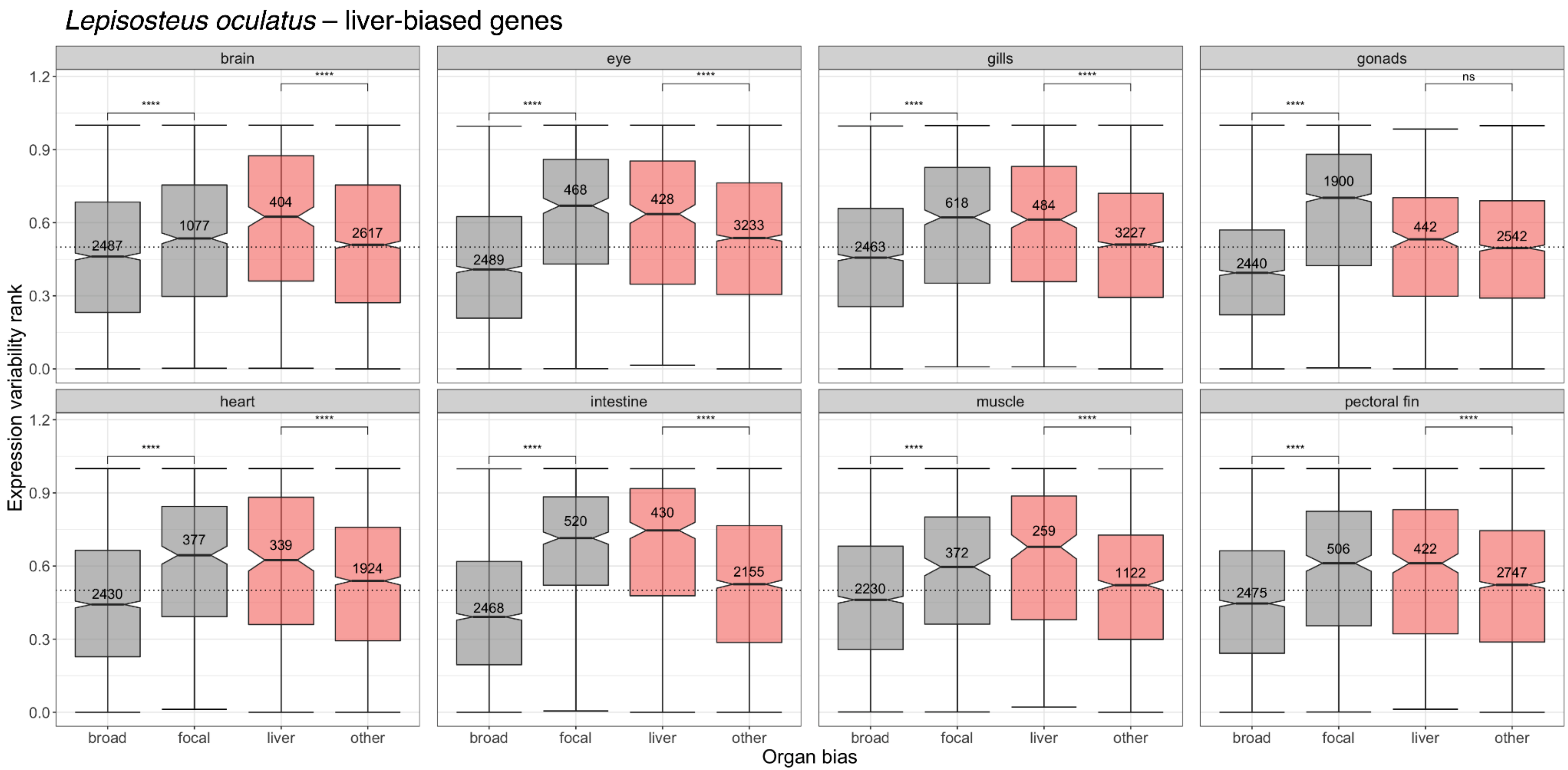


**Figure S53. Distribution of expression variability ranks across organs in spotted gar, for genes classified by organ bias and with an emphasis on liver-biased genes.** Broadly expressed genes ('broad') have a measured organ expression specificity index (tau) of $\tau \leq 0.3$; genes with $\tau > 0.3$ are considered organ-biased. Among the set of organ-biased genes, genes are further classified into three categories: focal-biased genes ('focal') are preferentially expressed in the indicated focal organ, liver-biased genes ('liver') are preferentially expressed in the liver, and the remaining ('other') are preferentially expressed in other organs. The number of genes included are annotated for each category. Comparisons between broadly expressed and focal-biased genes (gray boxplots), and between liver-biased and other organ-biased genes (salmon boxplots) were performed using Wilcoxon rank-sum tests ($p$-value significance levels: '****': $p \leq 0.0001$, '***': $p \leq 0.001$, '**': $p \leq 0.01$, '*': $p \leq 0.05$, 'ns': $p > 0.05$). The dashed line indicates a variability rank of 0.5.

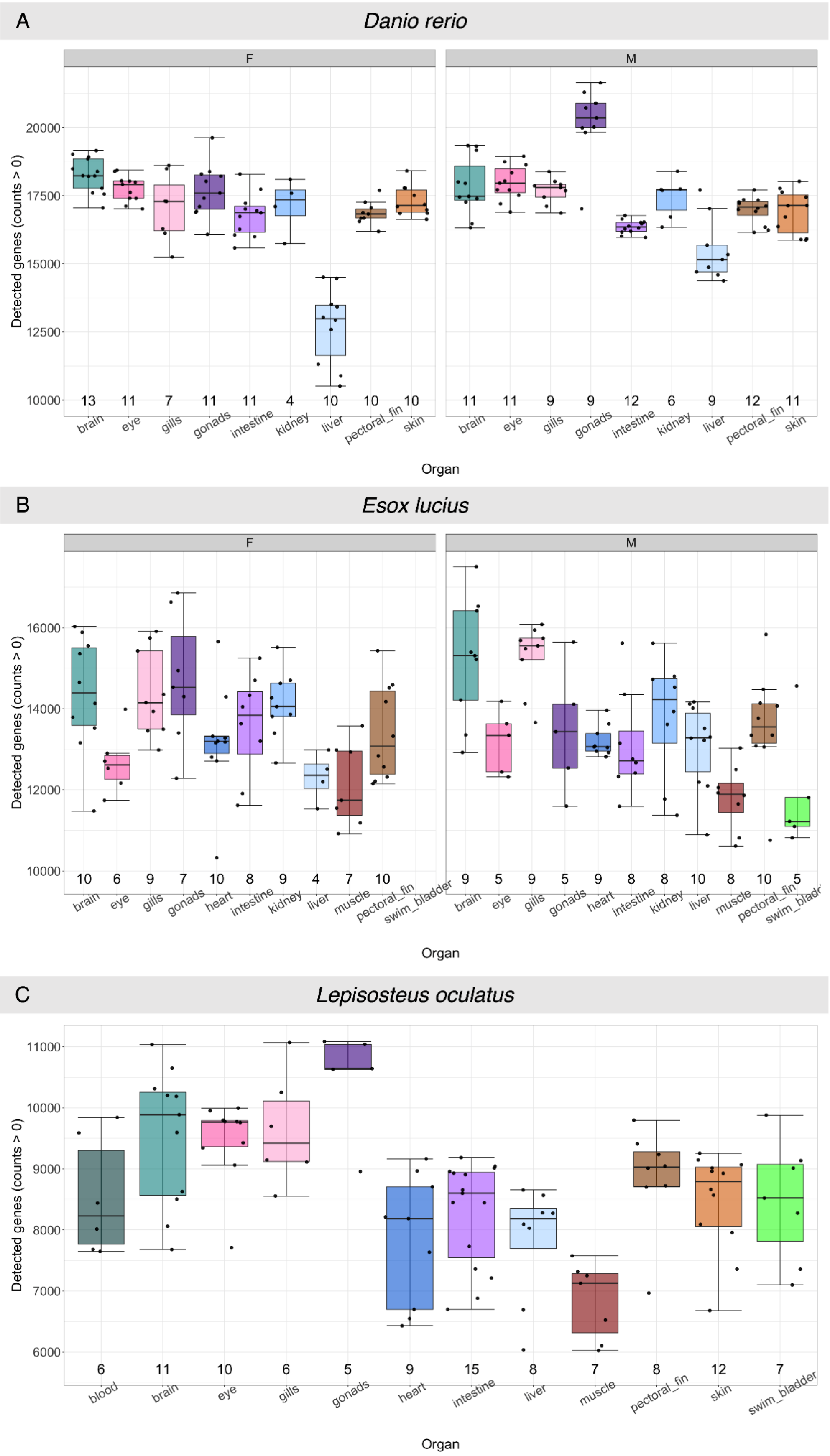


**Figure S54. Boxplots of detected genes per organ for (A) zebrafish, (B) Northern pike, and (C) spotted gar.** For zebrafish and Northern pike, boxplots for female ('F') and male ('M') samples are plotted separately. Per sample, a gene is 'detected' if the read count is greater than 0. The number of individuals per organ that have been retained after quality filtering are indicated below each boxplot. Only organs with at least 4 individuals are included.

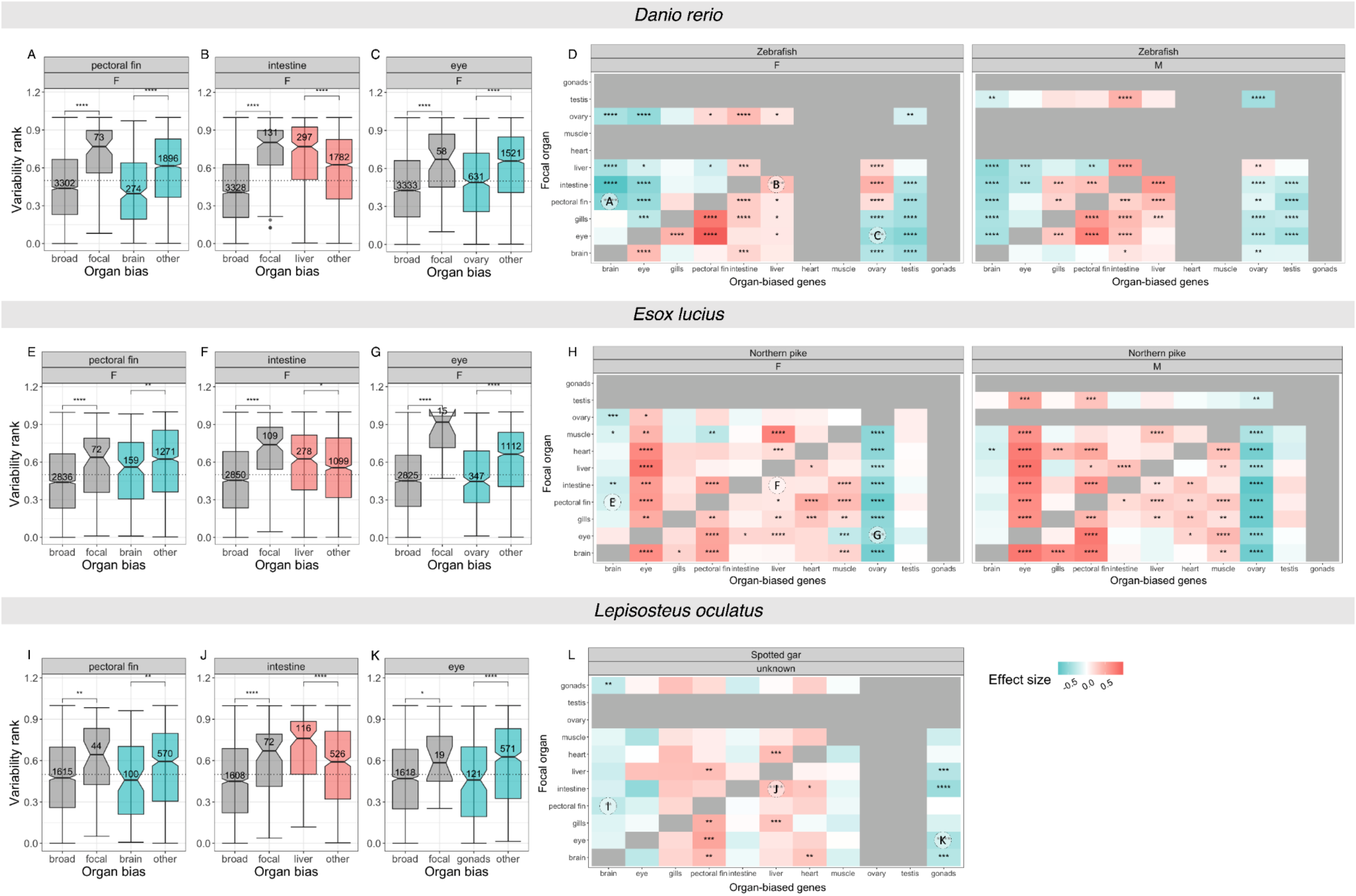
Danio rerio
A
B
C
D
pectoral fin
intestine
eye
F
M
Zebrafish
Variability rank
broad
focal
brain
liver
ovary
other
Organ bias
Focal organ
Organ-biased genes
gonads
testis
ovary
muscle
heart
liver
intestine
pectoral fin
gills
eye
brain
Esox lucius
E
F
G
H
Northern pike
Lepisosteus oculatus
I
J
K
L
Spotted gar
unknown
gonads
Effect size
-0.5
0.0
0.5

**Figure S55. Relationship between expression variability and organ-biased expression for (A-D) zebrafish, (E-H) Northern pike, and (I-L) spotted gar for genes with nonzero counts across all samples within-species.** Each component of the figure is as described in **Figure 5**, except that the set of genes included in the analysis is reduced. Per organ- or organ-sex condition within each species, expression variability ranks were computed considering only genes that have nonzero counts across all samples. **(A)** Boxplot of expression variability ranks of genes expressed in female ('F') zebrafish pectoral fin, for sets of genes classified by organ bias and with an emphasis on brain-biased genes. Among the set of organ-biased genes, genes are further classified into three categories: focal-biased genes ('focal') are preferentially expressed in the focal organ (i.e., pectoral fin), brain-biased genes ('brain') are preferentially expressed in the brain, and the remaining ('other') are preferentially expressed in other organs. Comparisons between broadly expressed and focal-biased genes, and between brain-biased and other organ-biased genes were performed using Wilcoxon rank-sum tests (*p*-value significance levels: '****': $p \leq 0.0001$, '***': $p \leq 0.001$, '**': $p \leq 0.01$, '*': $p \leq 0.05$, 'ns': $p > 0.05$). The dashed line indicates a variability rank of 0.5. **(B)** Boxplot of variability ranks of genes expressed in female zebrafish intestine, with an emphasis on liver-biased genes. **(C)** Boxplot of variability ranks of genes expressed in female zebrafish eye, with an emphasis on ovary-biased genes. **(D)** Heatmap summarizing pairwise comparisons of expression variability ranks between subsets of organ-biased genes for female ('F', left) and male ('M', right) zebrafish. Rows correspond to focal organ whereas columns correspond to subsets of organ-biased genes. Pairwise comparisons were performed using Wilcoxon rank-sum tests with Benjamini-Hochberg ('BH') correction for multiple testing (adjusted *p*-value < 0.05). Effect size was computed using Glass's delta ($\delta$), with color scale shown in (L). Cells in gray are not included in the dataset. Three cells are annotated, corresponding to the following comparisons in boxplots (A-C). A negative effect size entails lower expression variability relative to other organ-biased genes (A,C) (turquoise), whereas a positive effect size entails higher variability (B) (salmon). **(E-H)** Corresponding plots for Northern pike, with cells annotated corresponding to comparisons in boxplots (E-G). **(I-L)** Corresponding plots for unsexed spotted gar, with cells annotated corresponding to comparisons in boxplots (I-K).

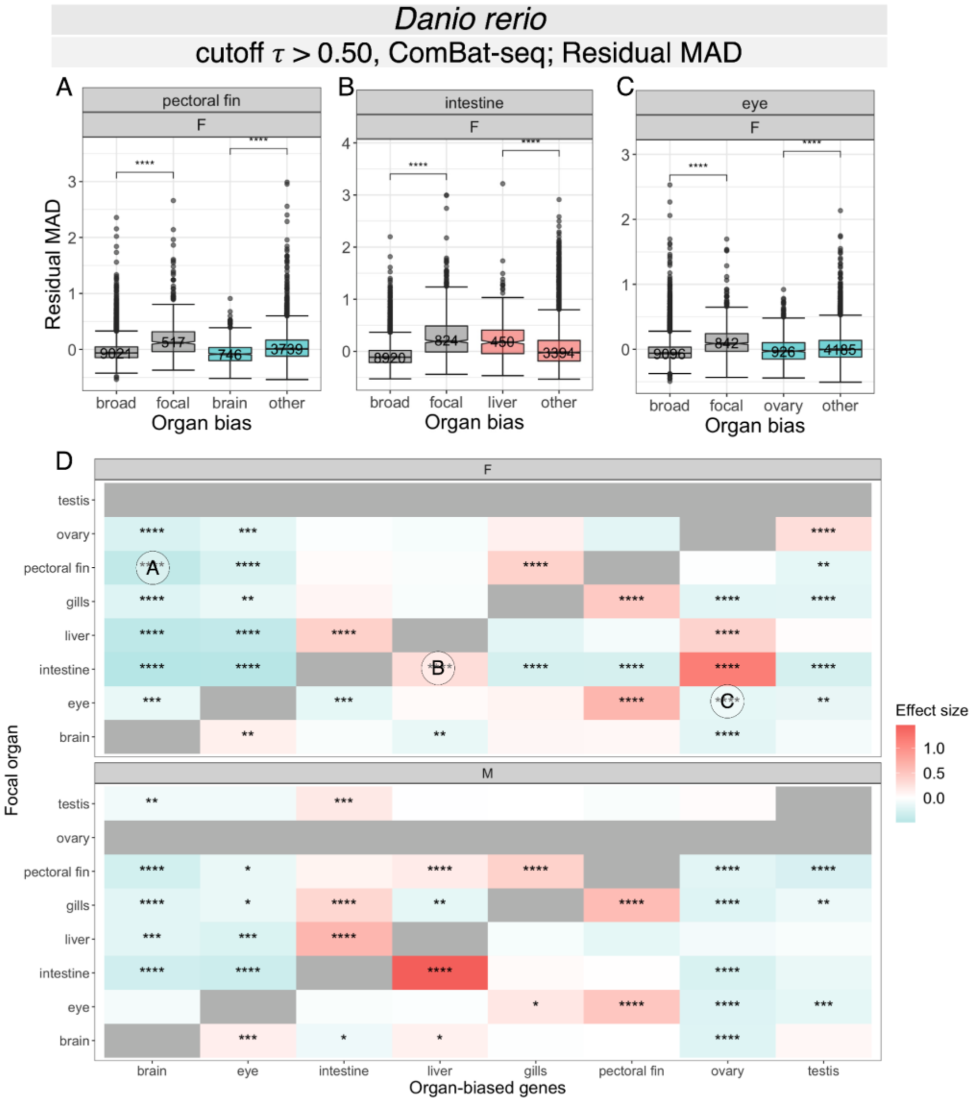


**Figure S56. Relationship between expression variability and organ-biased expression for zebrafish.** Batch correction with *ComBat-seq* (Zhang et al. 2020) was performed prior to gene filtering, normalization, expression specificity and variability analysis. Expression variability per gene within a condition was estimated using residual median absolute deviation (MAD) without jackknife resampling. **(A)** Boxplot of expression variability ranks of genes expressed in female ('F') zebrafish pectoral fin, for sets of genes classified by organ bias and with an emphasis on brain-biased genes. Broadly expressed genes ('broad') have a measured organ expression specificity index (tau) of $\tau \leq 0.5$; genes with $\tau > 0.5$ are considered organ-biased. Among the set of organ-biased genes, genes are further classified into three categories: focal-biased genes ('focal') are preferentially expressed in the focal organ (i.e., pectoral fin), brain-biased genes ('brain') are preferentially expressed in the brain, and the remaining ('other') are preferentially expressed in other organs. The number of genes included are annotated for each category. Comparisons between broadly expressed and focal-biased genes, and between brain-biased and other organ-biased genes were performed using Wilcoxon rank-sum tests ($p$-value significance levels: '****': $p \leq 0.0001$, '***': $p \leq 0.001$, '**': $p \leq 0.01$, '*': $p \leq 0.05$, 'ns': $p > 0.05$). The dashed line indicates a variability rank of 0.5. **(B)** Boxplot of variability ranks of genes expressed in female zebrafish intestine, with genes classified by organ bias and with an emphasis on liver-biased genes. **(C)** Boxplot of variability ranks of genes expressed in female zebrafish eye, with genes classified by organ bias and with an emphasis on ovary-biased genes. **(D)** Heatmap summarizing pairwise comparisons of expression variability ranks between subsets of organ-biased genes for female ('F', top) and male ('M', bottom) zebrafish. Rows correspond to focal organ whereas columns correspond to subsets of organ-biased genes. Pairwise comparisons were performed using Wilcoxon rank-sum tests with Benjamini-Hochberg ('BH') correction for multiple testing (adjusted $p$-value < 0.05). Effect size was computed using Glass's delta ($\delta$), with color scale shown in (L). Cells in gray are not included in the dataset. Three cells are annotated, corresponding to the following comparisons in boxplots (A-C): (A) brain-biased vs. other organ-biased genes expressed in female zebrafish pectoral fin, (B) liver-biased vs. other organ-biased biased genes expressed in female zebrafish intestine, and (C) ovary-biased vs. other organ-biased genes expressed in female zebrafish eye. A negative effect size entails lower expression variability relative to other organ-biased genes (A,C) (turquoise), whereas a positive effect size entails higher variability (B) (salmon).

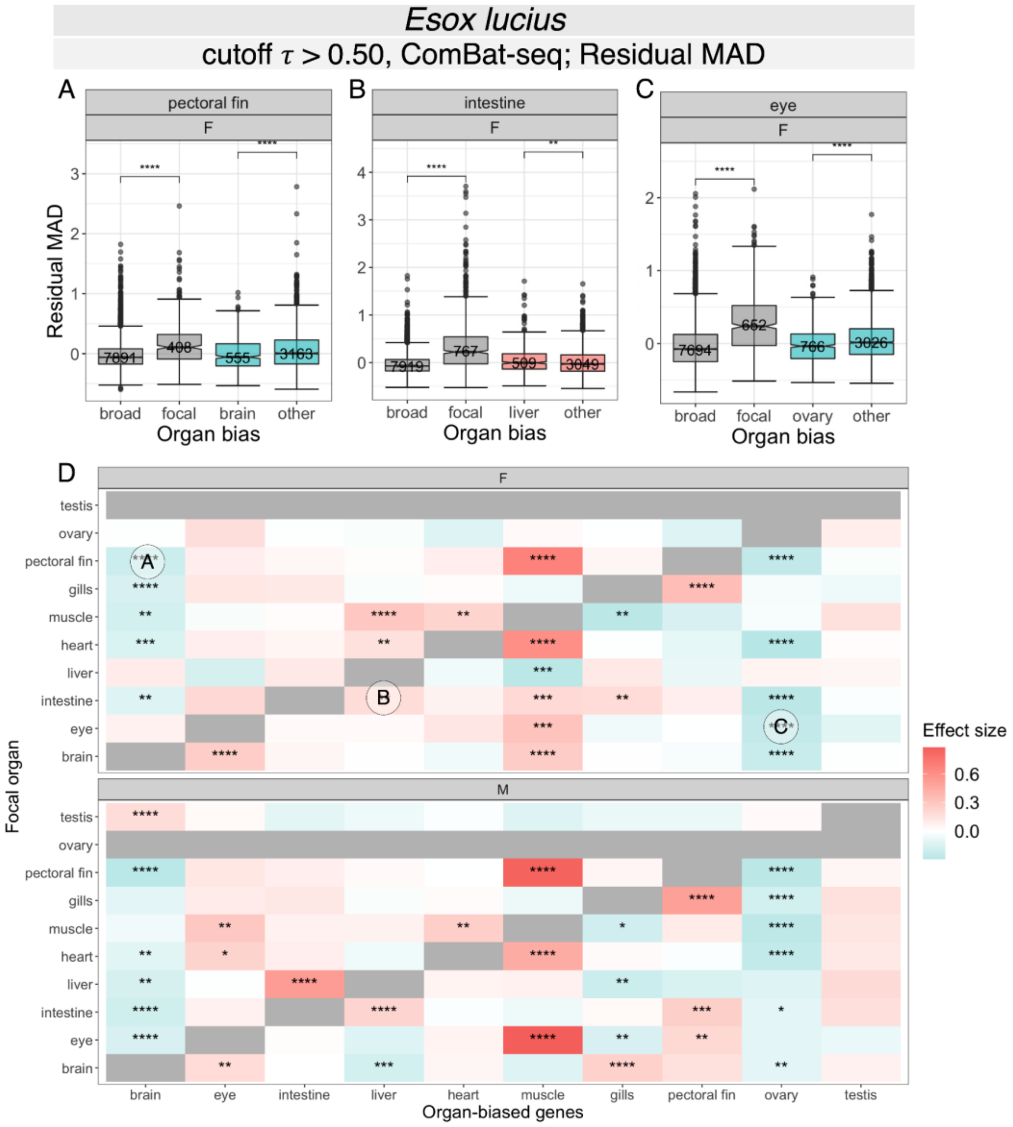


**Figure S57. Relationship between expression variability and organ-biased expression for Northern pike.** Batch correction with *ComBat-seq* (Zhang et al. 2020) was performed prior to gene filtering, normalization, expression specificity and variability analysis. Expression variability per gene within a condition was estimated using residual median absolute deviation (MAD) without jackknife resampling. **(A)** Boxplot of expression variability ranks of genes expressed in female ('F') Northern pike pectoral fin, for sets of genes classified by organ bias and with an emphasis on brain-biased genes. Broadly expressed genes ('broad') have a measured organ expression specificity index (tau) of $\tau \leq 0.5$; genes with $\tau > 0.5$ are considered organ-biased. Among the set of organ-biased genes, genes are further classified into three categories: focal-biased genes ('focal') are preferentially expressed in the focal organ (i.e., pectoral fin), brain-biased genes ('brain') are preferentially expressed in the brain, and the remaining ('other') are preferentially expressed in other organs. The number of genes included are annotated for each category. Comparisons between broadly expressed and focal-biased genes, and between brain-biased and other organ-biased genes were performed using Wilcoxon rank-sum tests (*p*-value significance levels: '****': $p \leq 0.0001$, '***': $p \leq 0.001$, '**': $p \leq 0.01$, '*': $p \leq 0.05$, 'ns': $p > 0.05$). The dashed line indicates a variability rank of 0.5. **(B)** Boxplot of variability ranks of genes expressed in female Northern pike intestine, with genes classified by organ bias and with an emphasis on liver-biased genes. **(C)** Boxplot of variability ranks of genes expressed in female Northern pike eye, with genes classified by organ bias and with an emphasis on ovary-biased genes. **(D)** Heatmap summarizing pairwise comparisons of expression variability ranks between subsets of organ-biased genes for female ('F', top) and male ('M', bottom) individuals. Rows correspond to focal organ whereas columns correspond to subsets of organ-biased genes**.** Pairwise comparisons were performed using Wilcoxon rank-sum tests with Benjamini-Hochberg ('BH') correction for multiple testing (adjusted *p*-value < 0.05). Effect size was computed using Glass's delta (δ), with color scale shown in (L). Cells in gray are not included in the dataset. Three cells are annotated, corresponding to the following comparisons in boxplots (A-C): (A) brain-biased vs. other organ-biased genes expressed in female Northern pike pectoral fin, (B) liver-biased vs. other organ-biased biased genes expressed in female intestine, and (C) ovary-biased vs. other organ-biased genes expressed in female eye. A negative effect size entails lower expression variability relative to other organ-biased genes (A,C) (turquoise), whereas a positive effect size entails higher variability (B) (salmon).

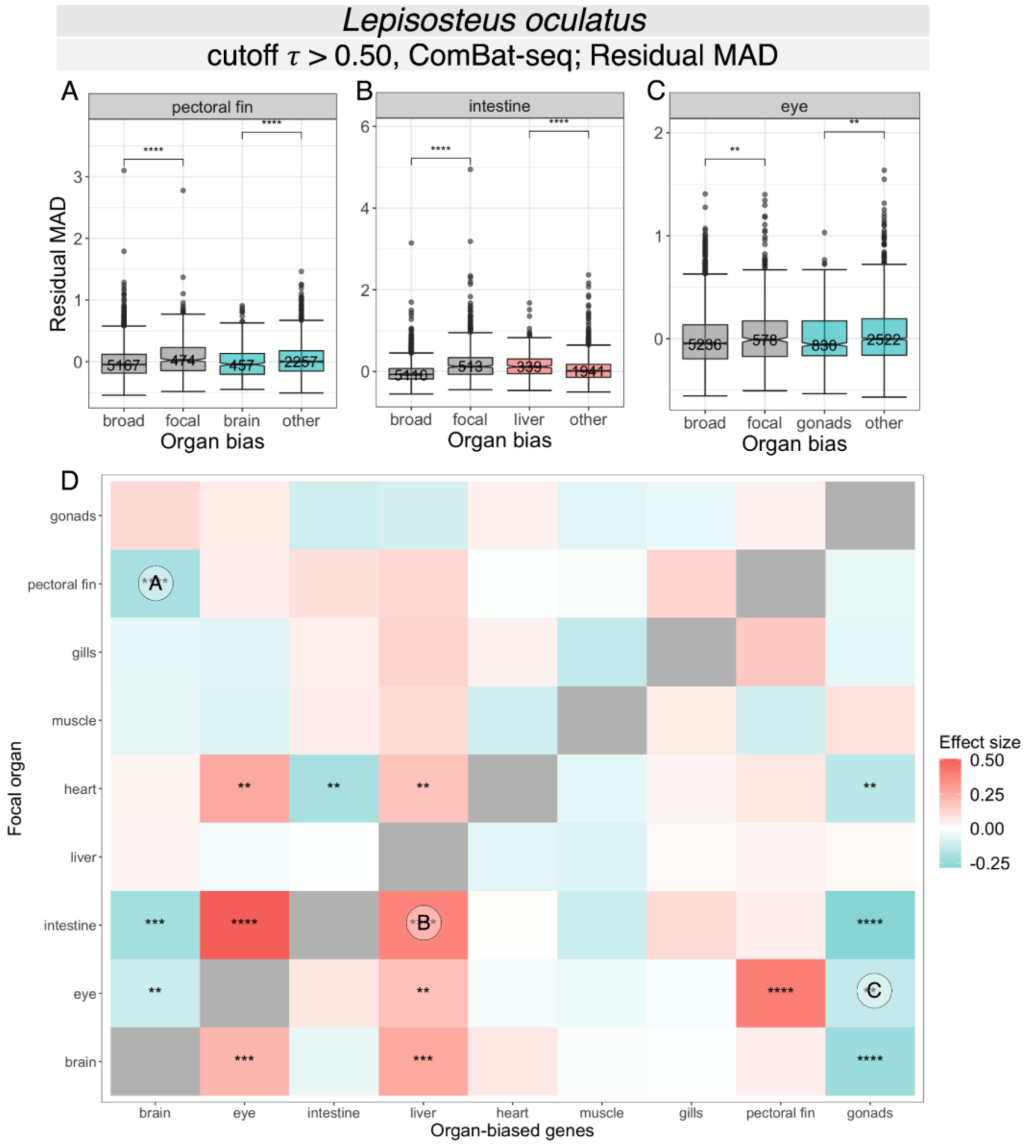


**Figure S58. Relationship between expression variability and organ-biased expression for spotted gar.** Batch correction with *ComBat-seq* (Zhang et al. 2020) was performed prior to gene filtering, normalization, expression specificity and variability analysis. Expression variability per gene within a condition was estimated using residual median absolute deviation (MAD) without jackknife resampling. **(A)** Boxplot of expression variability ranks of genes expressed in spotted gar pectoral fin, for sets of genes classified by organ bias and with an emphasis on brain-biased genes. Broadly expressed genes ('broad') have a measured organ expression specificity index (tau) of $\tau \leq 0.5$; genes with $\tau > 0.5$ are considered organ-biased. Among the set of organ-biased genes, genes are further classified into three categories: focal-biased genes ('focal') are preferentially expressed in the focal organ (i.e., pectoral fin), brain-biased genes ('brain') are preferentially expressed in the brain, and the remaining ('other') are preferentially expressed in other organs. The number of genes included are annotated for each category. Comparisons between broadly expressed and focal-biased genes, and between brain-biased and other organ-biased genes were performed using Wilcoxon rank-sum tests (*p*-value significance levels: '****': $p \leq 0.0001$, '***': $p \leq 0.001$, '**': $p \leq 0.01$, '*': $p \leq 0.05$, 'ns': $p > 0.05$). The dashed line indicates a variability rank of 0.5. **(B)** Boxplot of variability ranks of genes expressed in spotted gar intestine, with genes classified by organ bias and with an emphasis on liver-biased genes. **(C)** Boxplot of variability ranks of genes expressed in spotted gar eye, with genes classified by organ bias and with an emphasis on gonads-biased genes. **(D)** Heatmap summarizing pairwise comparisons of expression variability ranks between subsets of organ-biased genes. Rows correspond to focal organ whereas columns correspond to subsets of organ-biased genes**.** Pairwise comparisons were performed using Wilcoxon rank-sum tests with Benjamini-Hochberg ('BH') correction for multiple testing (adjusted *p*-value < 0.05). Effect size was computed using Glass's delta (δ), with color scale shown in (L). Cells in gray are not included in the dataset. Three cells are annotated, corresponding to the following comparisons in boxplots (A-C): (A) brain-biased vs. other organ-biased genes expressed in the pectoral fin, (B) liver-biased vs. other organ-biased biased genes expressed in the intestine, and (C) ovary-biased vs. other organ-biased genes expressed in the eye. A negative effect size entails lower expression variability relative to other organ-biased genes (A,C) (turquoise), whereas a positive effect size entails higher variability (B) (salmon).

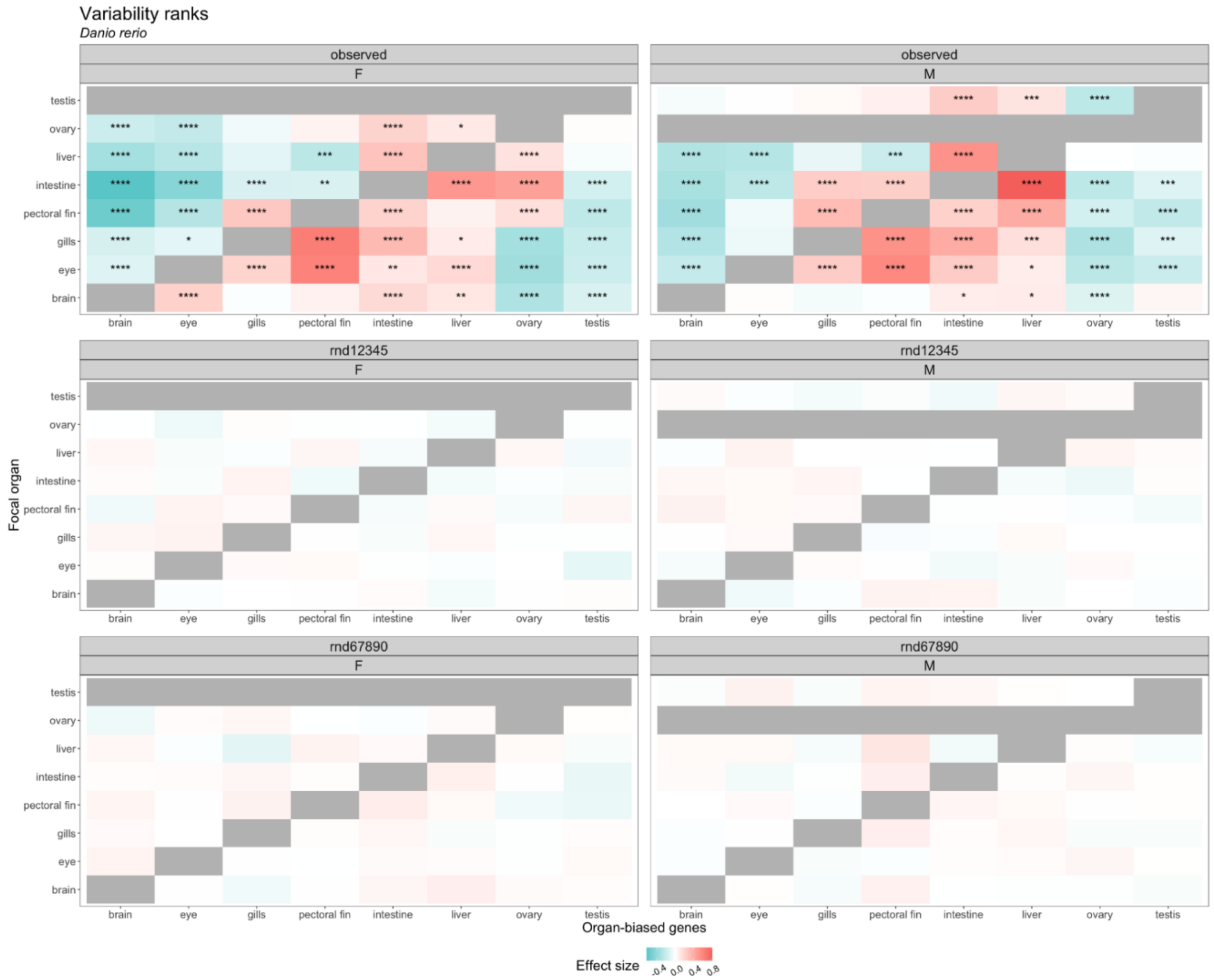


**Figure S59. Condition-dependent expression variability of organ-biased genes in zebrafish, given observed and randomized data.** Heatmaps summarizing multiple pairwise comparisons of expression variability ranks between subsets of organ-biased genes for female ('F', left column) and male ('M', right column) zebrafish, as previously described in **Figure 5**. Rows correspond to focal organ whereas columns correspond to subsets of organ-biased genes. Pairwise comparisons were performed using Wilcoxon rank-sum tests with Benjamini-Hochberg ('BH') correction for multiple testing (adjusted $p$-value < 0.05 with $p$-value significance levels: '****': $p \leq 0.0001$, '***': $p \leq 0.001$, '**': $p \leq 0.01$, '*': $p \leq 0.05$). Effect size was computed using Glass's delta (δ). Cells in gray are not included in the dataset. Top row: Results from observed data, as in **Figure 5D**. Middle row: Randomized data. The top organ of each organ-biased gene is assigned randomly with seed of 12345. **A-D**. Bottom row: Randomized data with seed of 67890.

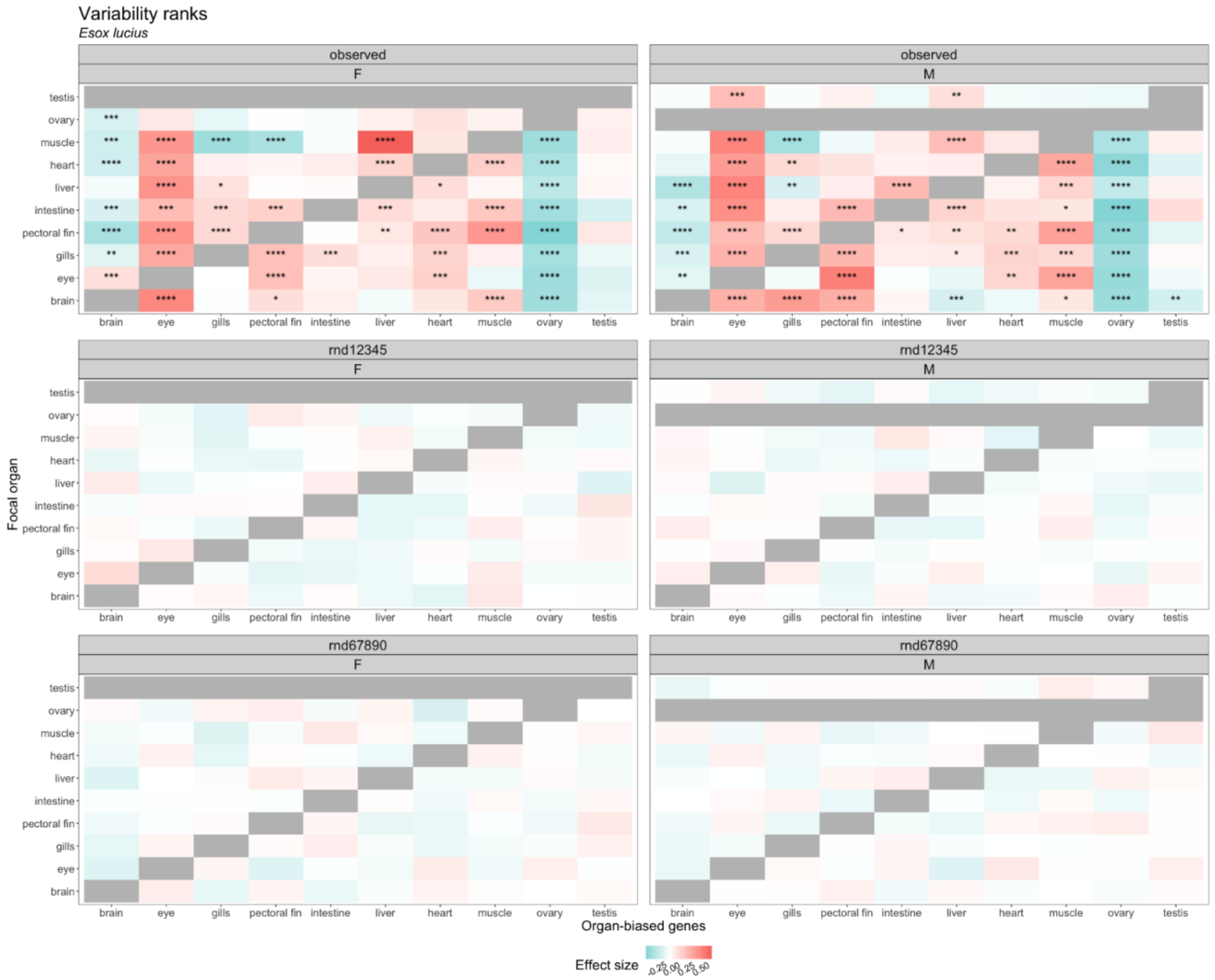


**Figure S60. Condition-dependent expression variability of organ-biased genes in Northern pike, given observed and randomized data.** Heatmaps summarizing multiple pairwise comparisons of expression variability ranks between subsets of organ-biased genes for female ('F', left column) and male ('M', right column) Northern pike, as previously described in **Figure 5**. Rows correspond to focal organ whereas columns correspond to subsets of organ-biased genes. Pairwise comparisons were performed using Wilcoxon rank-sum tests with Benjamini-Hochberg ('BH') correction for multiple testing (adjusted *p*-value < 0.05 with *p*-value significance levels: '****': $p \leq 0.0001$, '***': $p \leq 0.001$, '**': $p \leq 0.01$, '*': $p \leq 0.05$). Effect size was computed using Glass's delta (δ). Cells in gray are not included in the dataset. Top row: Results from observed data, as in **Figure 5H**. Middle row: Randomized data. The top organ of each organ-biased gene is assigned randomly. Bottom row: Randomized data with seed of 67890.

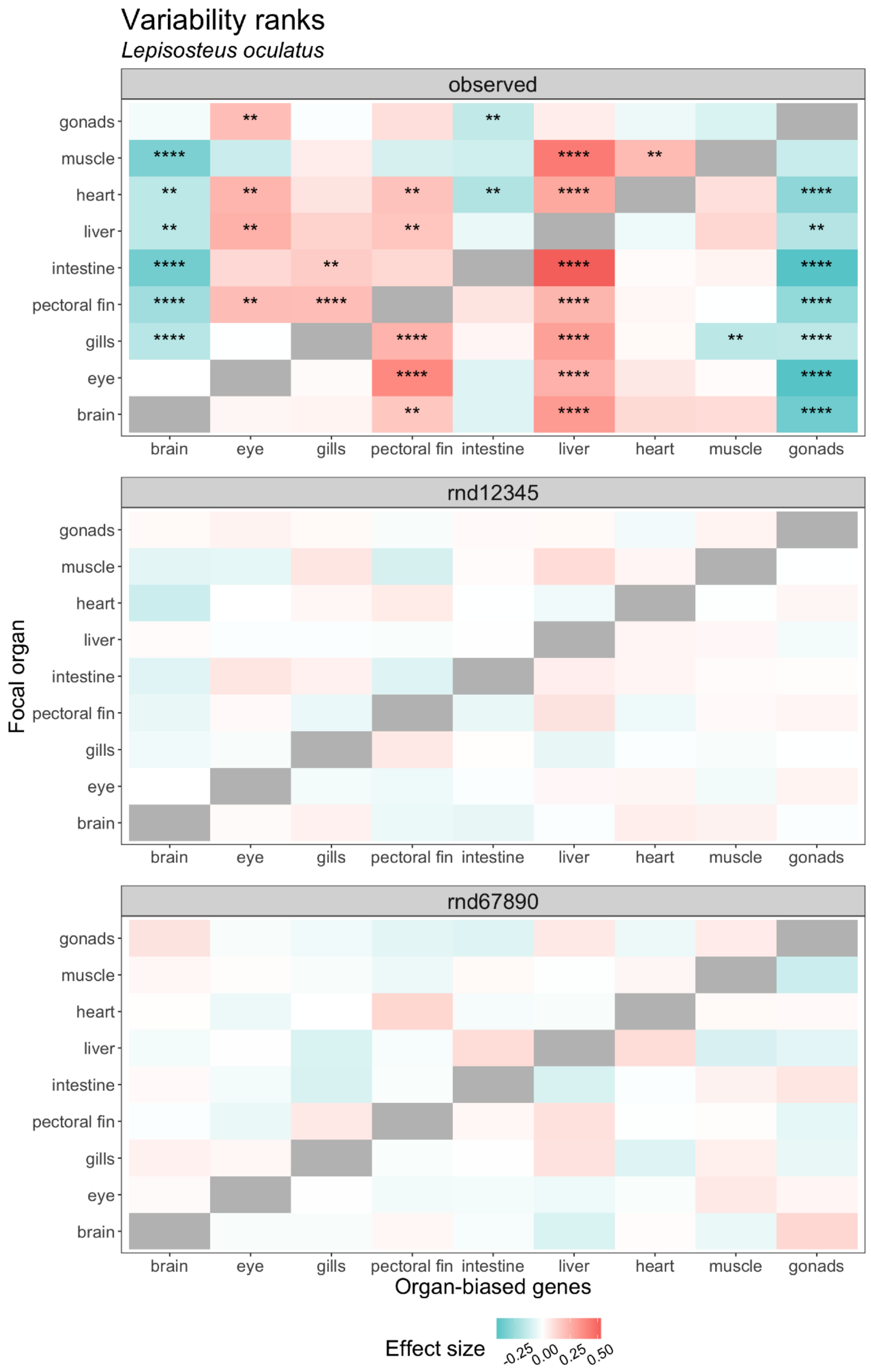


**Figure S61. Condition-dependent expression variability of organ-biased genes in spotted gar, given observed and randomized data.** Heatmaps summarizing multiple pairwise comparisons of expression variability ranks between subsets of organ-biased genes for unsexed spotted gar, as previously described in **Figure 5**. Rows correspond to focal organ whereas columns correspond to subsets of organ-biased genes. Pairwise comparisons were performed using Wilcoxon rank-sum tests with Benjamini-Hochberg ('BH') correction for multiple testing (adjusted *p*-value < 0.05 with *p*-value significance levels: '****': $p \leq 0.0001$, '***': $p \leq 0.001$, '**': $p \leq 0.01$, '*': $p \leq 0.05$). Effect size was computed using Glass's delta (δ). Cells in gray are not included in the dataset. Top row: Results from observed data, as in **Figure 5L**. Middle row: Randomized data. The top organ of each organ-biased gene is assigned randomly with seed of 12345. Bottom row: Randomized data with seed of 67890.

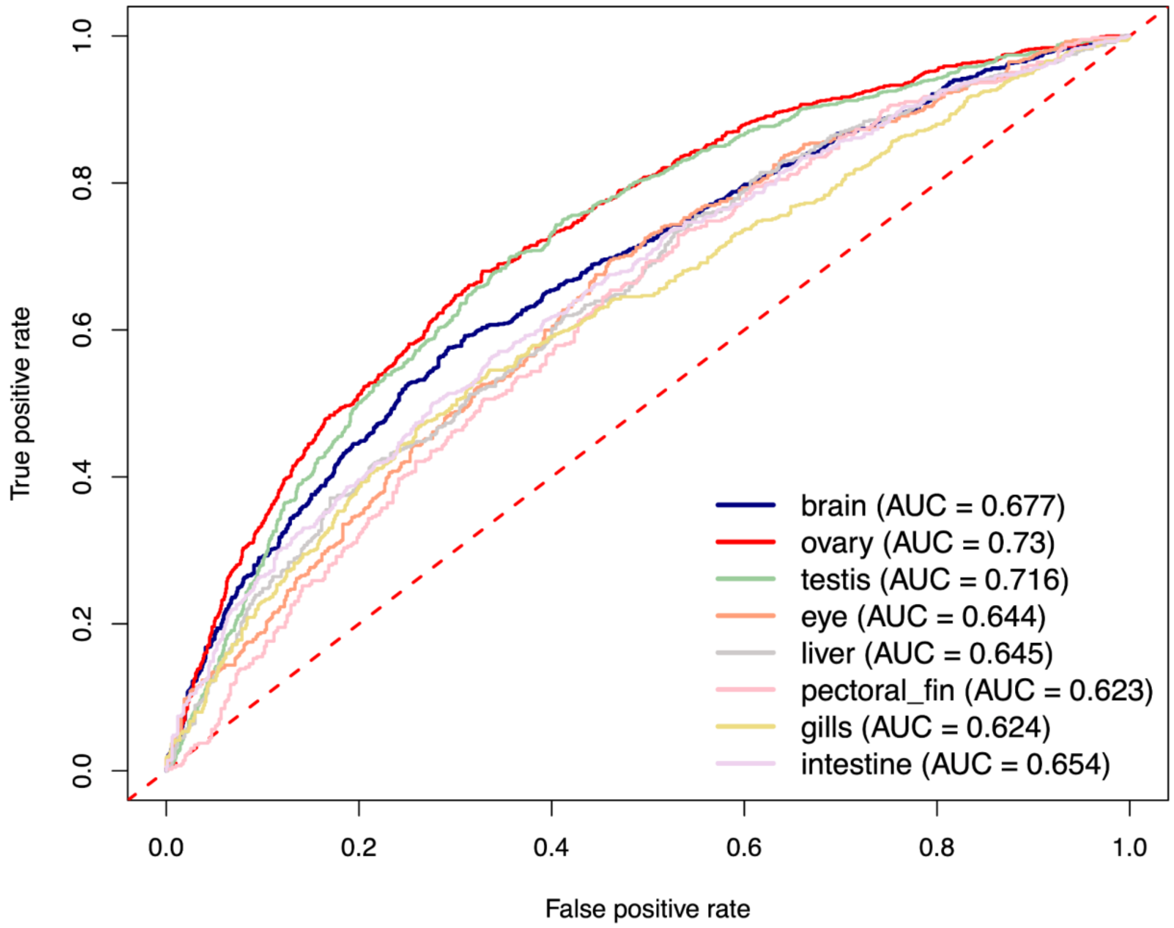


**Figure S62. ROC curves for gkmSVM classification performance on each organ-biased promoter set (N ≤ 1000 promoters) in zebrafish.** AUC values correspond to the area under the ROC curve from 5-fold cross-validation and provide an overall measure of predictive power. The dashed red line indicates an AUC of 0.5, corresponding to random performance. Each set of promoters was tested against random sequences from the DanRer11 genome assembly, matched for length, repeat content, and GC content.

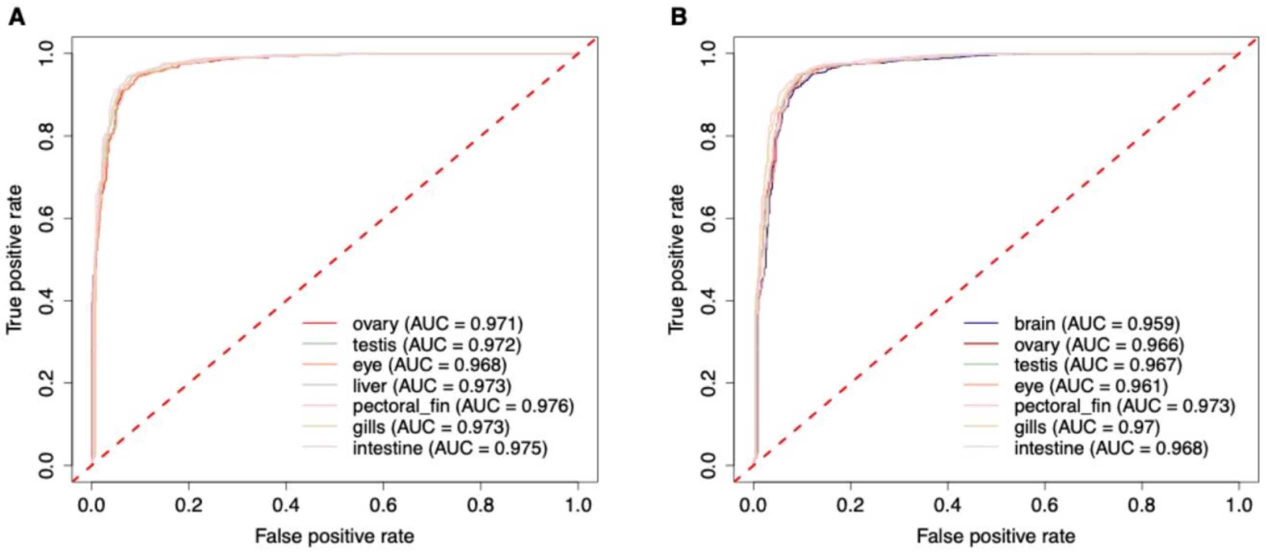


**Figure S63. ROC curves for the gkmSVM classification performance trained on the zebrafish (A) brain-biased promoter set and (B) liver-biased promoter set (N ≤ 1000 promoters), each tested against promoters from other organ-biased sets.** AUC values indicate the area under the ROC curve, reflecting the model's ability to distinguish the focal set from the others. The dashed red line indicates an AUC of 0.5, corresponding to random performance. Each set of promoters was tested against random sequences from the DanRer11 genome assembly, matched for length, repeat content, and GC content.

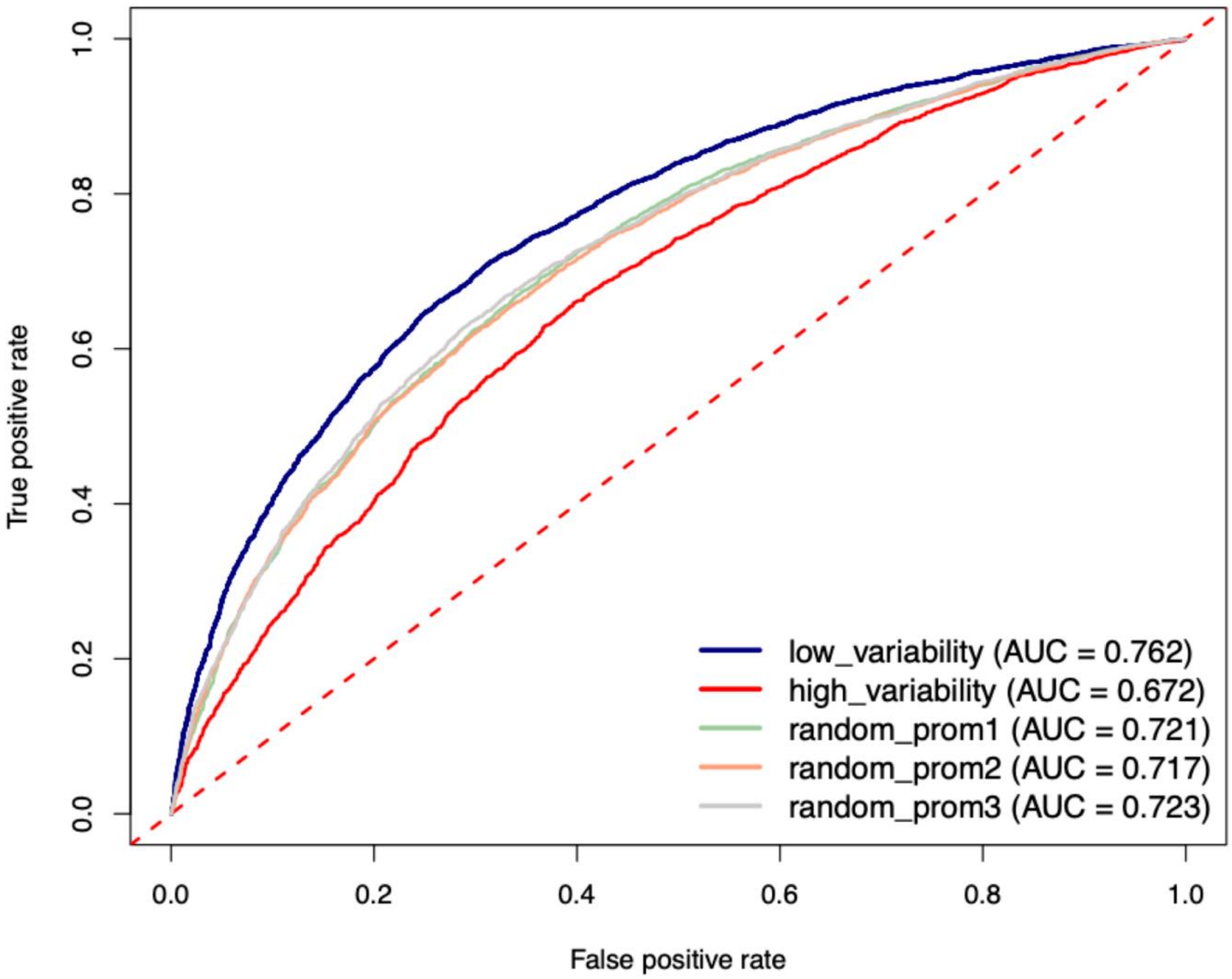


**Figure S64. ROC curves for gkmSVM classification performance on zebrafish promoters from two sets of organ-biased genes and three sets of randomly selected genes (N ≤ 5000 promoters).** The low variability set consists of zebrafish brain-, eye-, ovary- and testis-biased genes, while the high variability set consists of liver-, intestine-, pectoral fin-, and gills-biased genes. AUC values correspond to the area under the ROC curve from 5-fold cross-validation and provide an overall measure of predictive power. The dashed red line indicates an AUC of 0.5, corresponding to random performance. Each set of promoters was tested against random sequences from the DanRer11 genome assembly, matched for length, repeat content, and GC content.
